# Effects of copepod chemical cues on intra- and extracellular toxins in two species of *Dinophysis*

**DOI:** 10.1101/2024.07.17.603891

**Authors:** Milad Pourdanandeh, Véronique Séchet, Liliane Carpentier, Damien Réveillon, Fabienne Hervé, Clarisse Hubert, Philipp Hess, Erik Selander

## Abstract

Copepods may contribute to harmful algal bloom formation by selectively rejecting harmful cells. Additionally, copepods and the chemical cues they exude, copepodamides, have been shown to induce increased toxin production in paralytic and amnesic toxin producing microalgae. However, it is unknown if diarrhetic shellfish toxin (DST) producers such as *Dinophysis* respond to copepods or copepodamides in a similar fashion. Here we expose laboratory cultures of *Dinophysis sacculus* and *D. acuminata* to direct grazing by *Acartia* sp. copepods or copepodamides and measure their toxins after three days. Total *Dinophysis*- produced toxins (DPTs), okadaic acid, pectenotoxin-2, and C9-diol ester of okadaic acid, increased by 8 - 45% in *D. sacculus* but was significantly different from controls only in the highest (10 nM) copepodamide treatment whereas toxin content was not affected in *D. acuminata.* Growth rate was low across all groups and explained up to 91% of the variation in toxin content. DPTs were redistributed from internal compartments to the extracellular medium in the highest copepodamide treatments (5 - 10 nM), which were two to three times higher than controls and indicates an active release or passive leakage of toxins. Untargeted analysis of endometabolomes indicated significant changes in metabolite profiles for both species in response to the highest copepodamide treatments, independent of known toxins. However, it is not clear whether these are stress responses or caused by more complex mechanisms. The relatively small grazer-induced effect in *Dinophysis* observed here, compared to several species of *Alexandrium* and *Pseudo-nitzschia* reported previously, suggests that DPT production in *Dinophysis* is likely not induced by copepods, except perhaps in patches with high copepod densities. Thus, DPTs may, represent either a constitutive chemical defence for *Dinophysis*, or serve an altogether different purpose.

## 1. Introduction

*Dinophysis* is a highly specialised genus of non-constitutive mixotrophic dinoflagellates (Reguera et al., 2012; Mitra et al., 2016). They feed on ciliates of the genus *Mesodinium*, and retain the photosynthetically active chloroplasts (Hansen et al., 2013) that the ciliate, in turn, obtained from their own cryptomonad prey such as *Teleaulax* spp., *Plagioselmis* spp., and *Geminigera* spp. At least eight species of *Dinophysis* can be reared on a diet consisting of the ciliate *Mesodinium rubrum* (Park et al., 2006; Hansen et al., 2013; Séchet et al., 2021) which in turn are fed with cryptophytes. *Dinophysis* spp. grow at rates similar to other small dinoflagellates in fed cultures, but more slowly or not at all in the prolonged absence of prey (Reguera et al., 2012).

The link between diarrhetic shellfish toxins (DSTs: okadaic acid group toxins and dinophysistoxins (DTXs)) and the dinoflagellate *Dinophysis fortii* was established following multiple episodes of shellfish poisoning in Japan during the late 1970s (Yasumoto et al., 1980). The active compounds, identified as groups of polyethers, notably include okadaic acid (OA), DTXs, and their derivatives. They also produce pectenotoxins (PTXs), initially included as DSTs (Yasumoto et al., 1985; Ishige et al., 1988) but now concluded to be non-diarrhetic in humans (Terao et al., 1986; Matsushima et al., 2015). In sufficient amounts, the DSTs cause severe gastroenteritis in humans (Yasumoto et al., 1985) while pectenotoxins have been shown to negatively impact fertilisation success of oyster gametes (Gaillard et al., 2020),development of fish embryos (Gaillard et al., 2023), be hepatotoxic to some vertebrate hepatocytes (Terao et al., 1986; Fladmark et al., 1998), and are potently cytotoxic to several human cell lines (Jung et al., 1995).

*Dinophysis* is a cosmopolitan genus (Reguera et al., 2012) and the resulting syndrome, Diarrhetic Shellfish Poisoning (DSP), constitutes the largest share of harmful algal events in the North-East Atlantic (55%) reported in the Harmful Algae Event Database (Karlson et al., 2021). *Dinophysis* blooms are hard to foresee due to their unpredictable population dynamics and highly variable toxin production. The reason for their variable toxin content is not fully understood (Karlson et al., 2021) but is negatively correlated to cell density on the Swedish west coast. For example, Lindahl and colleagues (2007) reported okadaic acid content higher than 5 pg cell^-1^ in blooms with less than 100 cells L^-1^ but, with few exceptions, below 2 pg cell^-1^ in blooms with more than 250 cells L^-1^. Recent findings suggest that interactions with grazers may contribute to harmful algal bloom formation (Selander et al., 2019; Trapp et al., 2021). Chemical alarm cues from copepods, copepodamides (Selander et al., 2015), induce several-fold increases in cellular toxin content for paralytic and amnesic shellfish toxin (PSTs and ASTs, respectively) producers such as *Alexandrium* and *Pseudo-nitzschia*. (Selander et al., 2006; Tammilehto et al., 2015; Selander et al., 2019) and vary in composition between copepod taxa (Arnoldt et al., 2024). Moreover, copepods reject the more toxic cells and feed selectively on less defended cells (Xu and Kiørboe, 2018; Ryderheim et al., 2021; Olesen et al., 2022). As a result, the relative abundance of harmful cells will increase and to some extent contribute to harmful algal bloom formation (Xu et al., 2018). Recently, Trapp and colleagues (2021) tested this hypothesis by exploring the correlation between copepod biomass, or copepodamides in mussels, with the amount of DSTs in mussels on the Swedish west coast. They found that both copepod biomass and the concentration of copepodamides in blue mussels correlate to time-lagged DST content and can be used together with *Dinophysis* spp. cell densities to improve lead time and forecasting precision of DST concentrations in shellfish. It is, however, not known if copepods, or their chemical cues, induce the production of toxins in *Dinophysis*, or how their biosynthetic machinery responds to grazer presence.

Here, we address this knowledge gap by exposing laboratory cultures of *Dinophysis sacculus* and *D. acuminata* to direct grazing from *Acartia* sp. copepods, and to grazer-free copepodamide extracts. We measure growth rates, intra- and extracellular DPTs, and perform untargeted metabolomic analyses to determine the effect of copepod grazers on DPT production and metabolic state of *Dinophysis* spp. cells.

## 2. Materials and Methods

### 2.1 *Dinophysis* cultures

*Dinophysis sacculus* (IFR-DSa-01Lt, accession no. MT 37 1867) and *Dinophysis acuminata* (IFR-DAu-01Ey, accession no. MT 365104), isolated from Eyrac Pier, Arcachon, France in May 2015 and April 2020 respectively, were used. Both cultures were kept in diluted K (-Si)/ L1/20 (-Si) culture media (Guillard and Hargraves, 1993) prepared with filtered seawater (0.2 µm Steritop Corning, Corning, United States) at pH 8.2 and a salinity of 35. They were maintained in a thermo-regulated room at 17 °C and ∼55 µmol m^2^ s^-1^ photosynthetically active radiation on a 12h:12h light:dark cycle in 50 mL Erlenmeyer flasks. Irradiance was delivered by Osram Fluora 36W (Munich, Germany) and Philips Daylight 36W (Amsterdam, Netherlands). *Dinophysis* cultures were fed daily with 100 µL concentrated *Mesodinium rubrum* culture (DK-2009), isolated from Helsingør harbour (Denmark) in 2009, which in turn had been fed *Teleaulax amphioxeia* (strain AND-0710). Before the experiment, cultures were checked under a microscope to make sure that all *Mesodinium* cells had been consumed.

### 2.2 Copepodamides

The copepodamides used here were extracted and purified from commercially fished *Calanus finmarchicus*, as described by Selander and colleagues (2015). The copepodamide blend from *Calanus* sp. (Fig. 1A) is more complex than that of the *Acartia* sp. copepods used in experiments but also includes the copepodamides found in *Acartia* sp. (Fig. 1B, Supplementary Table S1).

**Fig. 1:**
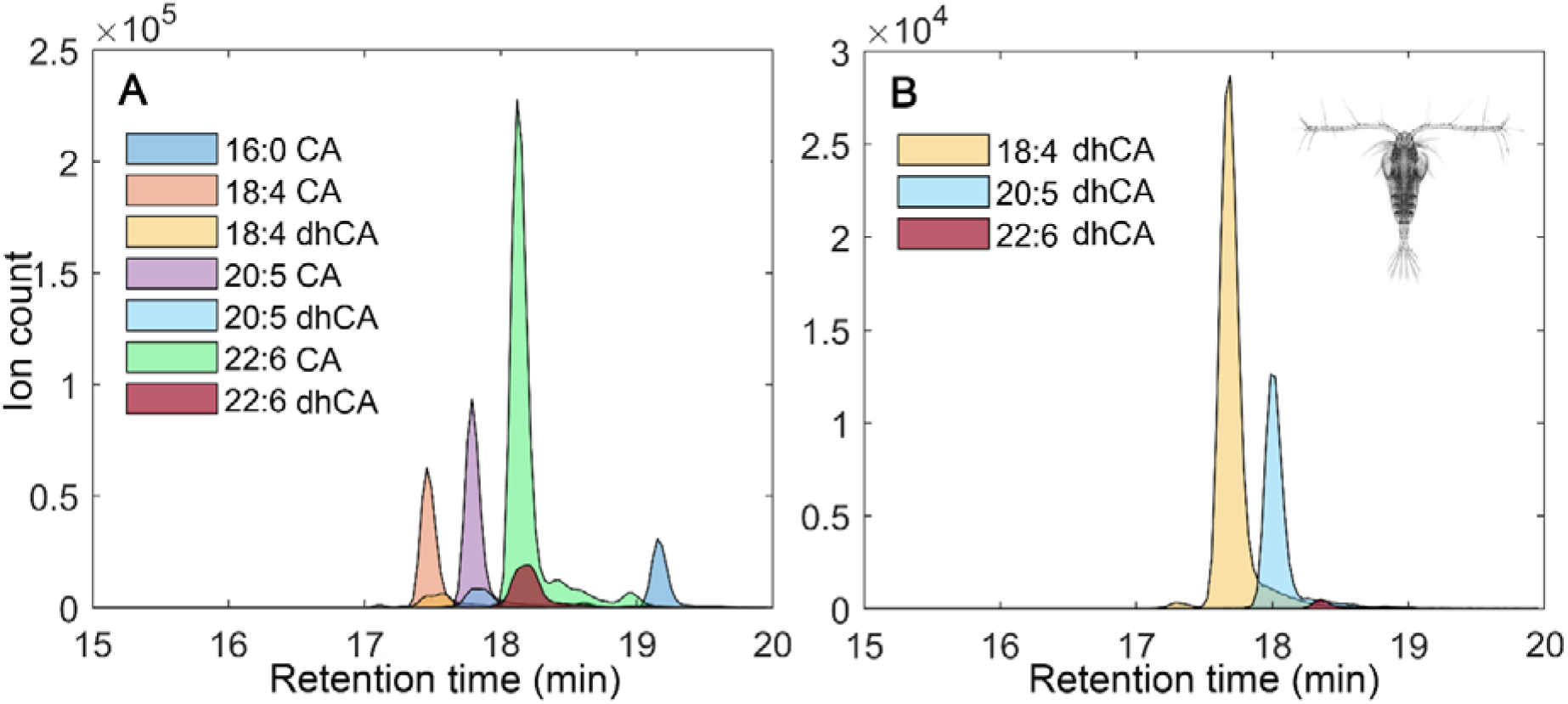
Chromatogram of (**A)** the *Calanus finmarchicus* derived and purified copepodamide mixture used in experiments and (**B**) copepodamides found in the *Acartia* sp. copepods used as live grazers in this induction experiment. The trace amounts of 14:0 CA found in the purified mixture were too low to be observed in this figure.

### 2.3 Experimental procedure

Cultures of *D. sacculus* and *D. acuminata*, with cell concentrations of 1728 ± 62 cells mL^-1^ and 1254 ± 76 cells mL^-1^ (mean ± SEM) respectively, were divided between fifteen 20 mL Erlenmeyer flasks. Nine flasks for each species were prepared by adding copepodamides dissolved in methanol (10 µL) to achieve nominal copepodamide concentrations of 1, 5, or 10 nM, each in triplicate. Triplicate controls for each species were prepared in the same way, but with 10 µL methanol without copepodamides. The methanol was allowed to evaporate before adding the *Dinophysis* cultures, leaving the copepodamides coated onto the culture flasks walls. The resulting effective concentration in the culture media, due to a combination of dissolution and degradation, typically averages between 0.1% and 1.5% of the nominal concentration (Selander et al., 2019). To maintain copepodamide stimuli throughout the experimental period, the experimental cultures were transferred to freshly coated Erlenmeyer flasks after 48 hours. The experiment was concluded after three days (68.5 h).

To control for the possibility that *Dinophysis* respond to other copepod cues than copepodamides, we included a living copepod treatment as a positive control. The copepods (*Acartia* sp.) were collected with a plankton net at Le Croisic (WGS84 decimal degrees: 47.299796, -2.517140) the day before the experimental start and individually isolated with a Pasteur pipette under a dissecting microscope into a multi-well dish. The water was carefully replaced by medium three times to avoid contaminating organisms or chemical stimuli from the net haul. Finally, a single copepod was added to three flasks (resulting in 50 ind L^-1^) for both *Dinophysis* species together with a small volume of medium (<100 µL). The copepods were inspected daily, and non-motile copepods were replaced. In total, two copepods, one in each *Dinophysis* species, were replaced. All experimental units received 100 µL *Mesodinium* on day 0, 1, and 2 (85, 53, and 89 ×10^3^ cells mL^-1^ respectively) to avoid starvation of the *Dinophysis* cells.

### 2.4 Toxin and metabolomic samples

Triplicate samples were taken from all treatments of both species at the start and end of the experiment. Well-mixed samples of 15 mL were centrifuged (4300g, 20 min, 4 °C) and supernatants carefully pipetted into a 50 mL centrifuge tube. The remaining pellets and supernatants were stored frozen at -80 °C until extraction.

### 2.5 Growth rate

At the end of the experiment, a well-mixed sample (1.5 mL) from each replicate was removed for cell enumeration. The sample was diluted to 15 mL and preserved with acid Lugol’s solution. Aliquots of 1 mL were counted under an inverted microscope until >500 cells per replicate had been counted. Specific growth rate (µ) was calculated as:

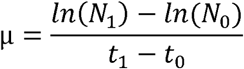

where N_0_ and N_1_ are cell concentrations at the start (t_0_) and end (t_1_) of the experiment (Frost, 1972).

### 2.6 Copepodamide analysis

Five adult *Acartia* copepods, possibly including some non-conspicuous late-stage copepodites (C5), were pipetted into an HPLC vial; the water removed with a Pasteur pipette, and extracted in 1.5 mL methanol for at least 24 h. The methanol extract was transferred to a glass tube and evaporated under a stream of nitrogen gas at 40 ^◦^C, resuspended in 50 µL methanol and transferred to a low volume HPLC-vial insert. Copepodamides were analysed on an Agilent 1200 series HPLC fitted with a 6470 triple quadrupole mass spectrometer (Santa Clara, CA, USA). A multiple reaction monitoring experiment targeting the 10 known copepodamides was performed to quantify both the copepodamide extract from *Calanus* sp. used in experiments and the extract from *Acartia* sp. Grebner et al. (2019), for further details see Selander et al. (2019).

### 2.7 Toxin analyses

Okadaic acid group toxins and pectenotoxins were extracted following Séchet and colleagues (2021). Briefly, intracellular toxins were extracted from cell pellets twice with 0.75 mL methanol, vortexed, and sonicated (25 kHz for 10 min). Samples were then centrifuged for 5 min (4300 g, 4 °C), the extracts were pooled, filtered (0.2 µm Nanosep MF filter, Pall), and stored at -20 °C until analysis with liquid chromatography coupled to tandem mass spectrometry (LC-MS/MS). Extracellular toxins were recovered from the supernatant after two liquid-liquid partitioning procedures with 25 mL dichloromethane. After evaporation to dryness, and resuspension in 1 mL methanol, the extracts were filtered and stored at -20 °C until LC-MS/MS analysis.

Toxins were separated on an a ultra-fast liquid chromatography system (Prominence UFLC-XR, Shimadzu, Japan), equipped with a reversed-phase C18 Kinetex column (100 Å, 2.6 μm, 100 × 2.1 mm, Phenomenex, Le Pecq, France) and coupled to a triple quadrupole/ion trap mass spectrometer (4000QTrap, Sciex, Framingham, MA, USA), as described by Séchet and colleagues (2021). Authentic standards (OA, DTX-1, DTX-2, and PTX-2) for quantification were obtained from the National Research Council of Canada (Halifax, Nova Scotia, Canada) and the uncertified C9-diol ester of OA (C9) from Cifga (through Novakits, Nantes, France).

### 2.8 Metabolomics

Metabolomic profiles of the extracts prepared for intracellular toxin analysis above were acquired by liquid chromatography coupled to high resolution mass spectrometry (LC-HRMS) as in Gémin et al. (2021). Briefly, an UHPLC (1290 Infinity II, Agilent technologies, Santa Clara, CA, USA) coupled to a quadrupole time-of-flight mass spectrometer (QTOF 6550, Agilent technologies, Santa Clara, CA, USA) equipped with a Dual Jet Stream ESI interface was used. Both positive (+MS) and negative (-MS) full scan modes were used over a mass-to-charge ratio (*m/z*) ranging from 100 to 1700. The injection volume was 5 µL. The batch was prepared as recommended by Broadhurst and colleagues (2018), by using a mix of four phycotoxins (OA, DTX-1, DTX2, PTX-2) as the standard reference material and a mixture of all samples for the qualitative control (QC) samples. The injection order was randomised, and QCs were injected every 5 samples.

LC-HRMS raw data was converted to mzXML-format using MS-Convert in ProteoWizard 3.0 (Chambers et al., 2012) and pre-processed with the Workflow4Metabolomics 4.0 e-infrastructure (Guitton et al., 2017). Peak picking, grouping, retention time correction, and peak filling were performed with its “CentWave”, “PeakDensity”, “PeakGroups”, and “FillchromPeaks” algorithms. Annotation (e.g. isotopes and adducts) was conducted with the “CAMERA” algorithm (Kuhl et al., 2012). Intra-batch signal intensity drift was corrected for, in the –MS data only, by fitting a locally quadratic (LOESS) regression model to the QC values (van der Kloet et al., 2009; Dunn et al., 2011). Three successive filtering steps based on signal-to-noise ratio with blanks, variability among pools, and autocorrelation were applied using in-house scripts in R, as described by Georges des Aulnois et al. (2020). Pre-processing of +MS data matrices for each species led to 19427/20570 features for *D. acuminata* and *D. sacculus*, respectively, while -MS data matrices led to 4406/4816. After filtrations 1788/695 and 422/324 features remained, respectively.

The data matrix in +MS and -MS for each species was log transformed and Pareto scaled, and a weighting factor based on the maximal cell number was applied for each sample. Unsupervised principal component analysis (PCA) was used for data exploration, while features significantly affected by copepodamides or copepods were selected by ANOVA followed by a Tukey’s post hoc test, with adjusted p-value cut-offs set to 0.001 or 0.01 depending on the species and mode of ionisation. Statistical analyses of metabolomics were performed with MetaboAnalyst 5.0 (Pang et al., 2021)

In parallel, HRMS data were acquired by autoMS/MS as in Georges des Aulnois et al. (2019) to attempt annotation of the significant features revealed by metabolomics, using both GNPS (Wang et al., 2016) and SIRIUS (Dührkop et al., 2019). The comparison of MS/MS spectra with these reference databases provided a level 2 of annotation, as defined by Schymanski and colleagues (2014). In addition, diagnostic ions in MS/MS spectra were considered, especially for the (lyso)diacylglycerylcarboxyhydroxymethylcholine (lysoDGCC) and (lyso)phosphatidylcholine (lysoPC) families (Roach et al., 2021; Rey et al., 2022). When the fatty acyl chain could not be confirmed, the features were considered “lysoDGCC-like” or “lysoPC-like” (.xlsx file; Supplementary spreadsheet 1).

### 2.9 Statistical analyses of growth rates and toxins

For analyses of toxin induction, the three toxin congeners (OA, PTX-2, C9) were pooled (referred to as total toxins or total DPTs). Differences in specific growth rate (µ) and intracellular toxins (% of total toxins) as a function of species (two levels: *D. sacculus* & *D. acuminata*) and treatment (five levels: Control, Copepod, 1 nM, 5 nM & 10 nM) were tested with two-way analyses of variance (ANOVA). Volume normalised total (intracellular and extracellular) DPT content (ng mL^-1^) was first analysed with a two-way analysis of covariance (ANCOVA) with growth rate as covariate. Because there was no interaction effect between treatment and species, but a significant difference between the species, they were analysed with separate one-way ANCOVAs instead. The proportions (%) of individual toxin congeners (OA, PTX-2, and C9) in *D. sacculus* were tested with separate one-way ANOVAs. All proportions were logit (Ln(p/1-p)) transformed for better statistical properties (Warton and Hui, 2011).

Assumptions of linearity were evaluated visually and with linear regressions. Normality was visually inspected and formally tested with Shapiro-Wilk’s test, and homogeneity of variances was tested with Levene’s test. Significant ANOVAs (p ≤ 0.05) were followed up with Tukey’s honest significant difference (HSD) post-hoc procedure. Estimates of covariate corrected mean toxin content, and their 95% confidence intervals (CIs hereafter), were extracted from the linear models using the base R *predict* function at covariate level (i.e. growth rate) set to zero. Unless otherwise stated, all summary statistics values are given as mean (95% CI: lower limit - upper limit) or (mean, lower CI limit - upper CI limit). Effect sizes are derived from log response ratios (LRR) and presented as the percentage difference (lower 95% CI limit – upper CI confidence) or as (mean, lower CI limit–upper CI limit) (Hedges et al., 1999). Partial eta-squared (η^2^_p_) was used as effect size for linear models.

Statistical analyses of growth rates and toxins, and visualisations of growth rate, toxins, and PCA scores derived from metabolomics data were performed in R v.4.4.1 (R Core Team, 2024) using RStudio v.2024.4.1.748 (Posit team, 2024). The code (.html file; Supplementary code & output) and the toxin analysis data it uses (.csv file; Supplementary spreadsheet 2) are available as supplementary materials. These, Supplementary spreadsheet 1, and the HRMS metabolomics data are also openly accessible from a public Zenodo repository (Pourdanandeh et al., 2024). Supplementary tables and figures directly referred to in the text is found in Supplementary information

## 3. Results

### 3.1 Copepodamides

The *Acartia* sp. copepods used as grazers in this experiment contained approximately ∼0.5 pmol ind^-1^ dihydro-copepodamides (hereafter dhCAs). The most abundant compound was 18:4-dhCA, followed by 20:5-dhCA and 22:6-dhCA, along with trace amounts of the deacylated molecular scaffold (Fig. 1B). The copepodamides extracted from *Calanus finmarchicus* used in the experiment included all three dhCAs found in *Acartia* sp. as minor components along with other copepodamides (Fig. 1A, Table S1).

### 3.2 Growth rate

Average growth rates were low and ranged from -0.10 to 0.05 day^-1^ (Fig. 2) but did not differ between treatments (p = 0.22, Table 1) or the two species of *Dinophysis* (p = 0.94). However, growth rate was positively correlated with total DSTs in both *D. sacculus* (p < 0.001) and *D. acuminata* (p = 0.025) and accounted for 91% and 44% of the variance left unexplained by treatment, respectively.

**Fig. 2:**
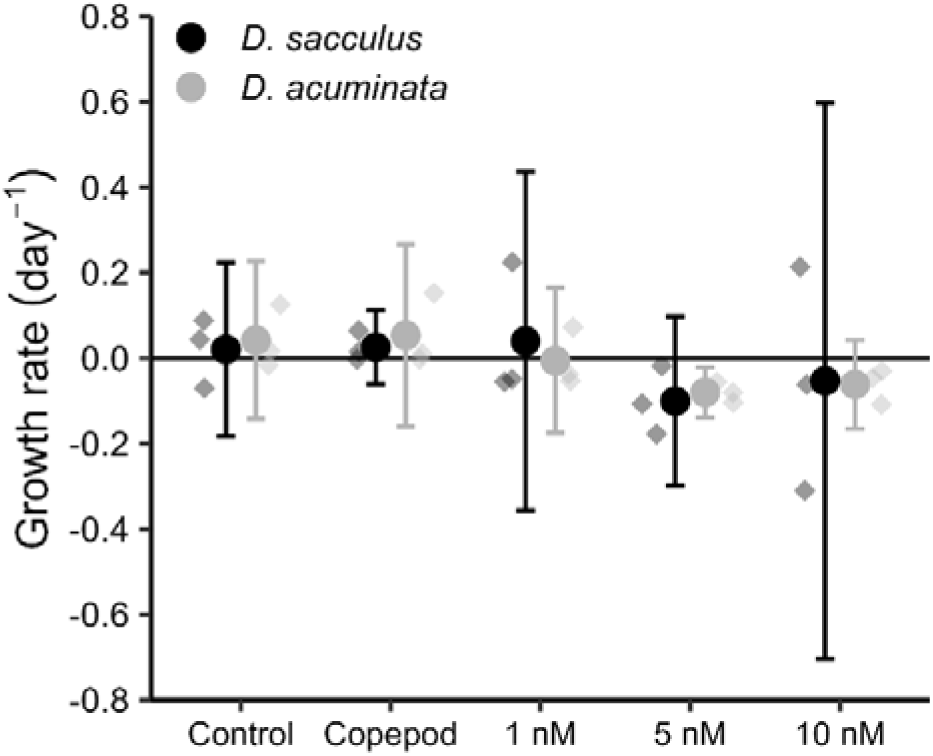
Growth rate of *Dinophysis sacculus* (black) and *Dinophysis acuminata* (grey) after 68.5 hours of exposure to 1-10 nM concentrations of copepodamides, a living *Acartia* sp. copepod, or control conditions without copepods or copepod cues. Transparent diamonds represent individual growth rates. Circles denote mean growth rates (n = 3) and error bars show their 95% confidence intervals.

**Table 1.** Results from linear models analysing growth rate (model I), total *Dinophysis* produced toxins (DPTs; models II-IV), proportion of toxins inside vs. outside of cells (model V), and individual toxin congener proportions of the total DPTs (models VI-VIII). Bold values indicate statistically significant (p ≤ 0.05) model terms, IV = independent variable, Cv = covariate.

| Model type | Model number | Response variable | Term | SS | MS | df | F | p | Partial $\eta^2$ |
| --- | --- | --- | --- | --- | --- | --- | --- | --- | --- |
| Two-way ANOVA | I | Growth rate | Treatment (IV) | 0.082 | 0.0205 | 4 | 1.59 | 0.22 | 0.241 |
|  |  |  | Species (IV) | 7.2E-05 | 7.2E-05 | 1 | 0.006 | 0.94 | 3E-04 |
|  |  |  | Treatment * Species | 0.006 | 0.0015 | 4 | 0.11 | 0.98 | 0.022 |
|  |  |  | Residuals | 0.26 | 0.0128 | 20 |  |  |  |
| Two-way ANCOVA | II | Total toxin content | Growth rate (Cv) | 24738 | 24738 | 1 | 95.6 | < **0.001** | 0.834 |
|  |  |  | Treatment (IV) | 7218 | 1805 | 4 | 6.97 | **0.00** | 0.595 |
|  |  |  | Species (IV) | 9446 | 9446 | 1 | 36.50 | < **0.001** | 0.658 |
|  |  |  | Treatment * Species | 1047 | 262 | 4 | 1.01 | 0.43 | 0.176 |
|  |  |  | Residuals | 4918 | 259 | 19 |  |  |  |
| One-way ANCOVA | III | Total toxin content of *D. sacculus* | Growth rate (Cv) | 23883.0 | 23883 | 1 | 89.3 | < **0.001** | 0.91 |
|  |  |  | Treatment (IV) | 6754 | 1689 | 4 | 6.3 | **0.011** | 0.74 |
|  |  |  | Residuals | 2406 | 267 | 9 |  |  |  |
| One-way ANCOVA | IV | Total toxin content of *D. acuminata* | Growth rate (Cv) | 1495 | 1495 | 1 | 7.19 | **0.025** | 0.444 |
|  |  |  | Treatment (IV) | 1060 | 265 | 4 | 1.28 | 0.35 | 0.362 |
|  |  |  | Residuals | 1871 | 208 | 9 |  |  |  |
| Two-way ANOVA | V | Intracellular toxin proportion | Treatment (IV) | 29.9 | 7.48 | 4 | 92 | < **0.001** | 0.949 |
|  |  |  | Species (IV) | 37.4 | 37.45 | 1 | 461 | < **0.001** | 0.958 |
|  |  |  | Treatment * Species | 1.38 | 0.345 | 4 | 4.24 | **0.012** | 0.459 |
|  |  |  | Residuals | 1.62 | 0.081 | 20 |  |  |  |
| One-way ANOVA | VI | OA proportion | Treatment (IV) | 0.034 | 0.0085 | 4 | 3.195 | 0.062 | 0.561 |
|  |  |  |  | 0.026 | 0.0026 | 10 |  |  |  |
| One-way ANOVA | VII | PTX-2 proportion | Treatment (IV) | 0.01 | 0.0030 | 4 | 0.73 | 0.59 | 0.227 |
|  |  |  |  | 0.04 | 0.0042 | 10 |  |  |  |
| One-way ANOVA | VIII | C9 proportion | Treatment (IV) | 0.17 | 0.0413 | 4 | 0.855 | 0.162 | 0.255 |
|  |  |  |  | 0.48 | 0.0484 | 10 |  |  |  |

### 3.3 Toxins

Although total *Dinophysis*-produced toxins (DPTs) in *D. sacculus* generally increased (8 - 45%) with higher copepodamide concentrations and exposure to live copepods, only the highest copepodamide treatment was significantly different from control conditions (p = 0.008, Fig. 3. Table 1). *D. sacculus* cells produced 45.1% (20.3 - 74.6) more DPTs in the 10 nM copepodamide treatment than in controls and 34% (12.2 - 50.4) more than in the live copepod treatment (p = 0.027). *D. acuminata* okadaic acid (OA)content displayed a similar trend, with 18.5 - 20.5% increases in all treatment groups compared to controls which, however, were not significant (p = 0.35, Table 1). N.B. raw means and confidence intervals, i.e. without correcting for growth rates, is available in Fig. S1.

**Fig. 3:**
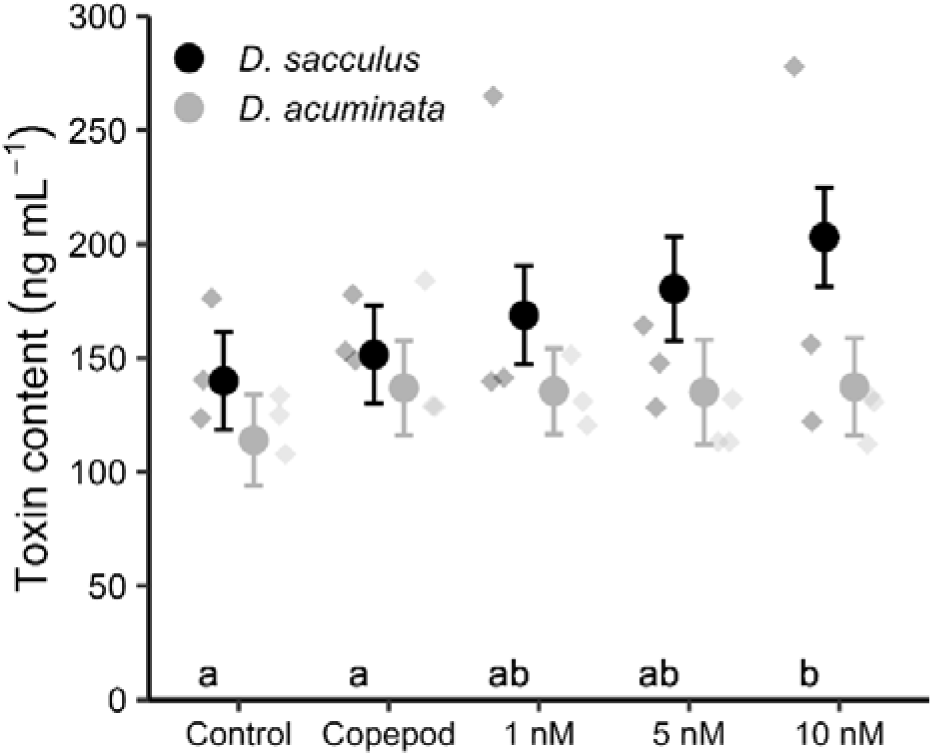
Total *Dinophysis* produced toxins, normalised by sample volume, for *Dinophysis sacculus* (black) and *Dinophysis acuminata* (grey) after 68.5 hours of exposure to 1-10 nM concentrations of copepodamides, a living *Acartia* sp. copepod, or control conditions without copepods or copepod cues. Circles represent the covariate (growth rate) adjusted mean values (n = 3), predicted at growth rate = 0, and error bars denote the 95% confidence intervals of the predicted means. Transparent diamonds show the measured toxin content in each replicate (i.e., not covariate adjusted). Letters denote statistically homogeneous subgroups for *D sacculus.* There were no significantly different subgroups in *D acuminata*.

Toxins were redistributed from the inside of cells to their surroundings in both species (Fig. 4), and the effect was most pronounced for *D. acuminata* where the extracellular fraction doubled from 46 - 54% to 90 - 94% (p < 0.001, Table 1). Intracellular toxins in the 5 and 10 nM copepodamide treatments were significantly lower than in all other treatments for both species (Table. 1), ranging from 53 to 72% and 13 to 21% of control levels for *D. sacculus* and *D. acuminata*, respectively.

**Fig. 4:**
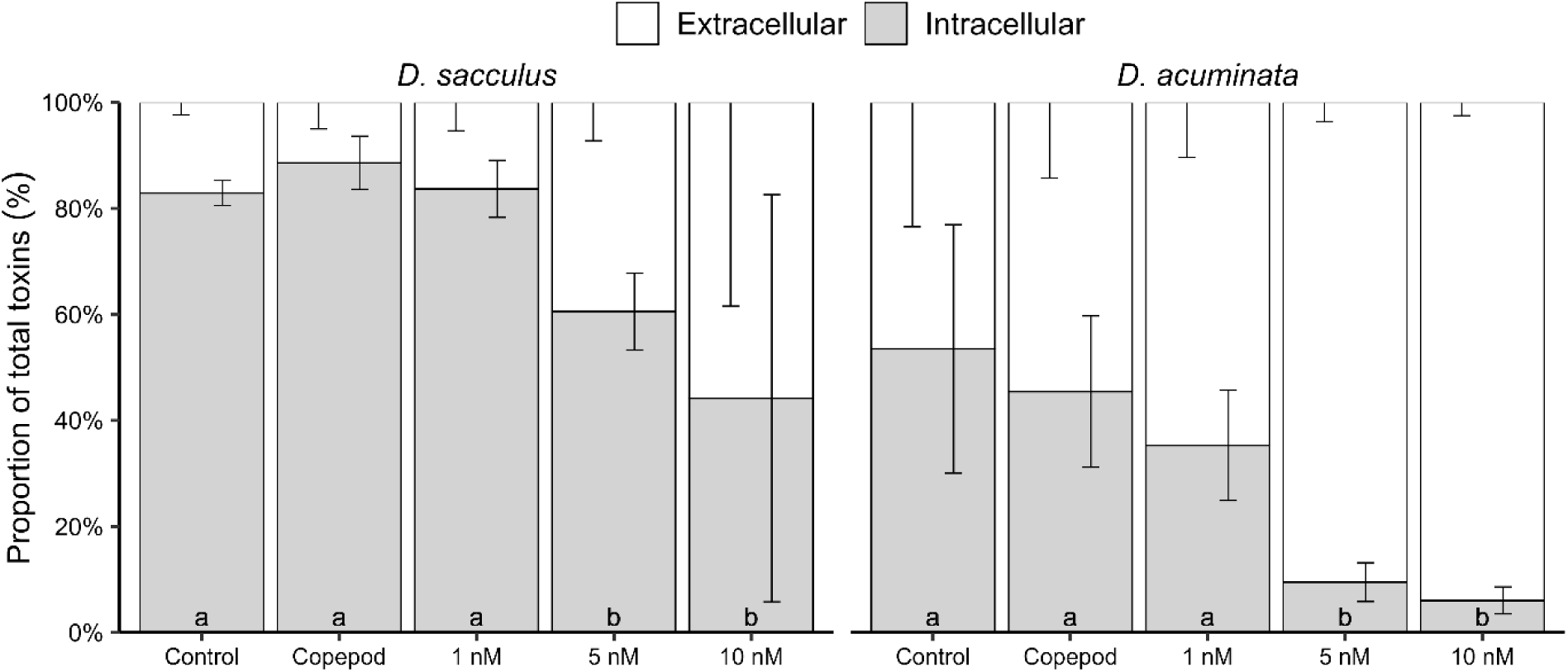
Proportion of intracellular (grey) and extracellular (white) toxins for *Dinophysis sacculus* (left) and *Dinophysis acuminata* (right) after 68.5 hours of exposure to 1-10 nM concentrations of copepodamides, a living *Acartia* sp. copepod, or control conditions without copepods or copepod cues. Bars represent the mean value of three replicates (n = 3) and error bars show the 95% confidence intervals of the means. Letters denote statistically homogeneous treatments. Note that only the lower confidence limits are shown for extracellular bars.

PTX-2 accounted for almost all the increase in total DPTs produced by *D. sacculus* (Fig. S2), but the proportions of individual toxin congeners (OA, PTX-2, C9) did not significantly differ between any treatment groups (p-values: 0.062, 0.6, 0.16; Table 1). On average, these congeners accounted for 4.7, 94, and 1.3% of total toxins, respectively (Fig. 5). *D. acuminata* only produced OA.

**Fig. 5:**
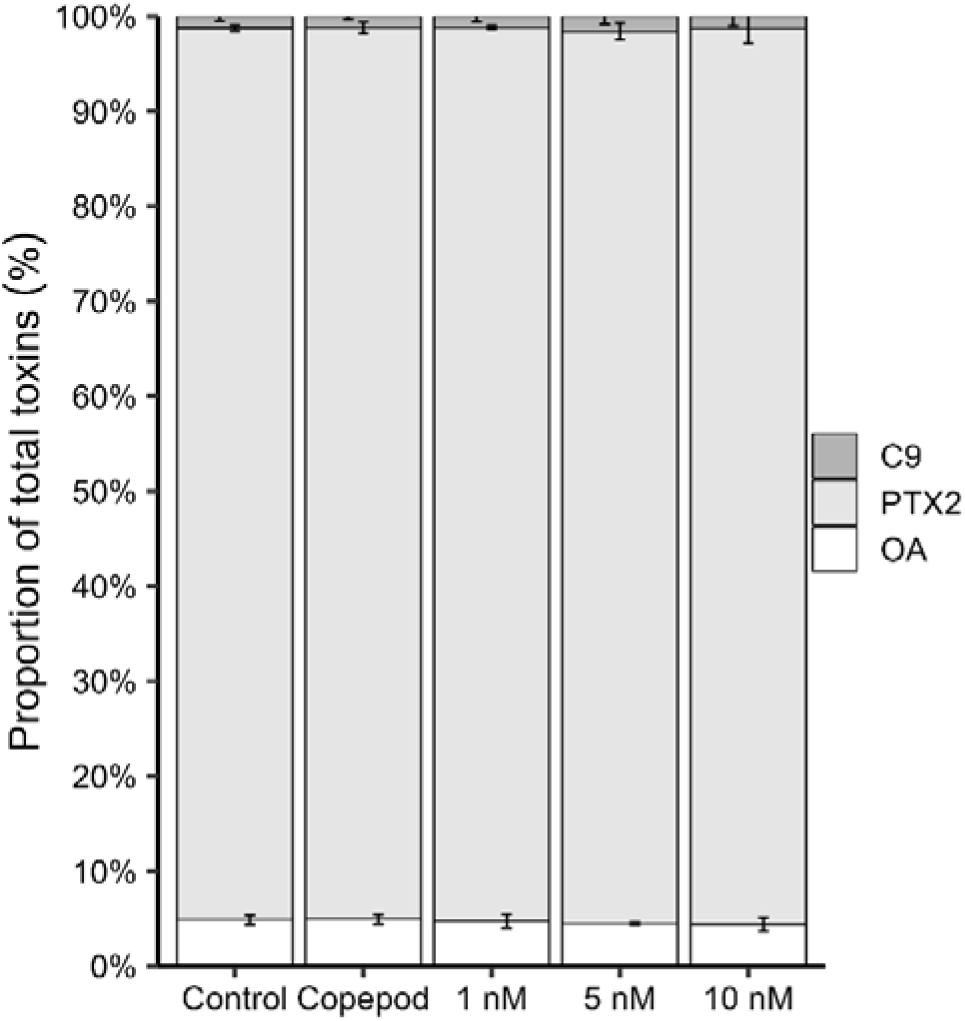
Composition of *Dinophysis* produced toxins (white = OA (Okadaic acid), light grey = PTX-2 (Pectenotoxin-2) and dark grey = C9 (OA-C9-diol ester) for *Dinophysis sacculus* (*D. acuminata* produced OA only) after 68.5 hours of exposure to 1-10 nM concentrations of copepodamides, a living *Acartia* sp. copepod or control conditions without copepods or copepod cues. Bars are the mean values of three replicates (n = 3) and error bars denote the 95% confidence intervals of the means. Note that only the lower confidence limits are shown for C9.

### 3.4 Metabolomics

Metabolomic profiles of *Dinophysis* cells exposed to control conditions, live copepods, or 1 nM copepodamides formed clear clusters for both species (Fig. 6, Fig. S3). Meanwhile, the 5 and 10 nM treatments diverged from this cluster and, to a lesser extent, from each other. This suggests a dose-dependent effect of copepodamides on the metabolomes of both *D. sacculus* and *D. acuminata.* Indeed, 55 and 15 features (in +MS and -MS respectively) differed significantly between treatments for *D. sacculus*, and 28 and 11 were significantly different for *D*. *acuminata* (Supplementary spreadsheet 3).

**Fig. 6:**
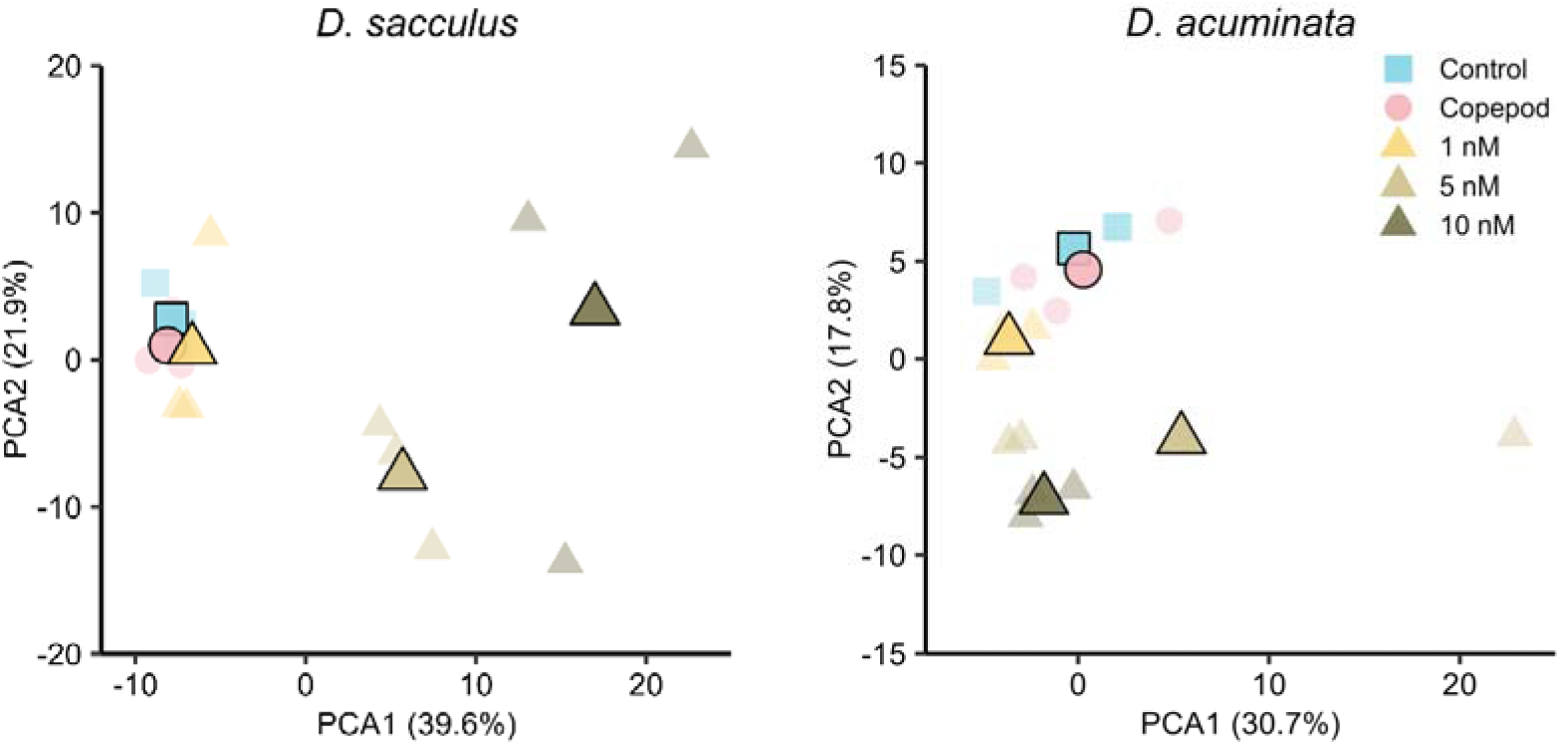
PCA score plot from LC-HRMS derived metabolomic profiles for *D. sacculus* (left, +MS data) and *D. acuminata* (right, -MS data) after 68.5 hours of exposure to three concentrations of chemical cues purified from copepods (copepodamides, triangles), a living *Acartia* sp. copepod (circles), or control conditions (squares). Transparent symbols without borders are individual replicate dimensionality reductions, and opaque symbols with black border denote the centroids (averages) for each group.

Overall, two expression profiles were observed for the two *Dinophysis* species: 24/11 and 10/2 features (+MS/-MS) were co-upregulated with increasing copepodamide concentration, while 30/4 and 18/9 features were co-downregulated in D. *sacculus* and D. *acuminata*, respectively (Figs. S4-S7). One peculiar over-expressed feature was also noted in the copepod treatment for *D. sacculus* only (M298T600, +MS). Proportionally, more features were co-downregulated with copepodamide concentration for *D. acuminata* than for *D. sacculus*.

## 4. Discussion

### 4.1 Toxin induction

We present the first experimental study that, to our knowledge, assesses the effects of copepod grazer cues (copepodamides) on *Dinophysis* toxin production and their metabolomes. Total *Dinophysis* produced toxins (DPTs) increased. slightly in *D. sacculus* cells that were exposed to grazing *Acartia* sp. copepods or copepodamides (Fig. 3) and displayed a positive trend as copepodamide concentration increased. However, this increase, ranging from 8 to 45%, was significantly different from non-induced cells only in the highest copepodamide treatment (10 nM). The effective copepodamide concentration in this treatment is one to two orders of magnitude higher than the average concentrations expected in natural conditions (Selander et al., 2019), but possibly on par with those experienced near copepods or in patches with high (>50 ind L^-1^) copepod densities (Lindström et al., 2017). In addition, *D. acuminata* toxins did not differ significantly from non-exposed cells. It’s therefore possible that the variation in cellular toxin content observed in *Dinophysis* cells in the field (Lindahl et al., 2007) are related to changes in strain composition as blooms develop, because variability in toxin content between strains of the same species or clade may be up to ten-fold (Séchet et al., 2021). In contrast to the present study, paralytic and amnesic shellfish toxins producers often respond to copepods and copepodamides by several fold increases in toxin production over similar time frames (Bergkvist et al., 2008; Lundholm et al., 2018; Griffin et al., 2019; Selander et al., 2019; Ryderheim et al., 2021). Hence, it seems unlikely that grazer-induced DPT production contributes significantly to the total of toxin content in natural populations of *Dinophysis* spp., at least based on the conditions and time frame covered here. It should, however, be noted that growth rate was low to absent in our experiment (Fig. 2), which may have constrained toxin production. Toxin production in *Alexandrium catenella* (previously *A. fundyense*) mainly takes place directly after division, in the G1 phase of the cell cycle (Taroncher-Oldenburg et al., 1997) and with limited growth few cells would have passed the G1 phase during our exposure period. Regardless, the modest induction of DPTs here raises question about the primary purpose of *Dinophysis* toxins. Do they serve as a constitutive defence trait? Do they function as deterrents for specific grazers not tested here? Are they used to immobilise prey? Or do they have a different role altogether?

Grazer induced toxin production indicates that the toxin functions as a defence against that grazer (Stahl, 1888), especially when the induction correlates to increase resistance to grazers (Ryderheim et al., 2021; Olesen et al., 2022). Both inducible and constitutional chemical defences are common in plant secondary metabolism (Stamp, 2003). Inducible defences are favoured when there is a reliable (i.e., source specific and non-persistent) cue associated with the level of threat, the threat level is variable, and the defence metabolites are costly (Karban and Baldwin, 1997; Tollrian and Harvell, 1999). Copepodamides are a reliable cue as they correlate to copepod density and rapidly degrade in seawater (Selander et al., 2019). Daily vertical migration by both copepods and *Dinophysis*, combined with differences in seasonal population dynamics of both predator and prey, indeed results in a variable threat level across both time and space, which should favour the evolution of inducible defences (Bergkvist et al., 2008; Ryderheim et al., 2021; Rigby et al., 2022). However, the costs of producing DPTs are unknown. Unlike the nitrogen-rich PSTs and AST alkaloids, DSTs and PTXs are cyclic polyethers and contain little to no nitrogen, potentially making them less costly to produce in the often nitrogen-limited marine ecosystem (Bristow et al., 2017). If metabolically cheap, there is less selective pressure on *Dinophysis* to modulate toxin production according to grazing pressure and a high constitutive chemical defence could instead be adaptive (Karban et al., 1997; Chen, 2008). This is supported by the plant apparency hypothesis (Feeny, 1975, 1976; Rhoades and Cates, 1976), which predicts that large and/or more apparent (e.g., highly motile) organisms have a higher degree of constitutive defence, unless this decreases their overall fitness (Harvell, 1990; Stamp, 2003). If DPTs function as chemical grazer deterrents it is possible that *Dinophysis* could employ a combination of both strategies, permanently producing toxins as a constitutive defence but able to increase this production when induced by increased grazing risk. PTX-2 was the main contributor to the increase in total DPTs produced by *D. sacculus* (Fig. S2) here. It could be that PTXs are used by *Dinophysis* as copepod specific deterrents, whereas DSTs lack any such function. Despite likely being non-diarrhetic to humans and differing from DSTs in both structure and effect, it remains a relevant inclusion as PTXs has demonstrated toxicity to some invertebrates (Pease et al., 2022) and recently suggested to reduce survival of the copepod *Acartia tonsa* (Ladds et al., 2024). Additionally, PTX-2 has shown toxicity to several vertebrate (Terao et al., 1986; Fladmark et al., 1998) and human cells (Jung et al., 1995), underscoring the potential ecological and health impacts of its production and introduction to food webs.

It is also possible that the copepods used here do not represent the main grazers of *Dinophysis* and that DPTs evolved to deter other grazers. Although many copepods co-exist with *Dinophysis* and are known to feed on them in both field and experimental settings (Carlsson et al., 1995; Frangoulis et al., 2022), the extent of this is unknown. Furthermore, *Acartia* spp. and several other copepod species (e.g., *Euterpina acutifrons*) avoid feeding on *D. acuminata* at bloom densities (Turner and Anderson, 1983; Maneiro et al., 2000). Copepods are estimated to consume between 10 and 25% of pelagic primary production (Calbet, 2001), whereas microzooplankton such as ciliates, heterotrophic flagellates. and rotifers consume an estimated 67% (Calbet and Landry, 2004). Microzooplankton grazers generally target prey as small or smaller than themselves (Hansen et al., 1994), and several species of mixotrophic and heterotrophic dinoflagellates such as *Alexandrium ostenfeldii* (Jacobson and Anderson, 1996), *Noctiluca scintillans* (Escalera et al., 2007), and *Fragilidium* spp. (Jacobsen, 1999; Nézan and Chomérat, 2009) have been observed to feed on *Dinophysis*. In fact, when offered a mixture of prey organisms (diatoms, a cryptophyte, a ciliate, a raphidophyte, and dinoflagellates) *F. duplocampanaeforme* fed exclusively on *Dinophysis* spp. (Park and Kim, 2010). Similarly, ciliates such as tintinnids (e.g., *Favella*) co-occur with *Dinophysis* populations in the field (Santhanam and Srinivasap, 1996) and feed on *Dinophysis* (Maneiro et al., 1998; Maneiro et al., 2000). Thus, microplankton could constitute an important group of grazers for *Dinophysis*, and a possible target for chemical defences that should be further explored.

### 4.2 Extracellular toxins – active release or passive leakage?

When exposed to high (5 – 10 nM) copepodamide concentrations, DPTs were redistributed from intracellular compartments to the extracellular medium in both species, which increased two to three-fold (Fig. 4). It is unknown if the redistribution was actively controlled or resulted from increased leakage from injured or lysed cells. However, a notable fraction of total *Dinophysis* toxins is commonly found in seawater (MacKenzie et al., 2004; Jørgensen and Andersen, 2007; Johansen, 2008). Phycotoxins can serve to immobilise prey before ingestion, which is experimentally supported for *Dinophysis* (Giménez Papiol et al., 2016; Mafra Jr et al., 2016) and other microzooplankton (Skovgaard and Hansen, 2003; Sheng et al., 2010; Blossom et al., 2012). It is, however, unclear whether DPTs serve this function for *Dinophysis* or if other compounds are involved. Mafra Jr et al. (2016) reported that purified solutions of OA, dinophysistoxin-1 (DTX1), and PTX-2 were ineffective against *Mesodinium rubrum* prey. Cell-free culture medium imprinted by *Dinophysis* compounds, including free polyunsaturated fatty acids such as eicosapentaenoic acid and docosahexaenoic acid, decreased their motility and survival substantially. This suggests that *Dinophysis* likely use some allelochemicals to aid in prey capture and may have actively released DPTs into the extracellular medium for this purpose in our experiment. However, this does not account for the consistent differential increase in extracellular DPTs in only the high copepodamide treatments for both species and lacks sufficient experimental evidence to be conclusively supported regardless.

Alternatively, DPTs released into the environment could act as a form of chemical aposematism (Weldon, 2013; Rojas et al., 2017), increasing its concentration in the phycosphere to discourage grazers capable of sensing them. It seems implausible, however, given the high spatial dispersion of cells, even in moderate turbulence, for this kind of allelopathic signal to confer an adaptive benefit to the individual cell (Jonsson et al., 2009). Interestingly though, a recent experiment found that extracellular DPTs significantly increased for *D. acuminata* cells when exposed to grazing *Acartia tonsa* copepods (Ladds et al., 2024). Although this could simply be due to toxins leaking into the medium when cells were fed on by the copepods, it tentatively supports our current findings and may suggest that *Dinophysis* actively releases toxins into the extracellular environment as a grazer deterrent.

The redistribution of toxins could also be due to passive leaking from injured or lysed cells. Cells exposed to high copepodamide concentrations grew more slowly than control cells, although not significantly so, potentially indicating a toxic effect or a cost associated with the copepodamide treatments on *Dinophysis*. However, all treatment groups showed negligible growth, which suggests a more general cause, such as the physical handling associated with the transfer of cultures to freshly coated culture vessels. *Dinophysis* is indeed sensitive to handling and leak their cellular toxins when physically disturbed (Jørgensen and Andersen, 2007; Johansen, 2008). This handling would, however, likely affect the treatments similarly and does not sufficiently explain the differential increase in only the two highest copepodamide groups for both species. Alternatively, the poor growth and consequent lysis was caused by an increased allocation cost associated with DPT production and/or other, unknown, grazer induced metabolites (Ryderheim et al., 2021; Brown et al., 2022; Olesen et al., 2022; Park et al., 2023).

### 4.3 Metabolomics

We examined endometabolomic profiles for both *Dinophysis* species after exposure to copepodamides or a living *Acartia* sp. copepod. Untargeted high resolution mass spectrometry analyses revealed potential grazer deterrent candidates among features that co-upregulated with increased copepodamide concentration. Alternatively, features that were both co-upregulated and co-downregulated with copepodamide concentration may be biomarkers indicating a stress-response or a potential growth cost associated with chemical defences induced by the copepodamides. However, it remains unclear if, how, or to what extent the expression pattern in the affected features is linked to growth rate. Overall, in all but two annotated features choline was involved, either directly or in polar lipids. Importantly, a series of five unsaturated lysoDGCC lipids and two lysoPC-like lipids, with fatty acid chains from 16 to 28 carbons, were identified in *D. sacculus*.

Diacylglycerylcarboxyhydroxymethylcholine (DGCC) are a family of betaine lipids that can be produced by dinoflagellates and may be their dominant representatives (Flaim et al., 2014; Cañavate et al., 2016). Although the exact location and roles of these DGCCs and their deacylated forms (i.e. lyso DGCCs) are unknown, they have been reported as biomarkers of thermal tolerance, depending on their abundance or degrees of acylation and unsaturation (Flaim et al., 2014; Rosset et al., 2019; Roach et al., 2021). Interestingly, the only putatively annotated feature in *D. acuminata* was choline, which constitutes the polar head of DGCC betaine lipids, but whether it is linked with DGCC or not remains unknown. All the significantly affected features were upregulated with increasing copepodamide concentrations but not affected by the live copepod, which is supported by the PCAs and likely reflect a true physiological response (Fig. 6, Fig. S3). A more comprehensive analysis of the total pool and nature of lysoPCs and lysoDGCCs in *Dinophysis* species could help understanding their function and how they are affected by stressors. While these phospho- and betaine lipids could mainly be localised in extra-chloroplastic membranes, they may be primary acceptors of *de novo* formed fatty acid before being redistributed from the cytoplasm to the chloroplast where they participate in membrane lipid synthesis (Eichenberger and Gribi, 1997).

While extracellular fractions would have been interesting to analyse in this context of chemically mediated induction of grazer deterrents, metabolomic analyses were only performed on intracellular samples (i.e. endometabolomes) given the limited amount of material, which is inherent to the highly challenging cultivation of *Dinophysis*. In addition, this may have led to sensitivity issues (e.g. no detection of low intensity features) and could have contributed to the low number of annotated features (e.g. poorly informative MS/MS spectra). This is, however, common when trying to annotate marine microalgal metabolomes (Zendong et al., 2016; Gémin et al., 2021; Yon et al., 2024). Regardless, knowledge about grazer deterring metabolites in microalgae is generally poor, especially for effects induced by chemical cues such as copepodamides, except for “known toxins” such as PSTs or ASTs.

### 4.4 Ecosystem implications for HAB formation

Harmful algal blooms (HABs) incur damages to human health and economic activities worth billions of dollars annually (Hoagland and Scatasta, 2006; Kouakou and Poder, 2019). Copepods are believed to contribute to HAB formation primarily by two mechanisms, inducing upregulation of toxin production (Bergkvist et al., 2008; Selander et al., 2015; Tammilehto et al., 2015) and/or by selectively feeding on non-toxic competitors (Selander et al., 2006; Prevett et al., 2019; Ryderheim et al., 2021; Olesen et al., 2022). Dinoflagellates grow slowly compared to e.g., many diatoms and are poor competitors for nutrients compared with similarly sized organisms (Banse, 1982). Thus, harmful algae in general, and dinoflagellates in particular, may be dependent on defensive traits (Fistarol et al., 2004; Selander et al., 2019) and/or selective grazers (Irigoien et al., 2005; Selander et al., 2006; Prevett et al., 2019). Indeed, when copepods are largely removed by the invasive ctenophore *Mnemiopsis leidyi*, community structure changes in favour of faster growing diatoms (Tiselius and Møller, 2017). Defended dinoflagellates are also positively correlated to copepods in the ocean (Prevett et al., 2019), which is also true for DSTs in mussels on the Swedish west coast (Trapp et al., 2021). If DSTs in mussels are indeed causally related to copepods, our results suggest that it is not driven by grazer-induced production but rather by selective feeding on less defended taxa, or other variables correlated to copepod density, leading to increased relative abundance of *Dinophyis* spp (Carlsson et al., 1995; Xu and Kiørboe, 2018). Future research should aim to determine the functional relationships between *Dinophysis* and their predators, the mechanism(s) causing increased extracellular toxins, and whether *Dinophysis* produced toxins result in any adaptive benefits for *Dinophysis* in interactions with copepods and other grazers.

## Supporting information

Supplementary code & output

## CRediT authorship contribution statement

**Milad Pourdanandeh:** Conceptualization, Methodology, Validation, Formal analysis, Investigation, Data curation, Writing – original draft, Writing – review & editing, Visualisation, Project administration

**Véronique Séchet**: Methodology, Resources, Investigation, Writing-review and editing

**Liliane Carpentier**: Methodology, Resources, Investigation

**Damien Réveillon:** Conceptualization, Validation, Formal analysis, Investigation, Data curation, Visualisation, Writing – original draft, Writing – review & editing

**Fabienne Hervé:** Formal analysis, Investigation, Methodology, Data curation, Writing-Review & Editing

**Clarisse Hubert**: Formal analysis, Investigation

**Philipp Hess:** Conceptualization, writing – review & editing, project administration

**Erik Selander:** Conceptualization, Methodology, Investigation, Formal analysis, Writing – original draft, Writing - review & editing, Visualisation, Project administration

## Data availability

Data and analysis code are available at Zenodo: https://doi.org/10.5281/zenodo.11091359

## Acknowledgements

We thank two anonymous reviewers for their insightful comments and constructive suggestions, which greatly improved the quality and clarity of this manuscript. We also thank Jonathan N. Havenhand for valuable insight and discussions regarding the statistical analyses.

