## Supplementary code & output for "Effects of copepod chemical cues on intra- and extracellular toxins in two species of *Dinophysis*": Supplementary_code_output.html


Code 

- Show All Code
- Hide All Code

### Supplementary code & output

##### Statistical analyses and data visualisations of a *Dinophysis* toxin induction experiment

###### Milad Pourdanandeh

#### 2024-07-16

### Load packages and set bibliography

```
# Check if the "pacman" package is installed; if not, install it
if (!require(pacman)) {
  install.packages("pacman")
}

# Load all packages with pacman
# if a package is not installed, pacman will install the package before loading
pacman::p_load(
  devtools,      # Tools for R package development
  readr,         # Fast reading of rectangular data (like CSV files)
  ggpubr,        # 'ggplot2' Based Publication Ready Plots
  multcomp,      # Simultaneous Inference in General Parametric Models
  rstatix,       # Pipe-friendly framework for basic statistical tests
  broom,         # Convert statistical analysis objects into tidy tibbles
  rcompanion,    # Functions to Support Extension Education Program Evaluation
  car,           # Companion to Applied Regression
  DescTools,     # Tools for Descriptive Statistics
  Rmisc,         # R miscellaneous functions
  viridis,       # Colorblind-Friendly Color Maps for R
  Hmisc,         # Harrell Miscellaneous
  ggthemes,      # Extra themes, scales, and geoms for 'ggplot2'
  AICcmodavg,    # Model Selection and Multimodel Inference
  ggpmisc,       # Miscellaneous Extensions to 'ggplot2'
  knitr,         # A general-purpose package for dynamic report generation in R
  chunkhooks,    # Custom Hooks for knitr Code Chunks
  kableExtra,    # Construct Complex Table with 'kable()'
  wesanderson,   # A 'ggplot2' Palette Generator Inspired by Wes Anderson Movies
  tidyverse      # A collection of R packages for data science
)

# Write the list of currently loaded packages to a .bib file
# .packages() retrieves a character vector of the names of currently attached packages
# write_bib creates a BibTeX file, which is useful for creating references in LaTeX documents
write_bib(
  .packages(),   # List of currently attached packages
  "packages.bib" # Output file name
)
```

### Purpose

Here we present the R-code used to analyse the results of the toxin
induction experiment on two species of *Dinophysis*
dinoflagellates, presented in our paper:

Effects of copepod chemical cues on intra- and extracellular toxins
in two species of *Dinophysis*.

### Overview

#### Experimental organisms

- *Dinophysis sacculus* (IFR-DSa-01Lt, accession no. MT 37
  1867)
- *Dinophysis acuminata* (IFR-DAu-01Ey, accession no. MT
  365104)

#### Datasets

- *Analysis\_data\_per\_mL.csv* contains all cell counts,
  growth rates, and toxin values for both species of *Dinophysis*
  used in this induction experiment.
- *pca\_score\_sacculus.csv* contains the scores of the first
  two principal components of the PCA performed on LC-HRMS derived
  metabolomic profiles of *D. sacculus*, in both positive and
  negative ion mode.
- *pca\_score\_acuminata.csv* contains the scores of the first
  two principal components of the PCA performed on LC-HRMS derived
  metabolomic profiles of *D. acuminata*, in both positive and
  negative ion mode.

### Availability

This analysis pipeline and the data sets used in it are publicly
avaialble at Zenodo: https://doi.org/10.5281/zenodo.11091359

### Analysis

#### Data import and preparation

##### Set the working directory and seed.

```
# Set the working directory to the specified path
# This path should be adjusted to match the user's file system structure
setwd("C:/Users/xpoumi/OneDrive/1. PhD GU/1. Research/Dinophysis induction/Dinophysis induction experiment")

# Set the seed for random number generation to ensure reproducibility
# Setting the seed ensures that random number generation produces the same results each time the code is run
set.seed(42)
```

##### Load and encode toxin data

```
# Read the data from a delimited file into a data frame
data_dinophysis_induction <- read_delim(
  "Analysis_data_per_mL.csv",  # file name
  delim = ";",                 # delimiter is a semicolon
  escape_double = FALSE,       # double quotes are not used to escape characters
  trim_ws = TRUE               # trim leading and trailing whitespace
)

# Coerce the imported data into a data frame
data_dinophysis_induction <- as.data.frame(data_dinophysis_induction)

# Set the order of levels in the factor "Treatment"
data_dinophysis_induction$Treatment <- factor(
  data_dinophysis_induction$Treatment,
  levels = c('Start', 'Control', 'Copepod', '1 nM', '5 nM', '10 nM')  # specific order for levels
)

# Set the order of levels in the factor "Species"
data_dinophysis_induction$Species <- factor(
  data_dinophysis_induction$Species,
  levels = c('D. sacculus', 'D. acuminata')  # specific order for levels
)
```

Create new data frame for analysis with the relevant columns from the
original dataset, rename the columns for easier use and calculate some
new parameters (totals and proportions)

```
# Filter the data to exclude the "Start" treatment and select specific columns
data_both_species <- 
  filter(data_dinophysis_induction, Treatment != "Start") %>%
  dplyr::select(
    Treatment = Treatment,           # treatment group
    Species = Species,               # species type
    Cop_conc = Cop_Conc,             # copepod concentration
    Growth_Rate_day = `Growth_rate_day-1`, # growth rate per day
    OA_free_intra = `OA_intracellular_ng_mL-1`, # intracellular OA free
    OA_hyd_intra = `OAHyd_intracellular_ng_mL-1`, # intracellular OA hydrolyzed
    PTX2_intra = `PTX2_intracellular_ng_mL-1`,   # intracellular PTX2
    C9_intra = `C9_diolOA_intracellular_ng_mL-1`, # intracellular C9 diol OA
    C9_iso_intra = `C9_diolOAisomer_intracellular_ng_mL-1`, # intracellular C9 diol OA isomer
    OA_extra = `OA_extracellular_ng_mL-1`, # extracellular OA
    PTX2_extra = `PTX2_extracellular_ng_mL-1`, # extracellular PTX2
    C9_extra = `C9_diolOA_extracellular_ng_mL-1`, # extracellular C9 diol OA
    C9_iso_extra = `C9_diolOAisomer_extracellular_ng_mL-1` # extracellular C9 diol OA isomer
  ) %>%
  
  # Create new variables for total toxin measurements and proportions
  mutate(
    OA_intra_total = OA_hyd_intra, # total intracellular OA
    PTX2_intra_total = PTX2_intra, # total intracellular PTX2
    C9_intra_total = C9_intra + C9_iso_intra, # total intracellular C9
    OA_extra_total = OA_extra, # total extracellular OA
    PTX2_extra_total = PTX2_extra, # total extracellular PTX2
    C9_extra_total = C9_extra + C9_iso_extra, # total extracellular C9
    OA_total = OA_intra_total + OA_extra_total, # total OA
    PTX2_total = PTX2_intra_total + PTX2_extra_total, # total PTX2
    C9_total = C9_intra_total + C9_extra_total, # total C9
    Tox_intra_total = OA_intra_total + PTX2_intra_total + C9_intra_total, # total intracellular toxins
    Tox_extra_total = OA_extra_total + PTX2_extra_total + C9_extra_total, # total extracellular toxins
    Tox_total = Tox_intra_total + Tox_extra_total, # total toxins
    prop_intra = Tox_intra_total / Tox_total, # proportion of intracellular toxins
    prop_extra = Tox_extra_total / Tox_total, # proportion of extracellular toxins
    prop_OA = OA_total / Tox_total, # proportion of OA
    prop_PTX2 = PTX2_total / Tox_total, # proportion of PTX2
    prop_C9 = C9_total / Tox_total # proportion of C9
  )

# Set the order of levels in the factor "Treatment"
data_both_species$Treatment <- factor(
  data_both_species$Treatment,
  levels = c('Control', 'Copepod', '1 nM', '5 nM', '10 nM') # specific order for levels
)
```

#### Analysis of growth rate

##### Exploratory data analysis and visualisation

###### Summary statistics table

```
# Calculate summary statistics
desc_both_growth <- data_both_species %>%
  dplyr::select(Species, Treatment, Growth_Rate_day) %>% # select relevant columns
  group_by(Species, Treatment) %>% # group by Species and Treatment
  summarise(
    N = length(Growth_Rate_day), # number of observations
    Mean = mean(Growth_Rate_day), # mean growth rate
    St.Dev = sd(Growth_Rate_day), # standard deviation
    SEM = St.Dev / sqrt(N), # standard error of the mean
    CI = qt(0.975, df = N - 1) * SEM, # 95% confidence interval
    LCI = Mean - CI, # lower confidence interval limit
    HCI = Mean + CI, # upper confidence interval limit
  ) %>%
  mutate(across(where(is.numeric), round, digits = 4)) %>% # round numeric variables to 4 digits
  as.data.frame() # convert to data frame
```

```
# Print the table with styling
desc_both_growth %>% 
  kable() %>% 
  kable_styling("striped") %>% 
  scroll_box(width = "100%", height = "450px")
```

Table C1.

| Species | Treatment | N | Mean | St.Dev | SEM | CI | LCI | HCI |
| --- | --- | --- | --- | --- | --- | --- | --- | --- |
| D. sacculus | Control | 3 | 0.0203 | 0.0819 | 0.0473 | 0.2034 | -0.1831 | 0.2237 |
| D. sacculus | Copepod | 3 | 0.0255 | 0.0347 | 0.0200 | 0.0862 | -0.0607 | 0.1118 |
| D. sacculus | 1 nM | 3 | 0.0400 | 0.1595 | 0.0921 | 0.3962 | -0.3562 | 0.4361 |
| D. sacculus | 5 nM | 3 | -0.1009 | 0.0799 | 0.0461 | 0.1984 | -0.2993 | 0.0975 |
| D. sacculus | 10 nM | 3 | -0.0532 | 0.2618 | 0.1512 | 0.6504 | -0.7036 | 0.5972 |
| D. acuminata | Control | 3 | 0.0418 | 0.0743 | 0.0429 | 0.1845 | -0.1427 | 0.2263 |
| D. acuminata | Copepod | 3 | 0.0533 | 0.0858 | 0.0496 | 0.2132 | -0.1600 | 0.2665 |
| D. acuminata | 1 nM | 3 | -0.0059 | 0.0683 | 0.0394 | 0.1696 | -0.1755 | 0.1637 |
| D. acuminata | 5 nM | 3 | -0.0803 | 0.0240 | 0.0138 | 0.0596 | -0.1399 | -0.0207 |
| D. acuminata | 10 nM | 3 | -0.0617 | 0.0418 | 0.0241 | 0.1038 | -0.1655 | 0.0421 |

###### Visual data exploration

Get Hex-codes for the “Moonrise3” palette from the “wesanderson”
package

```
wes_palettes$Moonrise3
```

```
## [1] "#85D4E3" "#F4B5BD" "#9C964A" "#CDC08C" "#FAD77B"
```

Display palette

```
wes_palette("Moonrise3")
```

Make custom colour palette for treatments

```
# Define a custom color palette using "Moonrise3" from the "wesanderson" package
# Reverse the order of the last three colours
custom_wes_palette <- c("#85D4E3", "#F4B5BD", "#FAD77B", "#CDC08C", "#78744B")
```

Box plots Grouped by treatment

```
# Plot the growth rate by treatment and species using ggplot2
data_both_species %>%
  ggplot(aes(
    x = Treatment, 
    y = Growth_Rate_day, 
    fill = Species # fill color: Species
  )) +
  
  geom_hline(yintercept = 0) + # add a horizontal line at y = 0

  scale_y_continuous(
    limits = c(-0.4, 0.4), # set y-axis limits
    n.breaks = 20, # set number of breaks on the y-axis
    expand = c(0, 0) # remove expansion at the axis limits
  ) +

  geom_boxplot(position = position_dodge(1)) + # add boxplots with dodged positions
  geom_point(position = position_dodge(1)) + # add points with dodged positions

  theme_classic() + # use classic theme
  theme(legend.position = "top") + # place the legend at the top

  scale_fill_brewer(palette = "Dark2") # use "Dark2" palette for fill colors
```

Figure C1.

Grouped by species

```
# Plot the growth rate by species and treatment using ggplot2
data_both_species %>%
  ggplot(aes(
    x = Species, # x-axis: Species
    y = Growth_Rate_day, # y-axis: Growth Rate per day
    fill = Treatment # fill color: Treatment
  )) +
  geom_hline(yintercept = 0) + # add a horizontal line at y = 0

  scale_y_continuous(
    limits = c(-0.4, 0.4), # set y-axis limits
    n.breaks = 20, # set number of breaks on the y-axis
    expand = c(0, 0) # remove expansion at the axis limits
  ) +

  geom_boxplot(position = position_dodge(1)) + # add boxplots with dodged positions
  geom_point(position = position_dodge(1)) + # add points with dodged positions

  theme_classic() + # use classic theme
  theme(legend.position = "top") + # place the legend at the top

  scale_fill_manual(values = custom_wes_palette) # use custom "Moonrise3" palette for fill colors
```

Figure C2.

We see the same general trend in both species, which indicates that
there is no interaction effect between the two factors. Also, there does
not seem to be any large difference in growth rate between the two
species.

We will make a scatter plot to see if the there is any linear
relationship between the growth rate and copepodamide concentration.
Observe that we filter out the Treatment level “Copepod”.

First, with the species separated:

```
# Filter out the "Copepod" treatment and plot growth rate by copepod concentration
data_both_species %>%
  filter(Treatment != "Copepod") %>%
  
  ggplot(aes(
    x = Cop_conc, # x-axis: Copepod concentration
    y = Growth_Rate_day # y-axis: Growth Rate per day
  )) +
  
  geom_point(
    size = 3, # size of points
    alpha = 0.5 # transparency of points
  ) +
  facet_wrap(~Species, nrow = 2) + # create separate plots for each species
  scale_x_continuous(n.breaks = 10) + # set number of breaks on x-axis
  
  stat_poly_line() + # add polynomial regression line
  stat_poly_eq(
    aes(label = after_stat(eq.label)), # add equation label
    label.x = 0.3 # position of equation label on x-axis
  ) +
  stat_poly_eq(
    label.y = 0.85, # position of equation label on y-axis
    label.x = 0.3 # position of equation label on x-axis
  ) +
  stat_correlation(
    mapping = aes(label = after_stat(r.label)), # add correlation coefficient
    label.y = 0.7, # position of correlation label on y-axis
    label.x = 0.3 # position of correlation label on x-axis
  ) +
  theme_classic() # use classic theme
```

Figure C3.

Secondly, with the two species pooled

```
# Filter out the "Copepod" treatment and plot growth rate by copepod concentration
data_both_species %>% 
  filter(Treatment != "Copepod") %>% # filter out "Copepod" treatment
  
  ggplot(aes(
    x = Cop_conc, # x-axis: Copepod concentration
    y = Growth_Rate_day # y-axis: Growth Rate per day
  )) +
  
  geom_point(
    size = 3, # size of points
    alpha = 0.5 # transparency of points
  ) +
  
  scale_y_continuous(
    limits = c(-0.4, 0.4), # set y-axis limits
    n.breaks = 9 # set number of breaks on y-axis
  ) +
  
  scale_x_continuous(
    n.breaks = 10 # set number of breaks on x-axis
  ) +
  
  stat_poly_line() + # add polynomial regression line
  
  stat_poly_eq(
    aes(label = after_stat(eq.label)), # add equation label
    label.x = 0.3 # position of equation label on x-axis
  ) +
  
  stat_poly_eq(
    label.y = 0.9, # position of second equation label on y-axis
    label.x = 0.3 # position of second equation label on x-axis
  ) +
  
  stat_correlation(
    mapping = use_label("p"), # add p-value
    label.y = 0.83, # position of p-value label on y-axis
    label.x = 0.3 # position of p-value label on x-axis
  ) +
  
  theme_classic() # use classic theme
```

Figure C4.

##### Statistical analysis

We fit a Two-Way ANOVA with growth rate as response variable, and
treatment and species as categorical predictor variables, including the
interaction between the two factors.

```
# Fit a linear model with interaction between Species and Treatment
growth_model_two_fact <- lm(Growth_Rate_day ~ Species * Treatment, 
                            data = data_both_species)

# Check the assumption of normality
hist(growth_model_two_fact$residuals, breaks = 12) # Histogram of residuals
```

```
shapiro_test(growth_model_two_fact$residuals) # Shapiro-Wilk test for normality
```

```
## # A tibble: 1 × 3
##   variable                        statistic p.value
##   <chr>                               <dbl>   <dbl>
## 1 growth_model_two_fact$residuals     0.934  0.0630
```

```
# Check the assumption of homogeneity of variances
levene_test(Growth_Rate_day ~ Treatment * Species, 
            data = data_both_species) # Levene's test for homogeneity of variances
```

```
## # A tibble: 1 × 4
##     df1   df2 statistic     p
##   <int> <int>     <dbl> <dbl>
## 1     9    20     0.953 0.504
```

```
# Perform the ANOVA
anova_test(growth_model_two_fact, 
           effect.size = "pes", 
           detailed = TRUE) # ANOVA with partial eta-squared effect size
```

```
## ANOVA Table (type II tests)
## 
##              Effect      SSn   SSd DFn DFd     F     p p<.05      pes
## 1           Species 7.21e-05 0.256   1  20 0.006 0.941       0.000281
## 2         Treatment 8.20e-02 0.256   4  20 1.590 0.216       0.241000
## 3 Species:Treatment 6.00e-03 0.256   4  20 0.111 0.977       0.022000
```

```
# Summary of the ANOVA
summary(aov(growth_model_two_fact)) # Detailed summary of the ANOVA
```

```
##                   Df  Sum Sq  Mean Sq F value Pr(>F)
## Species            1 0.00007 0.000072   0.006  0.941
## Treatment          4 0.08150 0.020376   1.590  0.216
## Species:Treatment  4 0.00568 0.001419   0.111  0.977
## Residuals         20 0.25627 0.012813
```

There are no significant main or interaction effects. Treatment
explains much more of the variation that remains unexplained by the
other model variables (24.1%) than species (0.03%) or their interaction
(2.2%) does.

##### Final growth plot

```
# Define a custom color palette for species
palette_species <- c("black", "grey70")

# Plot growth rate by treatment and species using ggplot2
data_both_species %>% 
  ggplot(aes(
    x = Treatment, # x-axis: Treatment
    y = Growth_Rate_day, # y-axis: Growth Rate per day
    colour = Species # color: Species
  )) +
  
  scale_y_continuous(
    limits = c(-0.8, 0.8), # set y-axis limits
    n.breaks = 9, # set number of breaks on y-axis
    expand = c(0, 0) # remove expansion at the axis limits
  ) +
  
  geom_hline(yintercept = 0) + # add a horizontal line at y = 0
  
  # Add jittered points to show replicate values
  geom_jitter(
    aes(
      x = Treatment,
      y = Growth_Rate_day,
      colour = Species
    ),
    size = 2, # size of points
    alpha = 0.4, # transparency of points
    shape = 18, # shape of points
    position = position_jitterdodge(
      jitter.width = 0.3, # width of jitter
      dodge.width = 1 # width of dodge
    )
  ) +
  
  # Add mean points
  geom_point(
    data = desc_both_growth, # data for mean points
    aes(
      y = Mean, # y-axis: Mean value
      colour = Species # color by Species
      ),
    size = 4, # size of mean points
    shape = 20, # shape of mean points
    position = position_dodge(width = 0.4) # position of mean points
    ) +
  
  # Add 95% Confidence intervals
  geom_errorbar(
    data = desc_both_growth, # data for error bars
    aes(
      y = Mean, # y-axis: Mean value
      ymin = LCI, # lower confidence interval
      ymax = HCI, # upper confidence interval
      colour = Species # color by Species
      ),
    linewidth = 0.5, # line width of error bars
    width = 0.25, # width of error bars
    position = position_dodge(width = 0.4), # position of error bars
    show.legend = FALSE # do not show legend for error bars
    ) +
  
  xlab("") + # remove x-axis label
  
  ylab(bquote(Growth~rate~(day^-1))) + # y-axis label with mathematical annotation
  
  scale_colour_manual(values = palette_species) + # use custom color palette
  
  theme_classic() + # use classic theme
  
  theme(
    legend.position = c(0.2, 0.95), # position the legend (x, y)
    
    legend.title = element_blank(), # remove legend title
    
    plot.margin = unit(c(0.3, 0, -0.15, 0), "cm"), # set plot margins (top, right, bottom, left)
    
    axis.title.y = element_text(
      vjust = 0, # vertical adjustment of y-axis title
      size = 10 # font size of y-axis title
    ),
    
    axis.text.y = element_text(
      colour = "black", # color of y-axis text
      size = 8 # font size of y-axis text
    ),
    
    axis.text.x = element_text(
      colour = "black", # color of x-axis text
      size = 7.5 # font size of x-axis text
    ),
    
    strip.background = element_rect(
      linetype = "blank" # no line type for strip background
    ),
    
    legend.background = element_rect(
      fill = 'transparent', # transparent fill for legend background
      colour = 'transparent' # transparent border for legend background
    ),
    
    legend.box.background = element_rect(
      fill = 'transparent', # transparent fill for legend box background
      colour = 'transparent' # transparent border for legend box background
    ),
    
    legend.key.size = unit(1, "mm"), # size of legend keys
    
    legend.text = element_text(
      colour = "black", # color of legend text
      size = 8, # font size of legend text
      face = "italic" # italic font style for legend text
    )
  )
```

Figure C5. Growth rate of *D. sacculus* (grey) and *D.
acuminata* (black) after 68.5 hours of exposure to three
concentrations of chemical cues from copepods (copepodamides), a
positive control (one live *Acartia* sp.) or a control treatment.
Partially transparent diamonds are individual growth rates in each
treatment level (n = 3), circles are the mean growth rates and error
bars are the 95% confidence intervals of the means.

```
# Save plot, commented out of use for knitting
# ggsave(path = "figures_manuscript", "Growth_plot_Dino_mL.tiff", width = 75, height = 65, units = "mm", dpi=700)
```

#### Analysis of total toxins

The primary and central analysis of this experiment is whether the
total amount of toxins produced is affected by the risk of grazing,
i.e., when exposed to chemical cues from copepods, and if this effect
differs between *Dinophysis* species. We will begin by visually
exploring the data.

##### Exploratory data analysis and visualisation

###### Summary statistics table

```
# Calculate summary statistics for total toxin content
desc_both_total_toxins <- data_both_species %>%
  dplyr::select(Species, Treatment, Tox_total) %>% # select relevant columns
  group_by(Species, Treatment) %>% # group by Species and Treatment
  summarise(
    N = length(Tox_total), # number of observations
    Mean = mean(Tox_total), # mean total toxin content
    St.Dev = sd(Tox_total), # standard deviation
    SEM = St.Dev / sqrt(N), # standard error of the mean
    CI = qt(0.975, df = N - 1) * SEM, # 95% confidence interval
    LCI = Mean - CI, # lower confidence interval limit
    HCI = Mean + CI, # upper confidence interval limit
  ) %>%
  mutate(across(where(is.numeric), round, digits = 4)) %>% # round numeric variables to 4 digits
  as.data.frame() # convert to data frame
```

```
# Print the table with styling
desc_both_total_toxins %>% 
  kable() %>% 
  kable_styling("striped") %>% 
  scroll_box(width = "100%", height = "450px")
```

Table C2.

| Species | Treatment | N | Mean | St.Dev | SEM | CI | LCI | HCI |
| --- | --- | --- | --- | --- | --- | --- | --- | --- |
| D. sacculus | Control | 3 | 146.7229 | 26.8830 | 15.5209 | 66.7812 | 79.9417 | 213.5040 |
| D. sacculus | Copepod | 3 | 159.9649 | 15.5693 | 8.9889 | 38.6762 | 121.2887 | 198.6411 |
| D. sacculus | 1 nM | 3 | 182.0866 | 71.8210 | 41.4659 | 178.4132 | 3.6734 | 360.4998 |
| D. sacculus | 5 nM | 3 | 146.8561 | 18.0199 | 10.4038 | 44.7638 | 102.0923 | 191.6199 |
| D. sacculus | 10 nM | 3 | 185.3710 | 81.8335 | 47.2466 | 203.2858 | -17.9148 | 388.6568 |
| D. acuminata | Control | 3 | 122.0882 | 13.0249 | 7.5199 | 32.3557 | 89.7325 | 154.4439 |
| D. acuminata | Copepod | 3 | 147.1946 | 31.8878 | 18.4104 | 79.2138 | 67.9808 | 226.4083 |
| D. acuminata | 1 nM | 3 | 134.2719 | 15.7391 | 9.0870 | 39.0982 | 95.1737 | 173.3701 |
| D. acuminata | 5 nM | 3 | 119.4468 | 10.8449 | 6.2613 | 26.9403 | 92.5066 | 146.3871 |
| D. acuminata | 10 nM | 3 | 125.3996 | 11.4609 | 6.6170 | 28.4705 | 96.9291 | 153.8701 |

###### Visual data exploration

Grouped by treatment

```
# Plot total toxin content, grouped by treatment and coloured by species
data_both_species %>% 
  ggplot(aes(
    x = Treatment, # x-axis: Treatment
    y = Tox_total, # y-axis: Total Toxin content
    fill = Species # fill color: Species
  )) + 
  
  scale_y_continuous(
    n.breaks = 14 # set number of breaks on y-axis
  ) +
  
  geom_boxplot(
    position = position_dodge(1) # add boxplots with dodged positions
  ) +
  
  geom_point(
    position = position_dodge(1), # add points with dodged positions
    show.legend = FALSE # do not show legend for points
  ) +
  
  theme_classic() + # use classic theme
  
  theme(
    legend.position = "top" # position the legend at the top
  ) +
  
  scale_fill_brewer(
    palette = "Dark2" # use "Dark2" palette for fill colors
  ) +
  
  ylab(bquote(Toxin~content~(ng~mL^-1))) # y-axis label with mathematical annotation
```

Figure C6.

Grouped by species

```
# Plot total toxin content grouped by species and coloured by treatment
data_both_species %>% 
  ggplot(aes(
    x = Species, # x-axis: Species
    y = Tox_total, # y-axis: Total Toxin content
    fill = Treatment # fill color: Treatment
  )) + 
  
  scale_y_continuous(
    n.breaks = 14 # set number of breaks on y-axis
  ) +

  geom_boxplot(
    position = position_dodge(1) # add boxplots with dodged positions
  ) +
  
  geom_point(
    position = position_dodge(1), # add points with dodged positions
    show.legend = FALSE # do not show legend for points
  ) +
  
  theme_classic() + # use classic theme
  
  theme(
    legend.position = "top" # position the legend at the top
  ) +
  
  scale_fill_manual(
    values = custom_wes_palette # use custom "Moonrise3" palette for fill colors
  ) +
  
  ylab(bquote(Toxin~content~(ng~mL^-1))) # y-axis label with mathematical annotation
```

Figure C7.

There appears to be a similar effect of both copepodamide and copepod
exposure on toxin production in both species of *Dinophysis*.
This indicates potential main effects of both treatment and species, but
no interaction effect (i.e. the effect of one factor does not depend on
the other factor). Note that these plots do not take the effect of
growth rate on toxin content into account.

Histogram of toxin content for all treatments, coloured by
treatment

```
# Plot histogram of total toxin content by treatment using ggplot2
data_both_species %>% 
  ggplot(aes(x = Tox_total)) +
  
  geom_histogram(
    aes(fill = Treatment), # fill color by Treatment
    colour = "black", # outline color of bins
    bins = 15 # number of bins
  ) +
  
  scale_y_continuous(
    n.breaks = 6, # set number of breaks on y-axis
    minor_breaks = NULL, # remove minor breaks
    expand = c(0, 0) # remove expansion at the axis limits
  ) +
  
  scale_x_continuous(
    n.breaks = 8 # set number of breaks on x-axis
  ) +
  
  coord_cartesian(
    xlim = c(100, 300) # set limits of the viewing window on x-axis
  ) +
  
  scale_fill_manual(
    values = custom_wes_palette # use custom "Moonrise3" palette for fill colors
  ) +
  
  theme_classic() + # use classic theme
  
  theme(
    legend.position = "top", # position the legend at the top
    legend.title = element_blank() # remove legend title
  ) +
  
  xlab(bquote(Toxin~content~(ng~mL^-1))) # x-axis label with mathematical annotation
```

Figure C8.

The toxin data largely seems to be approximately normally
distributed, although right skewed by two high toxin values.

Relationship between growth rate (independent variable) and total
toxins (dependent variable):

```
# Plot total toxin content vs. growth rate per day using ggplot2
data_both_species %>%
  ggplot(aes(
    x = Growth_Rate_day, # x-axis: Growth Rate per day
    y = Tox_total # y-axis: Total Toxin content
  )) +
  
  geom_point(
    size = 3, # size of points
    alpha = 0.5 # transparency of points
  ) +
  
  stat_poly_line() + # add polynomial regression line
  stat_poly_eq(
    aes(label = after_stat(eq.label)), # add equation label
    label.x = 0.3 # position of equation label on x-axis
  ) +
  stat_poly_eq(
    label.y = 0.9, # position of second equation label on y-axis
    label.x = 0.3 # position of second equation label on x-axis
  ) +
  stat_correlation(
    mapping = use_label("p"), # add p-value label
    label.y = 0.83, # position of p-value label on y-axis
    label.x = 0.3 # position of p-value label on x-axis
  ) +
  
  scale_y_continuous(
    limits = c(0, 300), # set y-axis limits
    breaks = c(0, 50, 100, 150, 200, 250, 300), # set y-axis breaks
    expand = c(0, 0) # remove expansion at the axis limits
  ) +
  
  scale_x_continuous(
    limits = c(-0.35, 0.3), # set x-axis limits
    breaks = c(-0.3, -0.2, -0.1, 0, 0.1, 0.2, 0.3), # set x-axis breaks
    expand = c(0, 0) # remove expansion at the axis limits
  ) +
  
  ylab(bquote(Total~toxins~(ng~mL^-1))) + # y-axis label with mathematical annotation
  xlab(bquote(Growth~rate~(day^-1))) + # x-axis label with mathematical annotation
  
  theme_classic() + # use classic theme
  
  theme(
    plot.margin = unit(c(3, 5, 0, 0), "mm") # set plot margins (top, right, bottom, left)
  )
```

Figure C9.

```
## ggsave(path = "figures_manuscript", "Scatter_tox_growth.tiff", width = 100, height = 100, units = "mm", dpi = 700)
```

Summary of linear model with growth rate and total toxins

```
# Fit a linear model for Tox_total as a function of Growth_Rate_day
tox_growth_model <- lm(Tox_total ~ Growth_Rate_day, data = data_both_species)

# Display the summary of the linear model
summary(tox_growth_model)
```

```
## 
## Call:
## lm(formula = Tox_total ~ Growth_Rate_day, data = data_both_species)
## 
## Residuals:
##     Min      1Q  Median      3Q     Max 
## -48.205 -19.570  -2.499  10.753  74.203 
## 
## Coefficients:
##                 Estimate Std. Error t value Pr(>|t|)    
## (Intercept)      149.990      5.212  28.777  < 2e-16 ***
## Growth_Rate_day  251.710     48.399   5.201  1.6e-05 ***
## ---
## Signif. codes:  0 '***' 0.001 '**' 0.01 '*' 0.05 '.' 0.1 ' ' 1
## 
## Residual standard error: 28.37 on 28 degrees of freedom
## Multiple R-squared:  0.4913, Adjusted R-squared:  0.4732 
## F-statistic: 27.05 on 1 and 28 DF,  p-value: 1.602e-05
```

There is a positive effect of growth rate on DST content in our
experiment, further pointing towards the importance of correcting for
this effect in our analysis.

##### Statistical analyses

###### Model fitting and evaluation

We will use Aikaike’s information criterion to evaluate and choose
among the models that we have identified as appropriate to analyse our
data.

```
# Construct the models
lm_model_1 <- lm(Tox_total ~ Treatment, 
                 data = data_both_species) # ANOVA

lm_model_2 <- lm(Tox_total ~ Growth_Rate_day + Treatment, 
                 data = data_both_species) # ANCOVA

lm_model_3 <- lm(Tox_total ~ Species, 
                 data = data_both_species) # ANOVA

lm_model_4 <- lm(Tox_total ~ Growth_Rate_day + Species, 
                 data = data_both_species) # ANCOVA

lm_model_5 <- lm(Tox_total ~ Species * Treatment, 
                 data = data_both_species) # Two-way ANOVA

lm_model_6 <- lm(Tox_total ~ Growth_Rate_day + Treatment * Species, 
                 data = data_both_species) # Two-way ANCOVA with interaction

lm_model_7 <- lm(Tox_total ~ Growth_Rate_day + Treatment + Species, 
                 data = data_both_species) # Two-way ANCOVA without interaction

# Create a list of models
model.set <- list(lm_model_1, lm_model_2, lm_model_3, 
                  lm_model_4, lm_model_5, lm_model_6, lm_model_7)

model_names <- c("lm_model_1", "lm_model_2", "lm_model_3", 
                 "lm_model_4", "lm_model_5", "lm_model_6", "lm_model_7")

# Calculate AICc and compare models
aictab(model.set, modnames = model_names)
```

```
## 
## Model selection based on AICc:
## 
##             K   AICc Delta_AICc AICcWt Cum.Wt      LL
## lm_model_7  8 266.77       0.00   0.99   0.99 -121.96
## lm_model_4  4 277.30      10.53   0.01   1.00 -133.85
## lm_model_6 12 280.47      13.70   0.00   1.00 -119.06
## lm_model_2  7 291.48      24.71   0.00   1.00 -136.19
## lm_model_3  3 304.22      37.45   0.00   1.00 -148.65
## lm_model_1  6 317.22      50.45   0.00   1.00 -150.78
## lm_model_5 11 328.69      61.92   0.00   1.00 -146.01
```

Models that contain growth rate as covariate are the most
parsimonious of the candidates. We will investigate if there is any
difference between the multi-factorial ANCOVA with the interaction
included (model 6) versus the two uni-factorial ANCOVA models (2 &
4)

```
# Perform ANOVA to compare lm_model_2 and lm_model_6
anova(lm_model_2, lm_model_6)
```

```
## Analysis of Variance Table
## 
## Model 1: Tox_total ~ Growth_Rate_day + Treatment
## Model 2: Tox_total ~ Growth_Rate_day + Treatment * Species
##   Res.Df     RSS Df Sum of Sq      F    Pr(>F)    
## 1     24 15411.2                                  
## 2     19  4917.9  5     10493 8.1081 0.0003083 ***
## ---
## Signif. codes:  0 '***' 0.001 '**' 0.01 '*' 0.05 '.' 0.1 ' ' 1
```

```
# Perform ANOVA to compare lm_model_4 and lm_model_6
anova(lm_model_4, lm_model_6)
```

```
## Analysis of Variance Table
## 
## Model 1: Tox_total ~ Growth_Rate_day + Species
## Model 2: Tox_total ~ Growth_Rate_day + Treatment * Species
##   Res.Df     RSS Df Sum of Sq      F   Pr(>F)   
## 1     27 13183.3                                
## 2     19  4917.9  8    8265.4 3.9916 0.006295 **
## ---
## Signif. codes:  0 '***' 0.001 '**' 0.01 '*' 0.05 '.' 0.1 ' ' 1
```

The multifactorial ANCOVA that includes the interaction between
treatment and species is significantly better than the unifactorial
models.

Now we compare the two multi-factorial models to see if removal of
the interaction term results in the best model (which the AICc values
suggest)

```
# Perform ANOVA to compare lm_model_7 and lm_model_6
anova(lm_model_7, lm_model_6)
```

```
## Analysis of Variance Table
## 
## Model 1: Tox_total ~ Growth_Rate_day + Treatment + Species
## Model 2: Tox_total ~ Growth_Rate_day + Treatment * Species
##   Res.Df    RSS Df Sum of Sq      F Pr(>F)
## 1     23 5965.0                           
## 2     19 4917.9  4    1047.1 1.0114 0.4263
```

These models do not differ from each other, we will therefore use the
multi-factorial ANCOVA that includes the interaction term for our
analysis. We will also use estimates of the covariate (i.e. growth rate)
corrected/adjusted mean toxin content for final visualisation.

###### Checking assumptions of the analysis

###### Linearity assumption

Here we make sure that there is a linear relationship between the
covariate and the response variable. For each species:

```
# Scatter plot of growth rate vs. total toxin content with regression lines, facetted by species
ggscatter(data_both_species, 
          x = "Growth_Rate_day", 
          y = "Tox_total",
          add = "reg.line") + # add regression line
  facet_wrap(~Species) + # facet by species
  
  # Add regression line equation and R-squared value
  stat_regline_equation(aes(label = paste(..eq.label.., 
                                          ..rr.label.., 
                                          sep = "~~~~")),
                        label.x = -0.2) + # position of the label on x-axis
  
  scale_y_continuous(
    n.breaks = 10 # set number of breaks on y-axis
  ) +
  
  xlab(bquote(Growth~rate~(day^-1))) + # x-axis label with mathematical annotation
  ylab(bquote(Toxin~content~(ng~mL^-1))) # y-axis label with mathematical annotation
```

Figure C10.

For each treatment within each species:

```
# Scatter plot of growth rate vs. total toxin content with regression lines, facetted by species, colored by treatment
ggscatter(data_both_species, 
          x = "Growth_Rate_day", 
          y = "Tox_total",
          color = "Treatment", # color points by Treatment
          add = "reg.line") + # add regression line
  facet_wrap(~Species) + # facet by species
  
  # Add regression line equation and R-squared value
  stat_regline_equation(aes(
    label = paste(..eq.label.., ..rr.label.., sep = "~~~~"), # add equation and R-squared value
    color = Treatment # color of label by Treatment
  ), 
  label.x = -0.3, # position of the label on x-axis
  label.y = c(280, 270, 260, 250, 240),# positions of the labels on y-axis
  show.legend = FALSE) + # Remove label object from the legend
  
  scale_y_continuous(
    n.breaks = 10 # set number of breaks on y-axis
  ) +
  
  scale_color_manual(
    values = custom_wes_palette # use custom "Moonrise3" palette for colors
  ) +
  
  xlab(bquote(Growth~rate~(day^-1))) + # x-axis label with mathematical annotation
  ylab(bquote(Toxin~content~(ng~mL^-1)))  # y-axis label with mathematical annotation
```

Figure C11.

For each treatment across both species:

```
# Scatter plot of growth rate vs. total toxin content with regression lines, facetted by treatment
ggscatter(data_both_species, 
          x = "Growth_Rate_day", 
          y = "Tox_total",
          add = "reg.line") + # add regression line
  facet_wrap(~Treatment, # facet by treatment
             nrow = 1, # number of rows
             ncol = 5) + # number of columns
  
  # Add regression line equation
  stat_regline_equation(aes(
    label = ..eq.label.., # add equation
  ), 
  label.x = -0.35, # position of the label on x-axis
  label.y = 250, # position of the label on y-axis
  parse = TRUE) + # parse labels for mathematical annotations
  
  #Add regression line R-squared
  stat_regline_equation(aes(
    label = ..rr.label.., # add R-squared value
  ), 
  label.x = -0.35, # position of the label on x-axis
  label.y = 240, # position of the label on y-axis
  parse = TRUE) + # parse labels for mathematical annotations
  
  scale_y_continuous(
    n.breaks = 10 # set number of breaks on y-axis
  ) +
  
  scale_x_continuous(
    n.breaks = 4 # set number of breaks on x-axis
  ) +
  
  scale_color_brewer(
    palette = "Dark2" # use "Dark2" palette for colors
  ) +
  
  xlab(bquote(Growth~rate~(day^-1))) + # x-axis label with mathematical annotation
  ylab(bquote(Toxin~content~(ng~mL^-1))) # y-axis label with mathematical annotation
```

Figure C12.

There is a positive linear relationship between the covariate (growth
rate) and the response variable (toxin content), as assessed visually
using these scatter plots.

###### Homogeneity of regression slopes

Here we make sure that there is no interaction between the covariate
and the two factors

```
# Combine the Treatment and Species columns into a single column named "group"
# Then perform an ANOVA test on Tox_total with the interaction term group * Growth_Rate_day
data_both_species %>%
  unite(
    col = "group", # new column name
    Treatment, # first column to combine
    Species, # second column to combine
  ) %>%
  
  anova_test(
    Tox_total ~ Growth_Rate_day * group   # ANOVA test formula with interaction term
  )
```

```
## ANOVA Table (type II tests)
## 
##                  Effect DFn DFd       F        p p<.05   ges
## 1       Growth_Rate_day   1  10 133.539 4.16e-07     * 0.930
## 2                 group   9  10  10.564 5.00e-04     * 0.905
## 3 Growth_Rate_day:group   9  10   1.839 1.78e-01       0.623
```

The assumption of homogeneity of regression slopes seems to hold, as
there is no interaction effect between the covariate and the grouped
factors. The effect of growth rate does seems to differ between the two
species though, which suggests that species should be separated in the
final analysis.

###### VIFs

```
# Calculate the Variance Inflation Factor (VIF) for the linear model
vif(lm(
  Tox_total ~ 
    Growth_Rate_day + # predictor: Growth Rate per day
    Treatment + # predictor: Treatment
    Species + # predictor: Species
    Treatment * Species + # interaction term: Treatment and Species
    Growth_Rate_day * Treatment + # interaction term: Growth Rate per day and Treatment
    Growth_Rate_day * Species + # interaction term: Growth Rate per day and Species
    Growth_Rate_day * Species * Treatment, # interaction term: Growth Rate per day, Species, and Treatment
  data = data_both_species # data set
))
```

```
##                                          GVIF Df GVIF^(1/(2*Df))
## Growth_Rate_day                     25.627057  1        5.062317
## Treatment                           90.327367  4        1.755810
## Species                              6.418713  1        2.533518
## Treatment:Species                 1733.851732  4        2.540250
## Growth_Rate_day:Treatment         1036.859941  4        2.382128
## Growth_Rate_day:Species             13.801802  1        3.715078
## Growth_Rate_day:Treatment:Species 3057.203815  4        2.726877
```

VIFs look good (i.e., not above ~10).

###### Normality of residuals

Here we assess the assumption that the residuals of the model are
approximately normally distributed.

```
# Extract model metrics for lm_model_6
# Augment the linear model with additional metrics
model.metrics <- augment(lm_model_6) %>%
  dplyr::select(-.hat, -.sigma, -.fitted) # Remove specific columns: .hat, .sigma, .fitted
```

We do this visually using a histogram and a QQ-plot Histogram:

```
# Plot a histogram of the residuals from the model metrics
model.metrics %>% 
  ggplot(aes(x = .resid)) + 
  geom_histogram(
    fill = "grey", # fill color of the bars
    colour = "black", # outline color of the bars
    binwidth = 5 # width of the bins
  ) +
  theme_classic() + # use classic theme
  scale_y_continuous(
    expand = c(0, 0) # remove expansion at the axis limits
  ) +
  xlab("Residuals") + # x-axis label
  ylab("Count") # y-axis label
```

Figure C13.

QQ-plot

```
# Create a Q-Q plot for the linear model lm_model_6 to check the distribution of studentized residuals
qqPlot(
  lm_model_6, # linear model
  grid = FALSE, # do not display grid
  ylab = "Studentised residuals" # y-axis label
)
```

Figure C14.

```
## [1]  6 11
```

The residuals appear to be approximately normally distributed, we
test this statistically with Shapiro-Wilks test.

```
# Perform Shapiro-Wilk test for normality on the residuals of the model
shapiro_test(model.metrics$.resid)
```

```
## # A tibble: 1 × 3
##   variable             statistic p.value
##   <chr>                    <dbl>   <dbl>
## 1 model.metrics$.resid     0.980   0.825
```

The assumption of normality holds true.

###### Outliers

To detect potential outliers, we check if the absolute value of any
of the standardised residuals are larger than 3.

```
# Convert model metrics to a data frame and filter for 
# observations with absolute standardized residuals greater than 3
model.metrics %>% 
  as.data.frame() %>% 
  filter(abs(.std.resid) > 3)
```

```
## [1] Tox_total       Growth_Rate_day Treatment       Species        
## [5] .resid          .cooksd         .std.resid     
## <0 rows> (or 0-length row.names)
```

There does not seem to be any strong outliers in our model.

Now we are ready to perform and summarise the total toxin
analysis.

###### Two-Way Analysis of Covariance

Our final model uses the total toxin mass of an experimental
replicate (normalised by sample volume) as response variable, species
and treatment as categorical variables (including their interaction) and
growth rate as covariate

```
# Fit a linear model for Tox_total as a function of Growth_Rate_day and the interaction between Treatment and Species
two_fact_totaltox <- lm(Tox_total ~ Growth_Rate_day + Treatment * Species,
                        data = data_both_species)

# Perform ANOVA on the linear model and calculate effect size (partial eta squared)
anova_test(two_fact_totaltox, 
           effect.size = "pes", # calculate partial eta squared
           detailed = TRUE) # provide detailed results
```

```
## ANOVA Table (type II tests)
## 
##              Effect       SSn      SSd DFn DFd      F        p p<.05   pes
## 1   Growth_Rate_day 24738.055 4917.873   1  19 95.574 7.57e-09     * 0.834
## 2         Treatment  7218.260 4917.873   4  19  6.972 1.00e-03     * 0.595
## 3           Species  9446.212 4917.873   1  19 36.495 8.23e-06     * 0.658
## 4 Treatment:Species  1047.142 4917.873   4  19  1.011 4.26e-01       0.176
```

There is a significant effect of the covariate and both main factors,
but no interaction effect. Because of the difference between the two
species, will analyse them with separate uni-factorial ANCOVAs.

###### One-way Analyses of Covariance

###### *D. sacculus*:

```
# Fit a linear model for Tox_total as a function of Growth_Rate_day and Treatment for D. sacculus
ancova_sacculus <- lm(Tox_total ~ Growth_Rate_day + Treatment,
                      data = filter(data_both_species, Species == "D. sacculus"))

# Perform ANOVA on the linear model and calculate effect size (partial eta squared)
anova_test(ancova_sacculus, 
           effect.size = "pes", # calculate partial eta squared
           detailed = TRUE) # provide detailed results
```

```
## ANOVA Table (type II tests)
## 
##            Effect      SSn      SSd DFn DFd      F        p p<.05   pes
## 1 Growth_Rate_day 23883.39 2406.202   1   9 89.332 5.71e-06     * 0.908
## 2       Treatment  6753.87 2406.202   4   9  6.315 1.10e-02     * 0.737
```

```
# Post-hoc pairwise comparisons using Tukey's method
sacculus.mc <- glht(ancova_sacculus, 
                    linfct = mcp(Treatment = 'Tukey')) # specify Treatment for multiple comparisons

# Display summary of the post-hoc comparisons
summary(sacculus.mc)
```

```
## 
##   Simultaneous Tests for General Linear Hypotheses
## 
## Multiple Comparisons of Means: Tukey Contrasts
## 
## 
## Fit: lm(formula = Tox_total ~ Growth_Rate_day + Treatment, data = filter(data_both_species, 
##     Species == "D. sacculus"))
## 
## Linear Hypotheses:
##                        Estimate Std. Error t value Pr(>|t|)   
## Copepod - Control == 0    11.51      13.35   0.862  0.90360   
## 1 nM - Control == 0       28.83      13.37   2.157  0.27630   
## 5 nM - Control == 0       40.39      14.01   2.882  0.10065   
## 10 nM - Control == 0      63.07      13.60   4.638  0.00817 **
## 1 nM - Copepod == 0       17.33      13.36   1.297  0.69944   
## 5 nM - Copepod == 0       28.88      14.07   2.053  0.31590   
## 10 nM - Copepod == 0      51.56      13.63   3.782  0.02707 * 
## 5 nM - 1 nM == 0          11.56      14.24   0.812  0.92039   
## 10 nM - 1 nM == 0         34.24      13.75   2.491  0.17596   
## 10 nM - 5 nM == 0         22.68      13.46   1.686  0.48603   
## ---
## Signif. codes:  0 '***' 0.001 '**' 0.01 '*' 0.05 '.' 0.1 ' ' 1
## (Adjusted p values reported -- single-step method)
```

Post-hoc table *D. saccuslus*

The 10 nM treatment differs significantly from both the control and
copepod treatments for *D. sacculus*, all other pairwise
contrasts are homogeneous.

###### *D. acuminata*:

```
# Fit a linear model for Tox_total as a function of Growth_Rate_day and Treatment for D. acuminata
ancova_acuminata <- lm(Tox_total ~ Growth_Rate_day + Treatment,
                       data = filter(data_both_species, Species == "D. acuminata"))

# Perform ANOVA on the linear model and calculate effect size (partial eta squared)
anova_test(ancova_acuminata, 
           effect.size = "pes", # calculate partial eta squared
           detailed = TRUE) # provide detailed results
```

```
## ANOVA Table (type II tests)
## 
##            Effect      SSn      SSd DFn DFd     F     p p<.05   pes
## 1 Growth_Rate_day 1494.999 1871.336   1   9 7.190 0.025     * 0.444
## 2       Treatment 1060.457 1871.336   4   9 1.275 0.348       0.362
```

All treatment groups are statistically homogeneous, there is no
effect of treatment on *D. acuminata*.

##### Final total toxin plot

To make a plot of the covariate (growth rate) adjusted means (&
95% CI) for total toxin content, we first predict the covariate
adjusted/corrected means and sampling error for each species ANCOVA
model. We will predict the means and sampling error at growth rate = 0
(the actual value does not matther though).

###### Predicted estimates for *D. sacculus* at growth rate = 0

```
# Create a new data frame for predictions with Growth_Rate_day set to 0 for D. sacculus
df_newdata_sacculus <- crossing(
  Growth_Rate_day = 0, # set Growth Rate per day to 0
  Species = levels(data_both_species$Species), # include all species levels
  Treatment = levels(data_both_species$Treatment) # include all treatment levels
) %>% 
  filter(Species == "D. sacculus") # filter for D. sacculus

# Predict Tox_total using the ANCOVA model for D. sacculus
preds_matrix_sacculus <- predict(
  ancova_sacculus, # model to use for predictions
  newdata = df_newdata_sacculus, # new data for predictions
  interval = "c" # prediction intervals (confidence)
)

# Display the new data frame and prediction results
df_newdata_sacculus
```

```
## # A tibble: 5 × 3
##   Growth_Rate_day Species     Treatment
##             <dbl> <chr>       <chr>    
## 1               0 D. sacculus 1 nM     
## 2               0 D. sacculus 10 nM    
## 3               0 D. sacculus 5 nM     
## 4               0 D. sacculus Control  
## 5               0 D. sacculus Copepod
```

```
preds_matrix_sacculus
```

```
##        fit      lwr      upr
## 1 168.8119 147.2215 190.4023
## 2 203.0495 181.2790 224.8200
## 3 180.3696 157.5575 203.1817
## 4 139.9773 118.5610 161.3936
## 5 151.4866 130.0350 172.9381
```

```
# Combine the new data frame with prediction results
sacculus_tox_predict_Growth0 <- bind_cols(
  df_newdata_sacculus, 
  preds_matrix_sacculus
) %>% 
  dplyr::rename(
    Mean = fit, # rename predicted mean
    LCI = lwr, # rename lower confidence interval
    HCI = upr # rename upper confidence interval
  )

# Set order of levels in factor Treatment
sacculus_tox_predict_Growth0$Treatment <- factor(
  sacculus_tox_predict_Growth0$Treatment,
  levels = c('Control', 'Copepod', '1 nM', '5 nM', '10 nM') # specific order for levels
)

# Display the final data frame with predictions
sacculus_tox_predict_Growth0
```

```
## # A tibble: 5 × 6
##   Growth_Rate_day Species     Treatment  Mean   LCI   HCI
##             <dbl> <chr>       <fct>     <dbl> <dbl> <dbl>
## 1               0 D. sacculus 1 nM       169.  147.  190.
## 2               0 D. sacculus 10 nM      203.  181.  225.
## 3               0 D. sacculus 5 nM       180.  158.  203.
## 4               0 D. sacculus Control    140.  119.  161.
## 5               0 D. sacculus Copepod    151.  130.  173.
```

###### Predicted estimated for *D. acuminata* at growth rate = 0

```
# Create a new data frame for predictions with Growth_Rate_day set to 0 for D. acuminata
df_newdata_acuminata <- crossing(
  Growth_Rate_day = 0, # set Growth Rate per day to 0
  Species = levels(data_both_species$Species), # include all species levels
  Treatment = levels(data_both_species$Treatment) # include all treatment levels
) %>% 
  filter(Species == "D. acuminata") # filter for D. acuminata

# Predict Tox_total using the ANCOVA model for D. acuminata
preds_matrix_acuminata <- predict(
  ancova_acuminata, # model to use for predictions
  newdata = df_newdata_acuminata, # new data for predictions
  interval = "c" # prediction intervals (confidence)
)

# Display the new data frame and prediction results
df_newdata_acuminata
```

```
## # A tibble: 5 × 3
##   Growth_Rate_day Species      Treatment
##             <dbl> <chr>        <chr>    
## 1               0 D. acuminata 1 nM     
## 2               0 D. acuminata 10 nM    
## 3               0 D. acuminata 5 nM     
## 4               0 D. acuminata Control  
## 5               0 D. acuminata Copepod
```

```
preds_matrix_acuminata
```

```
##        fit       lwr      upr
## 1 135.4153 116.55776 154.2729
## 2 137.3694 115.99999 158.7387
## 3 135.0231 112.05882 157.9873
## 4 113.9805  93.94396 134.0170
## 5 136.8613 116.10857 157.6139
```

```
# Combine the new data frame with prediction results
acuminata_tox_predict_Growth0 <- bind_cols(
  df_newdata_acuminata, 
  preds_matrix_acuminata
) %>% 
  dplyr::rename(
    Mean = fit, # rename predicted mean
    LCI = lwr, # rename lower confidence interval
    HCI = upr # rename upper confidence interval
  )

# Set order of levels in factor Treatment
acuminata_tox_predict_Growth0$Treatment <- factor(
  acuminata_tox_predict_Growth0$Treatment,
  levels = c('Control', 'Copepod', '1 nM', '5 nM', '10 nM') # specific order for levels
)

# Display the final data frame with predictions
acuminata_tox_predict_Growth0
```

```
## # A tibble: 5 × 6
##   Growth_Rate_day Species      Treatment  Mean   LCI   HCI
##             <dbl> <chr>        <fct>     <dbl> <dbl> <dbl>
## 1               0 D. acuminata 1 nM       135. 117.   154.
## 2               0 D. acuminata 10 nM      137. 116.   159.
## 3               0 D. acuminata 5 nM       135. 112.   158.
## 4               0 D. acuminata Control    114.  93.9  134.
## 5               0 D. acuminata Copepod    137. 116.   158.
```

###### Binding them together

```
# Ensure the levels of Treatment are set correctly before binding the data frames
sacculus_tox_predict_Growth0$Treatment <- factor(
  sacculus_tox_predict_Growth0$Treatment,
  levels = c('Control', 'Copepod', '1 nM', '5 nM', '10 nM')
)

acuminata_tox_predict_Growth0$Treatment <- factor(
  acuminata_tox_predict_Growth0$Treatment,
  levels = c('Control', 'Copepod', '1 nM', '5 nM', '10 nM')
)

# Bind the two data frames together
all_tox_predict_Growth0 <- bind_rows(sacculus_tox_predict_Growth0, 
                                     acuminata_tox_predict_Growth0)

# Ensure the levels of the Species factor are set correctly
all_tox_predict_Growth0$Species <- factor(
  all_tox_predict_Growth0$Species,
  levels = c('D. sacculus', 'D. acuminata')
)

# Ensure the levels of the Treatment factor are set correctly
all_tox_predict_Growth0$Treatment <- factor(
  all_tox_predict_Growth0$Treatment,
  levels = c('Control', 'Copepod', '1 nM', '5 nM', '10 nM')
)

# Order the data by Species and Treatment to verify the correct levels
all_tox_predict_Growth0 <- 
  all_tox_predict_Growth0 %>%
  arrange(Species, Treatment) %>% 
  rename("Growth rate" = Growth_Rate_day)
```

```
# Display the final data frame
all_tox_predict_Growth0 %>% 
  kable() %>% 
  kable_styling("striped") %>% 
  scroll_box(width = "100%", height = "450px")
```

Table C3

| Growth rate | Species | Treatment | Mean | LCI | HCI |
| --- | --- | --- | --- | --- | --- |
| 0 | D. sacculus | Control | 139.9773 | 118.56099 | 161.3936 |
| 0 | D. sacculus | Copepod | 151.4866 | 130.03503 | 172.9381 |
| 0 | D. sacculus | 1 nM | 168.8119 | 147.22146 | 190.4023 |
| 0 | D. sacculus | 5 nM | 180.3696 | 157.55749 | 203.1817 |
| 0 | D. sacculus | 10 nM | 203.0495 | 181.27897 | 224.8200 |
| 0 | D. acuminata | Control | 113.9805 | 93.94396 | 134.0170 |
| 0 | D. acuminata | Copepod | 136.8613 | 116.10857 | 157.6139 |
| 0 | D. acuminata | 1 nM | 135.4153 | 116.55776 | 154.2729 |
| 0 | D. acuminata | 5 nM | 135.0231 | 112.05882 | 157.9873 |
| 0 | D. acuminata | 10 nM | 137.3694 | 115.99998 | 158.7387 |

###### The plot

Using covariate adjusted means and CIs. note that the individual
replicate values are NOT covariate adjusted.

```
# Define color palette for species
palette_species <- c("black", "grey70")

# Create plot of total toxin content by treatment and species using ggplot2
data_both_species %>%
  ggplot(aes(
    x = Treatment, # x-axis: Treatment
    y = Tox_total, # y-axis: Total Toxin content
    colour = Species # color: Species
  )) +
  
  # Jittered points
  geom_jitter(
    aes(
      x = Treatment,
      y = Tox_total,
      colour = Species
    ),
    size = 2, # size of points
    alpha = 0.3, # transparency of points
    shape = 18, # shape of points
    position = position_jitterdodge(
      jitter.width = 0.3, # width of jitter
      dodge.width = 1 # width of dodge
    ),
    show.legend = FALSE # do not show legend for jittered points
  ) +
  
  # Mean (covariate adjusted)
  geom_point(
    data = all_tox_predict_Growth0, # data for mean points
    aes(
      x = Treatment, 
      y = Mean
    ),
    position = position_dodge(width = 0.4), # position of mean points
    size = 4, # size of mean points
    shape = 20, # shape of mean points
    show.legend = TRUE # show legend for mean points
  ) +
  
  # 95% CI (for covariate adjusted mean)
  geom_errorbar(
    data = all_tox_predict_Growth0, # data for error bars
    aes(
      x = Treatment, 
      y = Mean, 
      ymin = LCI, # Lower confidence limit
      ymax = HCI  # Upper confidence limit
    ),
    width = 0.25, # width of error bars
    linewidth = 0.5, # line width of error bars
    position = position_dodge(0.4), # position of error bars
    show.legend = FALSE # do not show legend for error bars
  ) +
  
  # Axes
  scale_y_continuous(
    limits = c(0, 300), # set y-axis limits
    n.breaks = 8, # set number of breaks on y-axis
    expand = c(0, 0) # remove expansion at the axis limits
  ) +
  
  # Labels
  xlab("") + # remove x-axis label
  ylab(bquote(Toxin~content~(ng~mL^-1))) + # y-axis label with mathematical annotation
  
  # Colours and theme modifications
  scale_colour_manual(
    values = palette_species # custom color palette for species
  ) +
  
  theme_classic() + # Set base theme to classic
  
  theme( # Modify theme
    legend.position = c(0.2, 0.92), # position the legend
    
    legend.title = element_blank(), # remove legend title
    
    plot.margin = unit(c(0.3, 0, -0.15, 0), "cm"), # set plot margins (top, right, bottom, left)
    
    axis.title.y = element_text( # vertical adjustment and font size for y-axis title
      vjust = 0, 
      size = 10
    ), 
    
    axis.text.y = element_text( # color and font size for y-axis text
      colour = "black", 
      size = 8
    ), 
    
    axis.text.x = element_text( # color and font size for x-axis text
      colour = "black", 
      size = 7.5
    ), 
    
    strip.background = element_rect( # remove background for strip
      linetype = "blank"
    ), 
    
    legend.background = element_rect( # transparent legend background
      fill = 'transparent', 
      colour = 'transparent'
    ), 
    
    legend.box.background = element_rect( # transparent legend panel background
      fill = 'transparent', 
      colour = 'transparent'
    ), 
    
    legend.key.size = unit(1, "mm"), # size of legend keys
    
    legend.text = element_text( # color, font size, and style for legend text
      colour = "black", 
      size = 8, 
      face = "italic"
    ) 
  ) +
  
  # Annotations
  geom_text(
    aes(
      x = 0.8, # x-position of the text
      y = 12, # y-position of the text
      label = "a", # text label
      family = "sans" # font family
    ),
    show.legend = FALSE, # do not show legend for text
    size = 3, # size of text
    colour = "black" # color of text
  ) +
  
  geom_text(
    aes(
      x = 1.8, 
      y = 12, 
      label = "a",
      family = "sans"
    ),
    show.legend = FALSE, 
    size = 3, 
    colour = "black"
  ) +
  
  geom_text(
    aes(
      x = 2.8, 
      y = 12, 
      label = "ab",
      family = "sans"
    ),
    show.legend = FALSE, 
    size = 3, 
    colour = "black"
  ) +
  
  geom_text(
    aes(
      x = 3.8, 
      y = 12, 
      label = "ab",
      family = "sans"
    ),
    show.legend = FALSE, 
    size = 3, 
    colour = "black"
  ) +
  
  geom_text(
    aes(
      x = 4.8, 
      y = 12, 
      label = "b",
      family = "sans"
    ),
    show.legend = FALSE, 
    size = 3, 
    colour = "black"
  )
```

Figure C15. Total toxin content, normalised by cell count, for
*Dinophysis sacculus* and *Dinophysis acuminata* after
68.5 hours of exposure to three concentrations of chemical cues from
copepods (copepodamides), a positive control (one live *Acartia*
sp.) or a control treatment. Circles are the covariate adjusted mean
values, predicted at growth rate = 0, of three replicates (n = 3) and
error bars show the 95% confidence intervals of the predicted means.
Transparent diamonds are the measured (raw) toxin contents of each
replicate and are NOT covariate adjusted. Based on Tukey’s Honest
Significant Differences post-hoc procedure with ⍺ set to 0.05, shared
(or lack of) letters denote statistically homogeneous treatment
subgroups.

```
#ggsave(path = "figures_manuscript", "Total_tox_corrected_Dino_mL.tiff", width = 75, height = 65, units = "mm", dpi=700)
```

The same plot, without adjusting means and CIs for the effect of
growth rate:

```
# Define color palette for species
palette_species <- c("black", "grey70")

# Create plot of total toxin content by treatment and species using ggplot2
data_both_species %>%
  ggplot(aes(
    x = Treatment, # x-axis: Treatment
    y = Tox_total, # y-axis: Total Toxin content
    colour = Species # color: Species
  )) +

  # Jittered points
  geom_jitter(
    aes(
      x = Treatment,
      y = Tox_total,
      colour = Species
    ),
    size = 2, # size of points
    alpha = 0.4, # transparency of points
    shape = 18, # shape of points
    position = position_jitterdodge(
      jitter.width = 0.5, # width of jitter
      dodge.width = 1 # width of dodge
    )
  ) +

  # Mean (not covariate adjusted)
  geom_point(
    data = desc_both_total_toxins, # data for mean points
    aes(
      x = Treatment, 
      y = Mean
    ),
    position = position_dodge(width = 0.4), # position of mean points
    size = 4, # size of mean points
    shape = 20, # shape of mean points
    show.legend = TRUE # show legend for mean points
  ) +

  # 95% Confidence Intervals
  geom_errorbar(
    data = desc_both_total_toxins, # data for error bars
    aes(
      x = Treatment, 
      y = Mean, 
      ymin = LCI, # Lower confidence limit
      ymax = HCI  # Upper confidence limit
    ),
    width = 0.25, # width of error bars
    linewidth = 0.5, # line width of error bars
    position = position_dodge(0.4), # position of error bars
    show.legend = FALSE # do not show legend for error bars
  ) +

  # Axes
  scale_y_continuous(
    limits = c(-25, 400), # set y-axis limits
    n.breaks = 9, # set number of breaks on y-axis
    expand = c(0, 0) # remove expansion at the axis limits
  ) +

  # Labels
  xlab("") + # remove x-axis label
  ylab(bquote(Toxin~content~(ng~mL^-1))) + # y-axis label with mathematical annotation

  # Colours and theme modifications
  scale_colour_manual(
    values = palette_species # custom color palette for species
  ) +
  theme_classic() +
  theme(
    legend.position = c(0.2, 0.92), # position the legend
    
    legend.title = element_blank(), # remove legend title
    
    plot.margin = unit(c(0.3, 0, -0.15, 0), "cm"), # set plot margins (top, right, bottom, left)
    
    axis.title.y = element_text( # vertical adjustment and font size for y-axis title
      vjust = 0, 
      size = 10
    ), 
    
    axis.text.y = element_text( # color and font size for y-axis text
      colour = "black", 
      size = 8
    ), 
    
    axis.text.x = element_text( # color and font size for x-axis text
      colour = "black", 
      size = 7.5
    ), 
    
    strip.background = element_rect( # remove background for strip
      linetype = "blank"
    ), 
    
    legend.background = element_rect( # transparent legend background
      fill = 'transparent', 
      colour = 'transparent'
    ), 
    
    legend.box.background = element_rect( # transparent legend panel background
      fill = 'transparent', 
      colour = 'transparent'
    ), 
    
    legend.key.size = unit(1, "mm"), # size of legend keys
    
    legend.text = element_text( # color, font size, and style for legend text
      colour = "black", 
      size = 8, 
      face = "italic"
    ) 
  ) +

  # Annotations
  geom_text(
    aes(
      x = 0.8, # x-position of the text
      y = -8, # y-position of the text
      label = "a", # text label
      family = "sans" # font family
    ),
    show.legend = FALSE, # do not show legend for text
    size = 3, # size of text
    colour = "black" # color of text
  ) +
  geom_text(
    aes(
      x = 1.8, 
      y = -8, 
      label = "a",
      family = "sans"
    ),
    show.legend = FALSE, 
    size = 3, 
    colour = "black"
  ) +
  geom_text(
    aes(
      x = 2.8, 
      y = -8, 
      label = "ab",
      family = "sans"
    ),
    show.legend = FALSE, 
    size = 3, 
    colour = "black"
  ) +
  geom_text(
    aes(
      x = 3.8, 
      y = -8, 
      label = "ab",
      family = "sans"
    ),
    show.legend = FALSE, 
    size = 3, 
    colour = "black"
  ) +
  geom_text(
    aes(
      x = 4.8, 
      y = -8, 
      label = "b",
      family = "sans"
    ),
    show.legend = FALSE, 
    size = 3, 
    colour = "black"
  )
```

Figure C16. Total toxin content, normalised by cell count, for
*Dinophysis sacculus* and *Dinophysis acuminata* after
68.5 hours of exposure to three concentrations of chemical cues from
copepods (copepodamides), a positive control (one live *Acartia*
sp.) or a control treatment. Circles are the mean values of three
replicates (n=3, transparent diamonds) and error bars show the 95%
confidence intervals of the means. Based on Tukey’s Honest Significant
Differences post-hoc procedure with ⍺ set to 0.05, shared (or lack of)
letters denote statistically homogeneous treatment subgroups.

```
#ggsave(path = "figures_manuscript", "Total_tox_raw_Dino.tiff", width = 75, height = 65, units = "mm", dpi=700)
```

##### Plot of effect sizes

Plot of covariate adjusted effect sizes as percentage increase in
total toxins, back-transformed from the standard log response ratio

###### Data wrangling

```
# Calculate summary statistics for all_tox_predict_Growth0
summary_stats_all <- all_tox_predict_Growth0 %>%
  mutate(
    CI = HCI - LCI, # Confidence Interval
    n = 3, # Sample size
    SE = CI / qt(0.975, df = n - 1), # Standard Error
    SD = SE * sqrt(n) # Standard Deviation
  ) %>%
  tibble() # Convert to tibble

# Round the Mean and SD columns to 2 decimal places
summary_stats_all$Mean <- round(summary_stats_all$Mean, digits = 2)
summary_stats_all$SD <- round(summary_stats_all$SD, digits = 2)
```

```
# Display the summary statistics in a styled table with scrolling
summary_stats_all %>%
  kable() %>%
  kable_styling("striped") %>%
  scroll_box(
    width = "100%", # set width of the scroll box
    height = "450px" # set height of the scroll box
  )
```

Table C4

| Growth rate | Species | Treatment | Mean | LCI | HCI | CI | n | SE | SD |
| --- | --- | --- | --- | --- | --- | --- | --- | --- | --- |
| 0 | D. sacculus | Control | 139.98 | 118.56099 | 161.3936 | 42.83260 | 3 | 9.954930 | 17.24 |
| 0 | D. sacculus | Copepod | 151.49 | 130.03503 | 172.9381 | 42.90311 | 3 | 9.971315 | 17.27 |
| 0 | D. sacculus | 1 nM | 168.81 | 147.22146 | 190.4023 | 43.18083 | 3 | 10.035862 | 17.38 |
| 0 | D. sacculus | 5 nM | 180.37 | 157.55749 | 203.1817 | 45.62416 | 3 | 10.603729 | 18.37 |
| 0 | D. sacculus | 10 nM | 203.05 | 181.27897 | 224.8200 | 43.54099 | 3 | 10.119568 | 17.53 |
| 0 | D. acuminata | Control | 113.98 | 93.94396 | 134.0170 | 40.07308 | 3 | 9.313576 | 16.13 |
| 0 | D. acuminata | Copepod | 136.86 | 116.10857 | 157.6139 | 41.50538 | 3 | 9.646462 | 16.71 |
| 0 | D. acuminata | 1 nM | 135.42 | 116.55776 | 154.2729 | 37.71514 | 3 | 8.765555 | 15.18 |
| 0 | D. acuminata | 5 nM | 135.02 | 112.05882 | 157.9873 | 45.92846 | 3 | 10.674451 | 18.49 |
| 0 | D. acuminata | 10 nM | 137.37 | 115.99998 | 158.7387 | 42.73876 | 3 | 9.933118 | 17.20 |

```
# Create a new dataframe for contrasts to calculate effect size for each treatment relative to control
effects_df <- data.frame(
  Species = c(
    rep("D. sacculus", 4), # Species: D. sacculus for 4 treatments
    rep("D. acuminata", 4) # Species: D. acuminata for 4 treatments
  ),
  Treatment = c(
    rep(c("Copepod", "1 nM", "5 nM", "10 nM"), 2) # Treatments for both species
  ),
  Mean_cont = c(
    rep(139.98, 4), # Mean of control for D. sacculus
    rep(113.98, 4) # Mean of control for D. acuminata
  ),
  Mean_treat = c(
    151.49, 168.81, 180.37, 203.05, # Mean of treatments for D. sacculus
    136.86, 135.42, 135.02, 137.37 # Mean of treatments for D. acuminata
  ),
  SD_cont = c(
    rep(17.24, 4), # Standard deviation of control for D. sacculus
    rep(16.13, 4) # Standard deviation of control for D. acuminata
  ),
  SD_treat = c(
    17.27, 17.38, 18.37, 17.53, # Standard deviation of treatments for D. sacculus
    16.71, 15.18, 18.49, 17.20 # Standard deviation of treatments for D. acuminata
  ),
  N_cont = c(rep(3, 4)), # Sample size of control for both species
  N_treat = c(rep(3, 4)) # Sample size of treatments for both species
)
```

```
# Display the effects_df dataframe in a styled table with scrolling
effects_df %>%
  kable() %>%
  kable_styling("striped") %>%
  scroll_box(
    width = "100%", # set width of the scroll box
    height = "400px" # set height of the scroll box
  )
```

Table C5

| Species | Treatment | Mean\_cont | Mean\_treat | SD\_cont | SD\_treat | N\_cont | N\_treat |
| --- | --- | --- | --- | --- | --- | --- | --- |
| D. sacculus | Copepod | 139.98 | 151.49 | 17.24 | 17.27 | 3 | 3 |
| D. sacculus | 1 nM | 139.98 | 168.81 | 17.24 | 17.38 | 3 | 3 |
| D. sacculus | 5 nM | 139.98 | 180.37 | 17.24 | 18.37 | 3 | 3 |
| D. sacculus | 10 nM | 139.98 | 203.05 | 17.24 | 17.53 | 3 | 3 |
| D. acuminata | Copepod | 113.98 | 136.86 | 16.13 | 16.71 | 3 | 3 |
| D. acuminata | 1 nM | 113.98 | 135.42 | 16.13 | 15.18 | 3 | 3 |
| D. acuminata | 5 nM | 113.98 | 135.02 | 16.13 | 18.49 | 3 | 3 |
| D. acuminata | 10 nM | 113.98 | 137.37 | 16.13 | 17.20 | 3 | 3 |

###### Effect size calculation

```
# Custom function to calculate SE of the log response ratio 
# Reference: Hedges, L. V., Gurevitch, J., & Curtis, P. S. (1999). 
# The meta-analysis of response ratios in experimental ecology. Ecology, 80(4), 1150-1156.
SE_rr <- function(mean_treatment, mean_control, 
                  sd_treatment, sd_control, 
                  n_treatment, n_control) {
  result <- sqrt(
    (sd_treatment^2 / (n_treatment * mean_treatment^2)) + # SE component for treatment
    (sd_control^2 / (n_control * mean_control^2)) # SE component for control
  )
  return(result)
}

# Calculate effect sizes (Log response ratio) and their 95% CI
effects_summary <- 
  effects_df %>% 
  mutate(
    RR = log(Mean_treat / Mean_cont), # Log response ratio
    Effect_percent = 100 * (exp(RR) - 1), # Effect size in percentage
    SE_RR = SE_rr(Mean_treat, Mean_cont, SD_treat, SD_cont, N_treat, N_cont), # Standard error of log response ratio
    CI = SE_RR * qt(0.975, 3), # One-sided confidence interval
    LCI = RR - CI, # Lower confidence limit
    HCI = RR + CI, # Upper confidence limit
    LCI_percent = 100 * (exp(LCI) - 1), # Lower confidence interval in percentage
    HCI_percent = 100 * (exp(HCI) - 1) # Upper confidence interval in percentage
  ) %>% 
  tibble()

# Set order of levels in factor "Treatment"
effects_summary$Treatment <- factor(
  effects_summary$Treatment,
  levels = c('Copepod', '1 nM', '5 nM', '10 nM')
)

# Set order of levels in factor Species
effects_summary$Species <- factor(
  effects_summary$Species, 
  levels = c('D. sacculus', 'D. acuminata')
)
```

```
# Display the effects_summary dataframe in a styled table with scrolling
effects_summary %>%
  kable() %>%
  kable_styling("striped") %>%
  scroll_box(
    width = "100%", # set width of the scroll box
    height = "450px" # set height of the scroll box
  )
```

Table C6

| Species | Treatment | Mean\_cont | Mean\_treat | SD\_cont | SD\_treat | N\_cont | N\_treat | RR | Effect\_percent | SE\_RR | CI | LCI | HCI | LCI\_percent | HCI\_percent |
| --- | --- | --- | --- | --- | --- | --- | --- | --- | --- | --- | --- | --- | --- | --- | --- |
| D. sacculus | Copepod | 139.98 | 151.49 | 17.24 | 17.27 | 3 | 3 | 0.0790201 | 8.222603 | 0.0968929 | 0.3083565 | -0.2293364 | 0.3873765 | -20.493899 | 47.31111 |
| D. sacculus | 1 nM | 139.98 | 168.81 | 17.24 | 17.38 | 3 | 3 | 0.1872743 | 20.595799 | 0.0926794 | 0.2949473 | -0.1076731 | 0.4822216 | -10.207888 | 61.96686 |
| D. sacculus | 5 nM | 139.98 | 180.37 | 17.24 | 18.37 | 3 | 3 | 0.2535107 | 28.854122 | 0.0922698 | 0.2936437 | -0.0401329 | 0.5471544 | -3.933827 | 72.83279 |
| D. sacculus | 10 nM | 139.98 | 203.05 | 17.24 | 17.53 | 3 | 3 | 0.3719527 | 45.056437 | 0.0868369 | 0.2763539 | 0.0955988 | 0.6483066 | 10.031757 | 91.22997 |
| D. acuminata | Copepod | 113.98 | 136.86 | 16.13 | 16.71 | 3 | 3 | 0.1829355 | 20.073697 | 0.1079106 | 0.3434198 | -0.1604843 | 0.5263553 | -14.826883 | 69.27516 |
| D. acuminata | 1 nM | 113.98 | 135.42 | 16.13 | 15.18 | 3 | 3 | 0.1723581 | 18.810318 | 0.1042309 | 0.3317092 | -0.1593512 | 0.5040673 | -14.730314 | 65.54408 |
| D. acuminata | 5 nM | 113.98 | 135.02 | 16.13 | 18.49 | 3 | 3 | 0.1693999 | 18.459379 | 0.1136956 | 0.3618302 | -0.1924303 | 0.5312302 | -17.504820 | 70.10236 |
| D. acuminata | 10 nM | 113.98 | 137.37 | 16.13 | 17.20 | 3 | 3 | 0.1866550 | 20.521144 | 0.1090935 | 0.3471841 | -0.1605291 | 0.5338392 | -14.830697 | 70.54673 |

###### The plot

```
# Define color palette for species
palette_species <- c("black", "grey70")

# Plot effect sizes (log response ratio) and their 95% CI
effects_summary %>%
  ggplot(aes(
    x = Treatment, # x-axis: Treatment
    y = Effect_percent, # y-axis: Effect size in percentage
    colour = Species # color: Species
  )) +
  
  geom_hline(
    yintercept = 0, # horizontal line at y = 0
    alpha = 0.5, # transparency of the line
    linetype = "dotted" # line type
  ) +

  # Mean (covariate adjusted)
  geom_point(
    position = position_dodge(width = 0.4), # position of mean points
    size = 4, # size of mean points
    shape = 20, # shape of mean points
    show.legend = TRUE # show legend for mean points
  ) +

  # 95% CI (for covariate adjusted mean)
  geom_errorbar(
    aes(
      x = Treatment, 
      y = Effect_percent, 
      ymin = LCI_percent, # Lower confidence interval in percentage
      ymax = HCI_percent # Upper confidence interval in percentage
    ),
    width = 0.25, # width of error bars
    linewidth = 0.5, # line width of error bars
    position = position_dodge(0.4), # position of error bars
    show.legend = FALSE # do not show legend for error bars
  ) +

  # Axes
  scale_y_continuous(
    limits = c(-25, 100), # set y-axis limits
    breaks = c(-25, 0, 25, 50, 75, 100), # set y-axis breaks
    expand = c(0, 0) # remove expansion at the axis limits
  ) +

  # Labels
  xlab("") + # remove x-axis label
  ylab("Toxin increase (%)") + # y-axis label

  # Colours and theme modifications
  scale_colour_manual(
    values = palette_species # custom color palette for species
  ) +
  theme_classic() +
  theme(
    legend.position = c(0.2, 0.92), # position the legend
    
    legend.title = element_blank(), # remove legend title
    
    plot.margin = unit(c(0.3, 0, -0.15, 0), "cm"), # set plot margins (top, right, bottom, left)
    
    axis.title.y = element_text( # vertical adjustment and font size for y-axis title
      vjust = -1, 
      size = 10
    ), 
    
    axis.text.y = element_text( # color and font size for y-axis text
      colour = "black", 
      size = 8
    ), 
    
    axis.text.x = element_text( # color and font size for x-axis text
      colour = "black", 
      size = 7.5
    ), 
    
    strip.background = element_rect( # remove background for strip
      linetype = "blank"
    ), 
    
    legend.background = element_rect( # transparent legend background
      fill = 'transparent', 
      colour = 'transparent'
    ), 
    
    legend.box.background = element_rect( # transparent legend panel background
      fill = 'transparent', 
      colour = 'transparent'
    ), 
    
    legend.key.size = unit(1, "mm"), # size of legend keys
    
    legend.text = element_text( # color, font size, and style for legend text
      colour = "black", 
      size = 8, 
      face = "italic"
    ) 
  ) +

  # Annotations
  geom_text(
    aes(
      x = 0.75, # x-position of the text
      y = -21, # y-position of the text
      label = "a", # text label
      family = "sans" # font family
    ),
    show.legend = FALSE, # do not show legend for text
    size = 3, # size of text
    colour = "grey50" # color of text
  ) +
  geom_text(
    aes(
      x = 1.75, 
      y = -21, 
      label = "a",
      family = "sans"
    ),
    show.legend = FALSE, 
    size = 3, 
    colour = "grey50"
  ) +
  geom_text(
    aes(
      x = 2.75, 
      y = -21, 
      label = "a",
      family = "sans"
    ),
    show.legend = FALSE, 
    size = 3, 
    colour = "grey50"
  ) +
  geom_text(
    aes(
      x = 3.75, 
      y = -21, 
      label = "b",
      family = "sans"
    ),
    show.legend = FALSE, 
    size = 3, 
    colour = "grey50"
  )
```

Figure C17.

```
#ggsave(path = "figures_manuscript", "Effect_sizes.png", width = 75, height = 65, units = "mm", dpi=700)
```

#### Analysis of intracellular and extracellular toxins

##### Exploratory data analysis and visualisation

###### Summary statistics tables

###### Intracellular

```
# Calculate descriptive statistics for total intracellular toxin content and proportion of intracellular toxins
desc_mean_ci_intra <- data_both_species %>%
  dplyr::select(
    Species, # select species
    Treatment, # select treatment
    Tox_intra_total, # select total intracellular toxin content
    prop_intra # select proportion of intracellular toxins
  ) %>%
  dplyr::group_by(Species, Treatment) %>% # group by species and treatment
  dplyr::summarise(
    N = length(Tox_intra_total), # sample size
    Mean = mean(Tox_intra_total), # mean total intracellular toxin content
    St.Dev = sd(Tox_intra_total), # standard deviation of total intracellular toxin content
    SEM = St.Dev / sqrt(N), # standard error of the mean
    CI = qt(0.975, df = N - 1) * SEM, # 95% confidence interval
    LCI = Mean - CI, # lower confidence interval
    HCI = Mean + CI, # upper confidence interval

    Mean_prop = mean(prop_intra), # mean proportion of intracellular toxins
    St.Dev_prop = sd(prop_intra), # standard deviation of proportion of intracellular toxins
    SEM_prop = St.Dev_prop / sqrt(N), # standard error of the mean for proportion
    CI_prop = qt(0.975, df = N - 1) * SEM_prop, # 95% confidence interval for proportion
    LCI_prop = Mean_prop - CI_prop, # lower confidence interval for proportion
    HCI_prop = Mean_prop + CI_prop # upper confidence interval for proportion
  ) %>%
  data.frame() %>% # convert to data frame
  mutate(Source = "Intracellular") # add source column with value "Intracellular"
```

###### Extracellular

```
# Calculate descriptive statistics for total extracellular toxin content and proportion of extracellular toxins
desc_mean_ci_extra <- data_both_species %>%
  dplyr::select(
    Species, # select species
    Treatment, # select treatment
    Tox_extra_total, # select total extracellular toxin content
    prop_extra # select proportion of extracellular toxins
  ) %>%
  dplyr::group_by(Species, Treatment) %>% # group by species and treatment
  dplyr::summarise(
    N = length(Tox_extra_total), # sample size
    Mean = mean(Tox_extra_total), # mean total extracellular toxin content
    St.Dev = sd(Tox_extra_total), # standard deviation of total extracellular toxin content
    SEM = St.Dev / sqrt(N), # standard error of the mean
    CI = qt(0.975, df = N - 1) * SEM, # 95% confidence interval
    LCI = Mean - CI, # lower confidence interval
    HCI = Mean + CI, # upper confidence interval

    Mean_prop = mean(prop_extra), # mean proportion of extracellular toxins
    St.Dev_prop = sd(prop_extra), # standard deviation of proportion of extracellular toxins
    SEM_prop = St.Dev_prop / sqrt(N), # standard error of the mean for proportion
    CI_prop = qt(0.975, df = N - 1) * SEM_prop, # 95% confidence interval for proportion
    LCI_prop = Mean_prop - CI_prop, # lower confidence interval for proportion
    HCI_prop = Mean_prop + CI_prop # upper confidence interval for proportion
  ) %>%
  data.frame() %>% # convert to data frame
  mutate(Source = "Extracellular") # add source column with value "Extracellular"
```

###### Total

This is needed for combining the datasets further down the
pipeline

```
desc_mean_ci_total <- 
    
  data_both_species %>% 
  dplyr::select(Species, Treatment, Tox_total) %>% 
  dplyr::group_by(Species, Treatment) %>% 
  dplyr::summarise(N = length(Tox_total),
                   Mean = mean(Tox_total),
                   St.Dev = sd(Tox_total),
                   SEM =  St.Dev/sqrt(N),
                   CI = qt(0.975, df = N-1)*SEM,
                   LCI = Mean - CI,
                   HCI = Mean + CI) %>% 
  data.frame() %>% 
  mutate(Mean_prop = 1,
         St.Dev_prop = 0,
         SEM_prop =  0,
         CI_prop = 0,
         LCI_prop = Mean_prop - CI_prop,
         HCI_prop = Mean_prop + CI_prop,
         Source = "Total")

# Calculate descriptive statistics for total toxin content
desc_mean_ci_total <- data_both_species %>%
  dplyr::select(
    Species, # select species
    Treatment, # select treatment
    Tox_total # select total toxin content
  ) %>%
  dplyr::group_by(Species, Treatment) %>% # group by species and treatment
  dplyr::summarise(
    N = length(Tox_total), # sample size
    Mean = mean(Tox_total), # mean total toxin content
    St.Dev = sd(Tox_total), # standard deviation of total toxin content
    SEM = St.Dev / sqrt(N), # standard error of the mean
    CI = qt(0.975, df = N - 1) * SEM, # 95% confidence interval
    LCI = Mean - CI, # lower confidence interval
    HCI = Mean + CI # upper confidence interval
  ) %>%
  data.frame() %>% # convert to data frame
  mutate(
    Mean_prop = 1, # mean proportion (set to 1 for total)
    St.Dev_prop = 0, # standard deviation of proportion (set to 0 for total)
    SEM_prop = 0, # standard error of the mean for proportion (set to 0 for total)
    CI_prop = 0, # confidence interval for proportion (set to 0 for total)
    LCI_prop = Mean_prop - CI_prop, # lower confidence interval for proportion
    HCI_prop = Mean_prop + CI_prop, # upper confidence interval for proportion
    Source = "Total" # add source column with value "Total"
  )
```

###### Combining data frames

Here we combine the three data frames

```
# Combine the descriptive statistics  tables for intracellular, extracellular, and total toxin content
desc_mean_ci_all <- bind_rows(
  desc_mean_ci_intra, 
  desc_mean_ci_extra, 
  desc_mean_ci_total
)

# Display the combined data frame in a styled table with scrolling
desc_mean_ci_all %>%
  kable() %>%
  kable_styling("striped") %>%
  scroll_box(
    width = "100%", # set width of the scroll box
    height = "450px" # set height of the scroll box
  )
```

Table C7.

| Species | Treatment | N | Mean | St.Dev | SEM | CI | LCI | HCI | Mean\_prop | St.Dev\_prop | SEM\_prop | CI\_prop | LCI\_prop | HCI\_prop | Source |
| --- | --- | --- | --- | --- | --- | --- | --- | --- | --- | --- | --- | --- | --- | --- | --- |
| D. sacculus | Control | 3 | 121.517471 | 21.1426043 | 12.2066883 | 52.5211407 | 68.996331 | 174.038612 | 0.8290626 | 0.0095481 | 0.0055126 | 0.0237187 | 0.8053440 | 0.8527813 | Intracellular |
| D. sacculus | Copepod | 3 | 141.562114 | 11.6806090 | 6.7438027 | 29.0162412 | 112.545873 | 170.578355 | 0.8858664 | 0.0201363 | 0.0116257 | 0.0500214 | 0.8358450 | 0.9358879 | Intracellular |
| D. sacculus | 1 nM | 3 | 153.203435 | 64.0347267 | 36.9704667 | 159.0710795 | -5.867645 | 312.274514 | 0.8367508 | 0.0215536 | 0.0124440 | 0.0535420 | 0.7832087 | 0.8902928 | Intracellular |
| D. sacculus | 5 nM | 3 | 89.239199 | 14.8341717 | 8.5645131 | 36.8501255 | 52.389073 | 126.089324 | 0.6054254 | 0.0292642 | 0.0168957 | 0.0726962 | 0.5327292 | 0.6781216 | Intracellular |
| D. sacculus | 10 nM | 3 | 90.319326 | 70.9749369 | 40.9773989 | 176.3115173 | -85.992192 | 266.630843 | 0.4417397 | 0.1546261 | 0.0892734 | 0.3841125 | 0.0576272 | 0.8258522 | Intracellular |
| D. acuminata | Control | 3 | 65.536808 | 14.6974161 | 8.4855571 | 36.5104056 | 29.026402 | 102.047214 | 0.5350056 | 0.0943314 | 0.0544622 | 0.2343321 | 0.3006735 | 0.7693377 | Intracellular |
| D. acuminata | Copepod | 3 | 66.240425 | 11.0134281 | 6.3586057 | 27.3588720 | 38.881553 | 93.599297 | 0.4546767 | 0.0574479 | 0.0331676 | 0.1427085 | 0.3119682 | 0.5973851 | Intracellular |
| D. acuminata | 1 nM | 3 | 46.986948 | 0.2407571 | 0.1390012 | 0.5980737 | 46.388874 | 47.585022 | 0.3532064 | 0.0418737 | 0.0241758 | 0.1040201 | 0.2491862 | 0.4572265 | Intracellular |
| D. acuminata | 5 nM | 3 | 11.243464 | 0.9383444 | 0.5417534 | 2.3309766 | 8.912487 | 13.574441 | 0.0949457 | 0.0147031 | 0.0084888 | 0.0365245 | 0.0584212 | 0.1314702 | Intracellular |
| D. acuminata | 10 nM | 3 | 7.500185 | 0.5244191 | 0.3027735 | 1.3027293 | 6.197456 | 8.802915 | 0.0604320 | 0.0102190 | 0.0059000 | 0.0253854 | 0.0350466 | 0.0858175 | Intracellular |
| D. sacculus | Control | 3 | 25.205382 | 5.8473956 | 3.3759954 | 14.5257360 | 10.679646 | 39.731118 | 0.1709374 | 0.0095481 | 0.0055126 | 0.0237187 | 0.1472187 | 0.1946560 | Extracellular |
| D. sacculus | Copepod | 3 | 18.402814 | 4.6165307 | 2.6653553 | 11.4680981 | 6.934716 | 29.870912 | 0.1141336 | 0.0201363 | 0.0116257 | 0.0500214 | 0.0641121 | 0.1641550 | Extracellular |
| D. sacculus | 1 nM | 3 | 28.883166 | 7.9982216 | 4.6177754 | 19.8686838 | 9.014482 | 48.751849 | 0.1632492 | 0.0215536 | 0.0124440 | 0.0535420 | 0.1097072 | 0.2167913 | Extracellular |
| D. sacculus | 5 nM | 3 | 57.616878 | 3.4735536 | 2.0054571 | 8.6287854 | 48.988093 | 66.245663 | 0.3945746 | 0.0292642 | 0.0168957 | 0.0726962 | 0.3218784 | 0.4672708 | Extracellular |
| D. sacculus | 10 nM | 3 | 95.051711 | 12.0234433 | 6.9417382 | 29.8678888 | 65.183822 | 124.919600 | 0.5582603 | 0.1546261 | 0.0892734 | 0.3841125 | 0.1741478 | 0.9423728 | Extracellular |
| D. acuminata | Control | 3 | 56.551420 | 11.8951400 | 6.8676623 | 29.5491658 | 27.002254 | 86.100586 | 0.4649944 | 0.0943314 | 0.0544622 | 0.2343321 | 0.2306623 | 0.6993265 | Extracellular |
| D. acuminata | Copepod | 3 | 80.954162 | 23.5477877 | 13.5953215 | 58.4959473 | 22.458215 | 139.450110 | 0.5453233 | 0.0574479 | 0.0331676 | 0.1427085 | 0.4026149 | 0.6880318 | Extracellular |
| D. acuminata | 1 nM | 3 | 87.284969 | 15.9774273 | 9.2245719 | 39.6901296 | 47.594839 | 126.975098 | 0.6467936 | 0.0418737 | 0.0241758 | 0.1040201 | 0.5427735 | 0.7508138 | Extracellular |
| D. acuminata | 5 nM | 3 | 108.203370 | 11.5289405 | 6.6562369 | 28.6394760 | 79.563894 | 136.842846 | 0.9050543 | 0.0147031 | 0.0084888 | 0.0365245 | 0.8685298 | 0.9415788 | Extracellular |
| D. acuminata | 10 nM | 3 | 117.899416 | 11.9829954 | 6.9183856 | 29.7674107 | 88.132005 | 147.666826 | 0.9395680 | 0.0102190 | 0.0059000 | 0.0253854 | 0.9141825 | 0.9649534 | Extracellular |
| D. sacculus | Control | 3 | 146.722853 | 26.8830386 | 15.5209296 | 66.7811701 | 79.941683 | 213.504023 | 1.0000000 | 0.0000000 | 0.0000000 | 0.0000000 | 1.0000000 | 1.0000000 | Total |
| D. sacculus | Copepod | 3 | 159.964927 | 15.5692601 | 8.9889165 | 38.6761862 | 121.288741 | 198.641114 | 1.0000000 | 0.0000000 | 0.0000000 | 0.0000000 | 1.0000000 | 1.0000000 | Total |
| D. sacculus | 1 nM | 3 | 182.086600 | 71.8209744 | 41.4658589 | 178.4131909 | 3.673409 | 360.499791 | 1.0000000 | 0.0000000 | 0.0000000 | 0.0000000 | 1.0000000 | 1.0000000 | Total |
| D. sacculus | 5 nM | 3 | 146.856077 | 18.0198561 | 10.4037687 | 44.7638040 | 102.092273 | 191.619881 | 1.0000000 | 0.0000000 | 0.0000000 | 0.0000000 | 1.0000000 | 1.0000000 | Total |
| D. sacculus | 10 nM | 3 | 185.371036 | 81.8335472 | 47.2466205 | 203.2858008 | -17.914764 | 388.656837 | 1.0000000 | 0.0000000 | 0.0000000 | 0.0000000 | 1.0000000 | 1.0000000 | Total |
| D. acuminata | Control | 3 | 122.088228 | 13.0249190 | 7.5199405 | 32.3556925 | 89.732535 | 154.443921 | 1.0000000 | 0.0000000 | 0.0000000 | 0.0000000 | 1.0000000 | 1.0000000 | Total |
| D. acuminata | Copepod | 3 | 147.194587 | 31.8878269 | 18.4104455 | 79.2137534 | 67.980834 | 226.408340 | 1.0000000 | 0.0000000 | 0.0000000 | 0.0000000 | 1.0000000 | 1.0000000 | Total |
| D. acuminata | 1 nM | 3 | 134.271917 | 15.7391390 | 9.0869961 | 39.0981887 | 95.173728 | 173.370105 | 1.0000000 | 0.0000000 | 0.0000000 | 0.0000000 | 1.0000000 | 1.0000000 | Total |
| D. acuminata | 5 nM | 3 | 119.446834 | 10.8449162 | 6.2613153 | 26.9402653 | 92.506568 | 146.387099 | 1.0000000 | 0.0000000 | 0.0000000 | 0.0000000 | 1.0000000 | 1.0000000 | Total |
| D. acuminata | 10 nM | 3 | 125.399601 | 11.4609081 | 6.6169584 | 28.4704741 | 96.929127 | 153.870075 | 1.0000000 | 0.0000000 | 0.0000000 | 0.0000000 | 1.0000000 | 1.0000000 | Total |

```
# Select and round the relevant columns
desc_mean_ci_all <- desc_mean_ci_all %>%
  dplyr::select(
    Species, 
    Treatment, 
    Source, 
    N,
    Mean, 
    LCI, 
    HCI, 
    Mean_prop, 
    LCI_prop, 
    HCI_prop
  ) %>%
  mutate(across(
    where(is.numeric), 
    round, 
    digits = 3 # round numeric variables to 3 digits
  ))
```

```
# Display the final rounded data frame in a styled table with scrolling
desc_mean_ci_all %>%
  kable() %>%
  kable_styling("striped") %>%
  scroll_box(
    width = "100%", # set width of the scroll box
    height = "450px" # set height of the scroll box
  )
```

Table C8.

| Species | Treatment | Source | N | Mean | LCI | HCI | Mean\_prop | LCI\_prop | HCI\_prop |
| --- | --- | --- | --- | --- | --- | --- | --- | --- | --- |
| D. sacculus | Control | Intracellular | 3 | 121.517 | 68.996 | 174.039 | 0.829 | 0.805 | 0.853 |
| D. sacculus | Copepod | Intracellular | 3 | 141.562 | 112.546 | 170.578 | 0.886 | 0.836 | 0.936 |
| D. sacculus | 1 nM | Intracellular | 3 | 153.203 | -5.868 | 312.275 | 0.837 | 0.783 | 0.890 |
| D. sacculus | 5 nM | Intracellular | 3 | 89.239 | 52.389 | 126.089 | 0.605 | 0.533 | 0.678 |
| D. sacculus | 10 nM | Intracellular | 3 | 90.319 | -85.992 | 266.631 | 0.442 | 0.058 | 0.826 |
| D. acuminata | Control | Intracellular | 3 | 65.537 | 29.026 | 102.047 | 0.535 | 0.301 | 0.769 |
| D. acuminata | Copepod | Intracellular | 3 | 66.240 | 38.882 | 93.599 | 0.455 | 0.312 | 0.597 |
| D. acuminata | 1 nM | Intracellular | 3 | 46.987 | 46.389 | 47.585 | 0.353 | 0.249 | 0.457 |
| D. acuminata | 5 nM | Intracellular | 3 | 11.243 | 8.912 | 13.574 | 0.095 | 0.058 | 0.131 |
| D. acuminata | 10 nM | Intracellular | 3 | 7.500 | 6.197 | 8.803 | 0.060 | 0.035 | 0.086 |
| D. sacculus | Control | Extracellular | 3 | 25.205 | 10.680 | 39.731 | 0.171 | 0.147 | 0.195 |
| D. sacculus | Copepod | Extracellular | 3 | 18.403 | 6.935 | 29.871 | 0.114 | 0.064 | 0.164 |
| D. sacculus | 1 nM | Extracellular | 3 | 28.883 | 9.014 | 48.752 | 0.163 | 0.110 | 0.217 |
| D. sacculus | 5 nM | Extracellular | 3 | 57.617 | 48.988 | 66.246 | 0.395 | 0.322 | 0.467 |
| D. sacculus | 10 nM | Extracellular | 3 | 95.052 | 65.184 | 124.920 | 0.558 | 0.174 | 0.942 |
| D. acuminata | Control | Extracellular | 3 | 56.551 | 27.002 | 86.101 | 0.465 | 0.231 | 0.699 |
| D. acuminata | Copepod | Extracellular | 3 | 80.954 | 22.458 | 139.450 | 0.545 | 0.403 | 0.688 |
| D. acuminata | 1 nM | Extracellular | 3 | 87.285 | 47.595 | 126.975 | 0.647 | 0.543 | 0.751 |
| D. acuminata | 5 nM | Extracellular | 3 | 108.203 | 79.564 | 136.843 | 0.905 | 0.869 | 0.942 |
| D. acuminata | 10 nM | Extracellular | 3 | 117.899 | 88.132 | 147.667 | 0.940 | 0.914 | 0.965 |
| D. sacculus | Control | Total | 3 | 146.723 | 79.942 | 213.504 | 1.000 | 1.000 | 1.000 |
| D. sacculus | Copepod | Total | 3 | 159.965 | 121.289 | 198.641 | 1.000 | 1.000 | 1.000 |
| D. sacculus | 1 nM | Total | 3 | 182.087 | 3.673 | 360.500 | 1.000 | 1.000 | 1.000 |
| D. sacculus | 5 nM | Total | 3 | 146.856 | 102.092 | 191.620 | 1.000 | 1.000 | 1.000 |
| D. sacculus | 10 nM | Total | 3 | 185.371 | -17.915 | 388.657 | 1.000 | 1.000 | 1.000 |
| D. acuminata | Control | Total | 3 | 122.088 | 89.733 | 154.444 | 1.000 | 1.000 | 1.000 |
| D. acuminata | Copepod | Total | 3 | 147.195 | 67.981 | 226.408 | 1.000 | 1.000 | 1.000 |
| D. acuminata | 1 nM | Total | 3 | 134.272 | 95.174 | 173.370 | 1.000 | 1.000 | 1.000 |
| D. acuminata | 5 nM | Total | 3 | 119.447 | 92.507 | 146.387 | 1.000 | 1.000 | 1.000 |
| D. acuminata | 10 nM | Total | 3 | 125.400 | 96.929 | 153.870 | 1.000 | 1.000 | 1.000 |

###### Visual data exploration

###### Intracellular toxins

```
# Grouped by treatment, coloured by species
data_both_species %>% 
  ggplot(aes(
    x = Treatment, # x-axis: Treatment
    y = Tox_intra_total, # y-axis: Total intracellular toxins
    fill = Species # fill color: Species
  )) + 

  # Configure y-axis
  scale_y_continuous(
    limits = c(0, 250), # set y-axis limits
    n.breaks = 11, # number of breaks on y-axis
    expand = c(0, 0) # remove expansion at the axis limits
  ) +
  
  # Boxplot
  geom_boxplot(
    position = position_dodge(1) # position dodge for boxplots
  ) +
  
  # Points
  geom_point(
    position = position_dodge(1), # position dodge for points
    show.legend = FALSE # do not show legend for points
  ) +
  
  # Theme
  theme_classic() +
  theme(
    legend.position = "top" # position the legend at the top
  ) +
  
  # Fill color palette
  scale_fill_brewer(
    palette = "Dark2" # use the "Dark2" palette for fill colors
  ) +
  
  # Y-axis label
  ylab(
    bquote(Intracellular~toxins~(ng~mL^-1)) # y-axis label with mathematical annotation
  )
```

Figure C18.

```
# Grouped by species, coloured by treatment
data_both_species %>% 
  ggplot(aes(
    x = Species, # x-axis: Species
    y = Tox_intra_total, # y-axis: Total intracellular toxins
    fill = Treatment # fill color: Treatment
  )) +
  
  # Configure y-axis
  scale_y_continuous(
    limits = c(0, 250), # set y-axis limits
    n.breaks = 11, # number of breaks on y-axis
    expand = c(0, 0) # remove expansion at the axis limits
  ) +
  
  # Boxplot
  geom_boxplot(
    position = position_dodge(1) # position dodge for boxplots
  ) +
  
  # Points
  geom_point(
    position = position_dodge(1), # position dodge for points
    show.legend = FALSE # do not show legend for points
  ) +
  
  # Theme
  theme_classic() +
  theme(
    legend.position = "top" # position the legend at the top
  ) +
  
  scale_fill_manual(
    values = custom_wes_palette # use custom "Moonrise3" palette for fill colors
  ) +
  
  # Y-axis label
  ylab(
    bquote(Intracellular~toxins~(ng~mL^-1)) # y-axis label with mathematical annotation
  )
```

Figure C19.

We see the same approximate trend across treatments in both species,
which indicates that there is no interaction effect between the two
factors. However, *D. acuminata* appears to have considerably
less intracellular toxins than *D. sacculus* across all treatment
levels.

Grouped bar chart to visualise this with means and their 95%
confidence intervals.

```
# Plot intracellular toxin content grouped by treatment and colored by species
data_both_species %>%
  ggplot(aes(
    x = Treatment, # x-axis: Treatment
    y = Tox_intra_total, # y-axis: Total intracellular toxins
    fill = Species # fill color: Species
  )) +

  # Bar plot with mean and confidence intervals
  stat_summary(
    fun.data = mean_ci, # function to compute mean and confidence interval
    geom = "bar", # geometric object: bar
    size = 4, # size of bars
    position = position_dodge(width = 0.9) # position dodge for bars
  ) +
  
  # Jittered points
  geom_jitter(
    size = 2, # size of points
    alpha = 0.6, # transparency of points
    show.legend = FALSE, # do not show legend for points
    position = position_jitterdodge(
      jitter.width = 0.7, # width of jitter
      dodge.width = 1 # width of dodge
    )
  ) +

  # Error bars for mean and confidence intervals
  stat_summary(
    fun.data = mean_ci, # function to compute mean and confidence interval
    geom = "errorbar", # geometric object: error bar
    width = 0.15, # width of error bars
    position = position_dodge(width = 0.9) # position dodge for error bars
  ) +
  
  # Configure y-axis
  scale_y_continuous(
    n.breaks = 8, # number of breaks on y-axis
    expand = c(0, 0) # remove expansion at the axis limits
  ) +
  
  # Set the y-axis limits for the viewing window
  coord_cartesian(ylim = c(0, 325)) +

  # Fill color palette
  scale_fill_brewer(
    palette = "Dark2" # use the "Dark2" palette for fill colors
  ) +
  
  # Theme and axis labels
  theme_classic() +
  theme(
    legend.position = "top", # position the legend at the top
    legend.title = element_blank() # remove legend title
  ) +
  
  # y-axis label
  ylab(
    bquote(Intracellular~toxins~(ng~mL^-1)) # y-axis label with mathematical annotation
  ) +
  
  # Horizontal line at y = 0
  geom_hline(
    yintercept = 0 # horizontal line at y = 0
  ) +
  
  # Additional theme modifications
  theme(
    axis.title.x = element_text(
      vjust = 0, 
      size = 15
    ), # vertical adjustment and font size for x-axis title
    
    axis.title.y = element_text(
      vjust = 1, 
      size = 15
    ), # vertical adjustment and font size for y-axis title
    
    axis.text = element_text(
      colour = "black", 
      size = 12
    ), # color and font size for axis text
    
    strip.background = element_rect(
      linetype = "blank"
    ), # remove background for strip
    
    strip.text = element_text(
      face = "bold.italic", 
      colour = "black", 
      size = 12
    ), # font style, color, and size for strip text
    
    legend.text = element_text(
      colour = "black", 
      size = 12, 
      face = "italic"
    ) # color, font size, and style for legend text
  )
```

Figure C20.

OBS: Note that error bars for *D. sacculus* 1 & 10 nM are
negative and out of figure bounds.

###### Extracellular toxins

```
# Plot extracellular toxin content grouped by treatment and colored by species
data_both_species %>%
  ggplot(aes(
    x = Treatment, # x-axis: Treatment
    y = Tox_extra_total, # y-axis: Total extracellular toxins
    fill = Species # fill color: Species
  )) +
  
  # Configure y-axis
  scale_y_continuous(
    limits = c(0, 140), # set y-axis limits
    n.breaks = 10, # number of breaks on y-axis
    expand = c(0, 0) # remove expansion at the axis limits
  ) +
  
  # Boxplot
  geom_boxplot(
    position = position_dodge(1) # position dodge for boxplots
  ) +
  
  # Points
  geom_point(
    position = position_dodge(1), # position dodge for points
    show.legend = FALSE # do not show legend for points
  ) +
  
  # Theme
  theme_classic() +
  theme(
    legend.position = "top" # position the legend at the top
  ) +
  
  # Fill color palette
  scale_fill_brewer(
    palette = "Dark2" # use the "Dark2" palette for fill colors
  ) +
  
  # y-axis label
  ylab(
    bquote(Extracellular~toxins~(ng~mL^-1)) # y-axis label with mathematical annotation
  )
```

Figure C21.

```
# Plot extracellular toxin content grouped by species and colored by treatment
data_both_species %>%
  ggplot(aes(
    x = Species, # x-axis: Species
    y = Tox_extra_total, # y-axis: Total extracellular toxins
    fill = Treatment # fill color: Treatment
  )) +
  
  # Configure y-axis
  scale_y_continuous(
    limits = c(0, 140), # set y-axis limits
    n.breaks = 10, # number of breaks on y-axis
    expand = c(0, 0) # remove expansion at the axis limits
  ) +
  
  # Boxplot
  geom_boxplot(
    position = position_dodge(1) # position dodge for boxplots
  ) +
  
  # Points
  geom_point(
    position = position_dodge(1), # position dodge for points
    show.legend = FALSE # do not show legend for points
  ) +
  
  # Theme
  theme_classic() +
  theme(
    legend.position = "top" # position the legend at the top
  ) +
  
  # Fill color palette
  scale_fill_manual(
    values = custom_wes_palette # use custom "Moonrise3" palette for fill colors
  ) +
  
  # y-axis label
  ylab(
    bquote(Extracellular~toxins~(ng~mL^-1)) # y-axis label with mathematical annotation
  )
```

Figure C22.

We again see the same general trend across treatments in both
species, which indicates that there is no interaction effect between the
two factors. However, toxins increase with higher concentration of
copepodamides and *D. acuminata* has considerably more
extracellular toxins than *D. sacculus* in all treatments.

Grouped bar chart:

```
# Plot extracellular toxin content grouped by treatment and colored by species
data_both_species %>%
  ggplot(aes(
    x = Treatment, # x-axis: Treatment
    y = Tox_extra_total, # y-axis: Total extracellular toxins
    fill = Species # fill color: Species
  )) +
  
  # Bar plot with mean and confidence intervals
  stat_summary(
    fun.data = mean_ci, # function to compute mean and confidence interval
    geom = "bar", # geometric object: bar
    size = 4, # size of bars
    position = position_dodge(width = 0.9) # position dodge for bars
  ) +
  
  # Jittered points
  geom_jitter(
    size = 2, # size of points
    alpha = 0.6, # transparency of points
    show.legend = FALSE, # do not show legend for points
    position = position_jitterdodge(
      jitter.width = 0.7, # width of jitter
      dodge.width = 1 # width of dodge
    )
  ) +
  
  # Error bars for mean and confidence intervals
  stat_summary(
    fun.data = mean_cl_normal, # function to compute mean and normal confidence interval
    geom = "errorbar", # geometric object: error bar
    width = 0.15, # width of error bars
    size = 0.5, # line width of error bars
    position = position_dodge(width = 0.9) # position dodge for error bars
  ) +
  
  # Configure y-axis
  scale_y_continuous(
    limits = c(0, 175), # set y-axis limits
    n.breaks = 8, # number of breaks on y-axis
    expand = c(0, 0) # remove expansion at the axis limits
  ) +
  
  # Fill color palette
  scale_fill_brewer(
    palette = "Dark2" # use the "Dark2" palette for fill colors
  ) +
  
  # Theme and axis labels
  theme_classic() +
  theme(
    legend.position = "top", # position the legend at the top
    legend.title = element_blank() # remove legend title
  ) +
  
  # y-axis label
  ylab(
    bquote(Extracellular~toxins~(ng~mL^-1)) # y-axis label with mathematical annotation
  ) +
  
  # Horizontal line at y = 0
  geom_hline(
    yintercept = 0 # horizontal line at y = 0
  ) +
  
  # Additional theme modifications
  theme(
    axis.title.x = element_text(
      vjust = 0, 
      size = 15
    ), # vertical adjustment and font size for x-axis title
    
    axis.title.y = element_text(
      vjust = 1, 
      size = 15
    ), # vertical adjustment and font size for y-axis title
    
    axis.text = element_text(
      colour = "black", 
      size = 12
    ), # color and font size for axis text
    
    strip.background = element_rect(
      linetype = "blank"
    ), # remove background for strip
    
    strip.text = element_text(
      face = "bold.italic", 
      colour = "black", 
      size = 12
    ), # font style, color, and size for strip text
    
    legend.text = element_text(
      colour = "black", 
      size = 12, 
      face = "italic"
    ) # color, font size, and style for legend text
  )
```

Figure C23.

##### Statistical analysis

Here we test whether the proportions of intracellular and
extracellular toxins change due to treatment and if it differs between
the species. To analyse proportion/percentage data, we first logit
(\(ln(p/[1-p])\)) transform the data
(1) and then fit a
Two-Way ANOVA using the transformed intracellular toxin proportion as
response variable (using the extracellular proportion would give the
same final results but inverse), and both treatment and species as
categorical independent variables (including their interaction).

```
# Fit a linear model to analyze the effect of species and treatment on the logit-transformed proportion of intracellular toxins
prop_tox_two_fact <- lm(
  logit(prop_intra) ~ Species * Treatment, 
  data = data_both_species # dataset: data_both_species
)
```

Evaluate assumption of normality:

```
# Histogram of residuals
hist(
  prop_tox_two_fact$residuals, # residuals from the linear model
  xlab = "Residuals", # x-axis label
  main = "Histogram of Residuals" # title of the histogram
)
```

```
# Shapiro-Wilks test
shapiro.test(
  prop_tox_two_fact$residuals # residuals from the linear model
)
```

```
## 
##  Shapiro-Wilk normality test
## 
## data:  prop_tox_two_fact$residuals
## W = 0.93613, p-value = 0.07156
```

Now homoscedasticity:

```
# Levene's test for homogeneity of variances
levene_test(
  logit(prop_intra) ~ Species * Treatment,
  data = data_both_species # dataset: data_both_species
)
```

```
## # A tibble: 1 × 4
##     df1   df2 statistic     p
##   <int> <int>     <dbl> <dbl>
## 1     9    20     0.600 0.783
```

The transformed proportion data is approximately normal in
distribution and has equal variances, so we will move on to perform the
actual analysis on our fitted linear model.

```
# Perform ANOVA on the linear model to analyze the effect of species and 
# treatment on the logit-transformed proportion of intracellular toxins
anova_test(
  prop_tox_two_fact, # linear model
  effect.size = "pes", # calculate partial eta squared as the effect size
  detailed = TRUE # provide detailed output
)
```

```
## ANOVA Table (type II tests)
## 
##              Effect    SSn   SSd DFn DFd       F        p p<.05   pes
## 1           Species 37.146 1.583   1  20 469.250 2.34e-15     * 0.959
## 2         Treatment 29.846 1.583   4  20  94.256 1.10e-12     * 0.950
## 3 Species:Treatment  1.387 1.583   4  20   4.381 1.00e-02     * 0.467
```

Both main effects and their interaction are significantly different
from zero. Let’s visualise this so that we can more easily determine the
likely reason for the significant interaction.

```
# Filter and rename columns for the plot
desc_mean_ci_all %>% 
  filter(Source != "Total") %>% 
  dplyr::rename(
    Mean_prop = Mean_prop, # rename for clarity
    Lower_prop = LCI_prop, # rename lower confidence interval
    Upper_prop = HCI_prop  # rename upper confidence interval
  ) %>% 
  mutate(
    Lower_prop = Lower_prop - Mean_prop, # adjust lower confidence interval
    Upper_prop = Upper_prop - Mean_prop  # adjust upper confidence interval
  ) %>% 
  group_by(Species, Treatment) %>% 
  arrange(
    rev(Source),  # arrange by source
    by.group = TRUE # arrange within groups
  ) %>% 
  mutate(
    mean2 = cumsum(Mean_prop), # cumulative sum of mean proportions
    lower = Lower_prop + mean2, # adjust lower confidence interval
    upper = Upper_prop + mean2  # adjust upper confidence interval
  ) %>% 
  
  # Create ggplot
  ggplot(aes(
    y = Mean_prop, # y-axis: mean proportion
    x = Treatment, # x-axis: treatment
    group = Source # group by source
  )) +
  
  # Stacked bars
  geom_bar(
    aes(
      y = Mean_prop, 
      fill = Source # fill color by source
    ),
    stat = "identity", # use raw values
    colour = "black", # border color of bars
    show.legend = TRUE # show legend
  ) +
  
  # Error bars
  geom_errorbar(
    aes(
      ymin = lower, # lower confidence interval
      ymax = upper  # upper confidence interval
    ),
    position = position_dodge(width = 0.3), # position dodge for error bars
    width = 0.15 # width of error bars
  ) +
  
  # Configure y-axis
  scale_y_continuous(
    limits = c(0, 1.5), # set y-axis limits
    n.breaks = 7, # number of breaks on y-axis
    labels = scales::percent, # format y-axis labels as percentages
    minor_breaks = NULL, # remove minor breaks
    expand = c(0, 0) # remove expansion at the axis limits
  ) +
  
  # Set the y-axis limits for the viewing window
  coord_cartesian(ylim = c(0, 1)) +
  
  # Facet by species
  facet_wrap(~Species, nrow = 2) +
  
  # Fill color palette
  scale_fill_manual(values = c('lightgray', 'white')) +
  
  # Theme and axis labels
  theme_classic() +
  theme(
    legend.position = "top", # position the legend at the top
    legend.title = element_blank() # remove legend title
  ) +
  
  # x-axis label
  xlab("Treatment") +
  
  # y-axis label
  ylab(bquote(Proportion~of~total~toxins~("%"))) +
  
  # Horizontal line at y = 0
  geom_hline(
    yintercept = 0 # horizontal line at y = 0
  ) +
  
  # Additional theme modifications
  theme(
    axis.title.x = element_text(
      vjust = 0, 
      size = 15
    ), # vertical adjustment and font size for x-axis title
    
    axis.title.y = element_text(
      vjust = 1, 
      size = 15
    ), # vertical adjustment and font size for y-axis title
    
    axis.text = element_text(
      colour = "black", 
      size = 12
    ), # color and font size for axis text
    
    strip.background = element_rect(
      linetype = "blank"
    ), # remove background for strip
    
    strip.text = element_text(
      face = "bold.italic", 
      colour = "black", 
      size = 12,
      vjust = 2
    ), # font style, color, and size for strip text
    
    legend.text = element_text(
      colour = "black", 
      size = 12
    ) # color and font size for legend text
  )
```

Figure C24.

Based on the stacked bar chart of proportions, we can conclude that
the significant interaction is likely due to the difference in negative
trends in intracellular toxins between the two species, specifically the
slightly parabolic change in *D. sacculus* compared to the
consistent negative change in *D. acuminata* as copepodamide
concentration increases. To determine any pairwise differences, we
perform Tukey’s Honest Significant Differences post-hoc procedure

```
# Perform Tukey's HSD post hoc test on the proportion of intracellular toxins
post_hoc_prop <- as.data.frame(tukey_hsd(prop_tox_two_fact))
```

```
# Display post hoc results in a styled table with scrolling
post_hoc_prop %>%
  dplyr::select(
    group1,        # select first group comparison
    group2,        # select second group comparison
    p.adj,         # select adjusted p-value
    p.adj.signif   # select significance of adjusted p-value
  ) %>%
  arrange(p.adj) %>% # arrange by adjusted p-value
  kable() %>%
  kable_styling("striped") %>%
  scroll_box(
    width = "100%", # set width of the scroll box
    height = "450px" # set height of the scroll box
  )
```

Table C9

| group1 | group2 | p.adj | p.adj.signif |
| --- | --- | --- | --- |
| D. sacculus | D. acuminata | 0.00e+00 | \*\*\*\* |
| D. sacculus:Copepod | D. acuminata:10 nM | 0.00e+00 | \*\*\*\* |
| D. sacculus:1 nM | D. acuminata:10 nM | 0.00e+00 | \*\*\*\* |
| D. sacculus:Control | D. acuminata:10 nM | 0.00e+00 | \*\*\*\* |
| D. sacculus:Copepod | D. acuminata:5 nM | 0.00e+00 | \*\*\*\* |
| D. sacculus:1 nM | D. acuminata:5 nM | 0.00e+00 | \*\*\*\* |
| D. sacculus:Control | D. acuminata:5 nM | 0.00e+00 | \*\*\*\* |
| Copepod | 10 nM | 0.00e+00 | \*\*\*\* |
| Control | 10 nM | 0.00e+00 | \*\*\*\* |
| D. sacculus:5 nM | D. acuminata:10 nM | 0.00e+00 | \*\*\*\* |
| 1 nM | 10 nM | 0.00e+00 | \*\*\*\* |
| D. acuminata:Control | D. acuminata:10 nM | 0.00e+00 | \*\*\*\* |
| Copepod | 5 nM | 0.00e+00 | \*\*\*\* |
| Control | 5 nM | 0.00e+00 | \*\*\*\* |
| D. sacculus:5 nM | D. acuminata:5 nM | 0.00e+00 | \*\*\*\* |
| D. sacculus:Copepod | D. acuminata:1 nM | 0.00e+00 | \*\*\*\* |
| D. acuminata:Copepod | D. acuminata:10 nM | 0.00e+00 | \*\*\*\* |
| D. sacculus:10 nM | D. acuminata:10 nM | 0.00e+00 | \*\*\*\* |
| D. acuminata:Control | D. acuminata:5 nM | 1.00e-07 | \*\*\*\* |
| D. sacculus:Copepod | D. sacculus:10 nM | 1.00e-07 | \*\*\*\* |
| D. sacculus:1 nM | D. acuminata:1 nM | 2.00e-07 | \*\*\*\* |
| D. sacculus:Copepod | D. acuminata:Copepod | 2.00e-07 | \*\*\*\* |
| 1 nM | 5 nM | 2.00e-07 | \*\*\*\* |
| D. sacculus:Control | D. acuminata:1 nM | 3.00e-07 | \*\*\*\* |
| D. acuminata:1 nM | D. acuminata:10 nM | 4.00e-07 | \*\*\*\* |
| D. acuminata:Copepod | D. acuminata:5 nM | 6.00e-07 | \*\*\*\* |
| D. acuminata:5 nM | D. sacculus:10 nM | 1.00e-06 | \*\*\*\* |
| D. acuminata:Control | D. sacculus:Copepod | 2.30e-06 | \*\*\*\* |
| D. sacculus:1 nM | D. sacculus:10 nM | 3.00e-06 | \*\*\*\* |
| D. sacculus:Control | D. sacculus:10 nM | 4.90e-06 | \*\*\*\* |
| D. acuminata:Copepod | D. sacculus:1 nM | 4.90e-06 | \*\*\*\* |
| D. sacculus:Control | D. acuminata:Copepod | 8.10e-06 | \*\*\*\* |
| D. acuminata:1 nM | D. acuminata:5 nM | 2.07e-05 | \*\*\*\* |
| D. sacculus:Copepod | D. sacculus:5 nM | 2.54e-05 | \*\*\*\* |
| D. acuminata:Control | D. sacculus:1 nM | 8.61e-05 | \*\*\*\* |
| D. sacculus:Control | D. acuminata:Control | 1.49e-04 | \*\*\* |
| D. sacculus:1 nM | D. sacculus:5 nM | 1.21e-03 | \*\* |
| D. sacculus:Control | D. sacculus:5 nM | 2.14e-03 | \*\* |
| D. acuminata:1 nM | D. sacculus:5 nM | 6.30e-03 | \*\* |
| 5 nM | 10 nM | 1.44e-02 |  |
| D. acuminata:Control | D. acuminata:1 nM | 8.36e-02 | ns |
| Copepod | 1 nM | 1.08e-01 | ns |
| D. sacculus:5 nM | D. sacculus:10 nM | 1.62e-01 | ns |
| Control | 1 nM | 2.44e-01 | ns |
| D. acuminata:Copepod | D. sacculus:5 nM | 2.52e-01 | ns |
| D. acuminata:5 nM | D. acuminata:10 nM | 5.27e-01 | ns |
| D. sacculus:Control | D. sacculus:Copepod | 5.53e-01 | ns |
| D. acuminata:Copepod | D. acuminata:1 nM | 6.98e-01 | ns |
| D. sacculus:Copepod | D. sacculus:1 nM | 7.11e-01 | ns |
| D. acuminata:Control | D. sacculus:10 nM | 7.87e-01 | ns |
| D. acuminata:1 nM | D. sacculus:10 nM | 8.38e-01 | ns |
| D. acuminata:Control | D. acuminata:Copepod | 9.03e-01 | ns |
| D. acuminata:Control | D. sacculus:5 nM | 9.57e-01 | ns |
| Control | Copepod | 9.89e-01 | ns |
| D. sacculus:Control | D. sacculus:1 nM | 1.00e+00 | ns |
| D. acuminata:Copepod | D. sacculus:10 nM | 1.00e+00 | ns |

We can see that the treatments cluster into the same two homogeneous
subgroups for both species, specifically with the treatments Control,
positive control (Copepod) and 1 nM treatments differing significantly
from the 5 and 10 nM treatments. We also find that a larger proportion
of toxins are found inside the cells for *D. sacculus* compared
to *D. acuminata* across all treatments.

**Table C10.** Statistically homogeneous groups of
intracellular toxins based on Tukey’s Honest Significant Differences
post-hoc procedure. Letters denote homogeneous groups within species,
asterisks (\*) denote a significant difference between the two species at
the given treatment level.

|  |  |  |  |  |  |
| --- | --- | --- | --- | --- | --- |
|  | Control | Copepod | 1 nM | 5 nM | 10 nM |
| D. sacculus | \* a | \* a | \* a | \* b | \* b |
| D. accuminata | \* a | \* a | \* a | \* b | \* b |

##### Final intra-/extracellular toxins plots

###### Stacked vertically

```
# Filter and rename columns for the plot
desc_mean_ci_all %>%
  filter(Source != "Total") %>%
  dplyr::rename(
    Mean_prop = Mean_prop, # rename for clarity
    Lower_prop = LCI_prop, # rename lower confidence interval
    Upper_prop = HCI_prop  # rename upper confidence interval
  ) %>%
  mutate(
    Lower_prop = Lower_prop - Mean_prop, # adjust lower confidence interval
    Upper_prop = Upper_prop - Mean_prop  # adjust upper confidence interval
  ) %>%
  group_by(Species, Treatment) %>%
  arrange(
    rev(Source),  # arrange by source
    by.group = TRUE # arrange within groups
  ) %>%
  mutate(
    mean2 = cumsum(Mean_prop), # cumulative sum of mean proportions
    lower = Lower_prop + mean2, # adjust lower confidence interval
    upper = Upper_prop + mean2  # adjust upper confidence interval
  ) %>%

  # Create ggplot
  ggplot(aes(
    y = Mean_prop, # y-axis: mean proportion
    x = Treatment, # x-axis: treatment
    group = Source # group by source
  )) +

  # Stacked bars
  geom_bar(
    aes(
      y = Mean_prop, 
      fill = Source # fill color by source
    ),
    stat = "identity", # use raw values
    colour = "black", # border color of bars
    size = 0.25, # line width of bar borders
    show.legend = TRUE # show legend
  ) +

  # Error bars
  geom_errorbar(
    aes(
      ymin = lower, # lower confidence interval
      ymax = upper  # upper confidence interval
    ),
    position = position_dodge(width = 0.6), # position dodge for error bars
    width = 0.25, # width of error bars
    linewidth = 0.25 # line width of error bars
  ) +

  # Configure y-axis
  scale_y_continuous(
    limits = c(0, 1.5), # set y-axis limits
    n.breaks = 7, # number of breaks on y-axis
    labels = scales::percent, # format y-axis labels as percentages
    minor_breaks = NULL, # remove minor breaks
    expand = c(0, 0) # remove expansion at the axis limits
  ) +

  # Set the y-axis limits for the viewing window
  coord_cartesian(ylim = c(0, 1)) +

  # Facet by species
  facet_wrap(~Species, nrow = 2) +

  # Colours and theme modifications
  scale_fill_manual(values = c('white', 'lightgray')) + 

  # x-axis label
  xlab("") +

  # y-axis label
  ylab(bquote(Proportion~of~total~toxins~("%"))) +

  # Horizontal line at y = 0
  geom_hline(yintercept = 0) +

  # Theme and axis labels
  theme_classic() +
  theme(
    plot.margin = unit(c(0.1, 0.1, -0.15, 0), "cm"), # set plot margins (top, right, bottom, left)
    
    legend.position = "top", # position the legend at the top
    
    legend.title = element_blank(), # remove legend title
    
    axis.title.y = element_text(
      vjust = 0, 
      size = 10
    ), # vertical adjustment and font size for y-axis title
    
    axis.text.y = element_text(
      colour = "black", 
      size = 8
    ), # color and font size for y-axis text
    
    axis.text.x = element_text(
      colour = "black", 
      size = 7.5
    ), # color and font size for x-axis text
    
    strip.background = element_rect(
      linetype = "blank"
    ), # remove background for strip
    
    strip.text = element_text(
      face = "italic", 
      colour = "black", 
      size = 10
    ), # font style, color, and size for strip text
    
    legend.text = element_text(
      colour = "black", 
      size = 10
    ), # color and font size for legend text
    
    legend.key.size = unit(0.5, "cm"), # size of legend keys
    
    legend.box.margin = margin(c(0, 0, -0.2, 0), unit = "cm"), # margin for legend box
    
    legend.margin = margin(c(0, 0.15, -5, 0), "cm") # margin for legend
  ) +

  # Annotations
  # For D. sacculus
  geom_text(
    data = subset(desc_mean_ci_all, 
                  Species == "D. sacculus"), # use subset of data for D. sacculus
    aes(
      x = 1, # x-position of the text
      y = 0.03, # y-position of the text
      label = "a", # text label
      family = "sans" # font family
    ),
    show.legend = FALSE, # do not show legend for text
    size = 3 # size of text
)+
  
  geom_text(
    data = subset(desc_mean_ci_all, Species == "D. sacculus"),
    aes(
      x = 2, 
      y = 0.03, 
      label = "a", 
      family = "sans"
    ),
    show.legend = FALSE, 
    size = 3
  ) +
  geom_text(
    data = subset(desc_mean_ci_all, Species == "D. sacculus"),
    aes(
      x = 3, 
      y = 0.03, 
      label = "a", 
      family = "sans"
    ),
    show.legend = FALSE, 
    size = 3
  ) +
  geom_text(
    data = subset(desc_mean_ci_all, Species == "D. sacculus"),
    aes(
      x = 4, 
      y = 0.03, 
      label = "b", 
      family = "sans"
    ),
    show.legend = FALSE, 
    size = 3
  ) +
  geom_text(
    data = subset(desc_mean_ci_all, Species == "D. sacculus"),
    aes(
      x = 5, 
      y = 0.03, 
      label = "b", 
      family = "sans"
    ),
    show.legend = FALSE, 
    size = 3
  ) +
  
  # For D. acuminata
  geom_text(
    data = subset(desc_mean_ci_all, Species == "D. acuminata"),
    aes(
      x = 1, 
      y = 0.03, 
      label = "a", 
      family = "sans"
    ),
    show.legend = FALSE, 
    size = 3
  ) +
  geom_text(
    data = subset(desc_mean_ci_all, Species == "D. acuminata"),
    aes(
      x = 2, 
      y = 0.03, 
      label = "a", 
      family = "sans"
    ),
    show.legend = FALSE, 
    size = 3
  ) +
  geom_text(
    data = subset(desc_mean_ci_all, Species == "D. acuminata"),
    aes(
      x = 3, 
      y = 0.03, 
      label = "a", 
      family = "sans"
    ),
    show.legend = FALSE, 
    size = 3
  ) +
  geom_text(
    data = subset(desc_mean_ci_all, Species == "D. acuminata"),
    aes(
      x = 4, 
      y = 0.03, 
      label = "b", 
      family = "sans"
    ),
    show.legend = FALSE, 
    size = 3
  ) +
  geom_text(
    data = subset(desc_mean_ci_all, Species == "D. acuminata"),
    aes(
      x = 5, 
      y = 0.03, 
      label = "b", 
      family = "sans"
    ),
    show.legend = FALSE, 
    size = 3
  )
```

Figure C25. Proportion of intracellular (grey) and extracellular (white)
toxins for *Dinophysis sacculus* and *Dinophysis
acuminata* after 68.5 hours of exposure to three concentrations of
chemical cues from copepods (copepodamides), a positive control (one
live *Acartia* sp.) or a control treatment. Bars are the means of
three replicates (n = 3) and error bars show the 95% confidence
intervals of the means. Letters denote statistically homogeneous subsets
within species based on Tukey’s Honest Significant Differences post-hoc
procedure with ⍺ set to 0.05. Note that both upper and lower confidence
limits are shown for intracellular bars, but only the lower limit is
shown for the extracellular bars, this is to avoid a y-axis > 100%
while still enabling simple comparison of any overlapping intervals.

```
# ggsave("Inta_Extra_plot_Dino.tiff", width = 80, height = 150, units = "mm", dpi=700)
```

###### Ordered horizontally

```
# Filter and rename columns for the plot
desc_mean_ci_all %>%
  filter(Source != "Total") %>%
  dplyr::rename(
    Mean_prop = Mean_prop, # rename for clarity
    Lower_prop = LCI_prop, # rename lower confidence interval
    Upper_prop = HCI_prop  # rename upper confidence interval
  ) %>%
  mutate(
    Lower_prop = Lower_prop - Mean_prop, # adjust lower confidence interval
    Upper_prop = Upper_prop - Mean_prop  # adjust upper confidence interval
  ) %>%
  group_by(Species, Treatment) %>%
  arrange(
    rev(Source),  # arrange by source
    by.group = TRUE # arrange within groups
  ) %>%
  mutate(
    mean2 = cumsum(Mean_prop), # cumulative sum of mean proportions
    lower = Lower_prop + mean2, # adjust lower confidence interval
    upper = Upper_prop + mean2  # adjust upper confidence interval
  ) %>%

  # Create ggplot
  ggplot(aes(
    y = Mean_prop, # y-axis: mean proportion
    x = Treatment, # x-axis: treatment
    group = Source # group by source
  )) +

  # Stacked bars
  geom_bar(
    aes(
      y = Mean_prop, 
      fill = Source # fill color by source
    ),
    stat = "identity", # use raw values
    colour = "black", # border color of bars
    size = 0.25, # line width of bar borders
    show.legend = TRUE # show legend
  ) +

  # Error bars
  geom_errorbar(
    aes(
      ymin = lower, # lower confidence interval
      ymax = upper  # upper confidence interval
    ),
    position = position_dodge(width = 0.6), # position dodge for error bars
    width = 0.25, # width of error bars
    linewidth = 0.25 # line width of error bars
  ) +

  # Configure y-axis
  scale_y_continuous(
    limits = c(0, 1.5), # set y-axis limits
    n.breaks = 7, # number of breaks on y-axis
    labels = scales::percent, # format y-axis labels as percentages
    minor_breaks = NULL, # remove minor breaks
    expand = c(0, 0) # remove expansion at the axis limits
  ) +

  # Set the y-axis limits for the viewing window
  coord_cartesian(ylim = c(0, 1)) +

  # Facet by species
  facet_wrap(~Species, ncol = 2) +

  # Colours and theme modifications
  scale_fill_manual(values = c('white', 'lightgray')) + 

  # x-axis label
  xlab("") +

  # y-axis label
  ylab(bquote(Proportion~of~total~toxins~("%"))) +

  # Horizontal line at y = 0
  geom_hline(yintercept = 0) +

  # Theme and axis labels
  theme_classic() +
  theme(
    plot.margin = unit(c(0.1, 0.1, -0.15, 0), "cm"), # set plot margins (top, right, bottom, left)
    
    legend.position = "top", # position the legend at the top
    
    legend.title = element_blank(), # remove legend title
    
    axis.title.y = element_text(
      vjust = -0.5, 
      size = 10
    ), # vertical adjustment and font size for y-axis title
    
    axis.text.y = element_text(
      colour = "black", 
      size = 8
    ), # color and font size for y-axis text
    
    axis.text.x = element_text(
      colour = "black", 
      size = 7.5
    ), # color and font size for x-axis text
    
    strip.background = element_rect(
      linetype = "blank"
    ), # remove background for strip
    
    strip.text = element_text(
      face = "italic", 
      colour = "black", 
      size = 10
    ), # font style, color, and size for strip text
    
    legend.text = element_text(
      colour = "black", 
      size = 10
    ), # color and font size for legend text
    
    legend.key.size = unit(0.5, "cm"), # size of legend keys
    
    legend.box.margin = margin(c(0, 0, -0.2, 0), unit = "cm"), # margin for legend box
    
    legend.margin = margin(c(0, 0.15, -5, 0), "cm") # margin for legend
  ) +

  # Annotations for D. sacculus
  geom_text(
    data = subset(desc_mean_ci_all, Species == "D. sacculus"), # use subset of data for D. sacculus
    aes(
      x = 1, # x-position of the text
      y = 0.03, # y-position of the text
      label = "a", # text label
      family = "sans" # font family
    ),
    show.legend = FALSE, # do not show legend for text
    size = 3 # size of text
  ) +
  geom_text(
    data = subset(desc_mean_ci_all, Species == "D. sacculus"), # use subset of data for D. sacculus
    aes(
      x = 2, # x-position of the text
      y = 0.03, # y-position of the text
      label = "a", # text label
      family = "sans" # font family
    ),
    show.legend = FALSE, # do not show legend for text
    size = 3 # size of text
  ) +
  geom_text(
    data = subset(desc_mean_ci_all, Species == "D. sacculus"), # use subset of data for D. sacculus
    aes(
      x = 3, # x-position of the text
      y = 0.03, # y-position of the text
      label = "a", # text label
      family = "sans" # font family
    ),
    show.legend = FALSE, # do not show legend for text
    size = 3 # size of text
  ) +
  geom_text(
    data = subset(desc_mean_ci_all, Species == "D. sacculus"), # use subset of data for D. sacculus
    aes(
      x = 4, # x-position of the text
      y = 0.03, # y-position of the text
      label = "b", # text label
      family = "sans" # font family
    ),
    show.legend = FALSE, # do not show legend for text
    size = 3 # size of text
  ) +
  geom_text(
    data = subset(desc_mean_ci_all, Species == "D. sacculus"), # use subset of data for D. sacculus
    aes(
      x = 5, # x-position of the text
      y = 0.03, # y-position of the text
      label = "b", # text label
      family = "sans" # font family
    ),
    show.legend = FALSE, # do not show legend for text
    size = 3 # size of text
  ) +

  # Annotations for D. acuminata
  geom_text(
    data = subset(desc_mean_ci_all, Species == "D. acuminata"), # use subset of data for D. acuminata
    aes(
      x = 1, # x-position of the text
      y = 0.03, # y-position of the text
      label = "a", # text label
      family = "sans" # font family
    ),
    show.legend = FALSE, # do not show legend for text
    size = 3 # size of text
  ) +
  geom_text(
    data = subset(desc_mean_ci_all, Species == "D. acuminata"), # use subset of data for D. acuminata
    aes(
      x = 2, # x-position of the text
      y = 0.03, # y-position of the text
      label = "a", # text label
      family = "sans" # font family
    ),
    show.legend = FALSE, # do not show legend for text
    size = 3 # size of text
  ) +
  geom_text(
    data = subset(desc_mean_ci_all, Species == "D. acuminata"), # use subset of data for D. acuminata
    aes(
      x = 3, # x-position of the text
      y = 0.03, # y-position of the text
      label = "a", # text label
      family = "sans" # font family
    ),
    show.legend = FALSE, # do not show legend for text
    size = 3 # size of text
  ) +
  geom_text(
    data = subset(desc_mean_ci_all, Species == "D. acuminata"), # use subset of data for D. acuminata
    aes(
      x = 4, # x-position of the text
      y = 0.03, # y-position of the text
      label = "b", # text label
      family = "sans" # font family
    ),
    show.legend = FALSE, # do not show legend for text
    size = 3 # size of text
  ) +
  geom_text(
    data = subset(desc_mean_ci_all, Species == "D. acuminata"), # use subset of data for D. acuminata
    aes(
      x = 5, # x-position of the text
      y = 0.03, # y-position of the text
      label = "b", # text label
      family = "sans" # font family
    ),
    show.legend = FALSE, # do not show legend for text
    size = 3 # size of text
  )
```

Figure C26. Proportion of intracellular (grey) and extracellular (white)
toxins for *Dinophysis sacculus* and *Dinophysis
acuminata* after 68.5 hours of exposure to three concentrations of
chemical cues from copepods (copepodamides), a positive control (one
live *Acartia* sp.) or a control treatment. Bars are the means of
three replicates (n = 3) and error bars show the 95% confidence
intervals of the means. Letters denote statistically homogeneous subsets
within species based on Tukey’s Honest Significant Differences post-hoc
procedure with ⍺ set to 0.05. Note that both upper and lower confidence
limits are shown for intracellular bars, but only the lower limit is
shown for the extracellular bars, this is to avoid a y-axis > 100%
while still enabling simple comparison of any overlapping intervals.

```
#ggsave(path = "figures_manuscript","Inta_Extra_plot_Dino_hori.tiff", width = 180, height = 80, units = "mm", dpi=700)
```

#### Analysis of toxin composition

##### Exploratory data analysis and visualisation

###### Summary statistics tables

###### Okadaic acid

```
# Calculate descriptive statistics for OA toxin content and proportion
mean_ci_OA <- data_both_species %>%
  dplyr::select(
    Species, # select species
    Treatment, # select treatment
    OA_total, # select total OA toxin content
    prop_OA # select proportion of OA toxin content
  ) %>%
  group_by(Species, Treatment) %>%
  summarise(
    N = length(OA_total), # sample size
    Mean = mean(OA_total), # mean total OA toxin content
    St.Dev = sd(OA_total), # standard deviation of total OA toxin content
    SEM = St.Dev / sqrt(N), # standard error of the mean
    CI = qt(0.975, df = N - 1) * SEM, # 95% confidence interval
    LCI = Mean - CI, # lower confidence interval
    HCI = Mean + CI, # upper confidence interval
    Mean_prop = mean(prop_OA), # mean proportion of OA toxin content
    St.Dev_prop = sd(prop_OA), # standard deviation of proportion of OA toxin content
    SEM_prop = St.Dev_prop / sqrt(N), # standard error of the mean for proportion
    CI_prop = qt(0.975, df = N - 1) * SEM_prop, # 95% confidence interval for proportion
    LCI_prop = Mean_prop - CI_prop, # lower confidence interval for proportion
    HCI_prop = Mean_prop + CI_prop # upper confidence interval for proportion
  ) %>%
  data.frame() %>%   
  mutate(Toxin = "OA") # add toxin type
```

###### PTX2

```
# Calculate descriptive statistics for PTX2 toxin content and proportion in the same way
mean_ci_PTX2 <-
 data_both_species %>% 
  dplyr::select(Species, Treatment, PTX2_total, prop_PTX2) %>% 
  group_by(Species, Treatment) %>% 
  summarise(N = length(PTX2_total),
            Mean = mean(PTX2_total),
            St.Dev = sd(PTX2_total),
            SEM = St.Dev / sqrt(N),
            CI = qt(0.975, df = N-1) * SEM,
            LCI = Mean - CI,
            HCI = Mean + CI,
            Mean_prop = mean(prop_PTX2),
            St.Dev_prop = sd(prop_PTX2),
            SEM_prop = St.Dev_prop / sqrt(N),
            CI_prop = qt(0.975, df = N-1) * SEM_prop,
            LCI_prop = Mean_prop - CI_prop,
            HCI_prop = Mean_prop + CI_prop) %>% 
  data.frame() %>%   
  mutate(Toxin = "PTX2")
```

##### C9

```
# Calculate descriptive statistics for C9 toxin content and proportion in the same way
mean_ci_C9  <-
 data_both_species %>% 
  dplyr::select(Species, Treatment, C9_total, prop_C9) %>% 
  group_by(Species, Treatment) %>% 
  summarise(N = length(C9_total),
            Mean = mean(C9_total),
            St.Dev = sd(C9_total),
            SEM = St.Dev / sqrt(N),
            CI = qt(0.975, df = N-1) * SEM,
            LCI = Mean - CI,
            HCI = Mean + CI,
            Mean_prop = mean(prop_C9),
            St.Dev_prop = sd(prop_C9),
            SEM_prop = St.Dev_prop / sqrt(N),
            CI_prop = qt(0.975, df = N-1) * SEM_prop,
            LCI_prop = Mean_prop - CI_prop,
            HCI_prop = Mean_prop + CI_prop) %>% 
  data.frame() %>%   
  mutate(Toxin = "C9")
```

###### Combining the tables

```
desc_mean_ci_toxin_all <-
  bind_rows(mean_ci_OA, 
            mean_ci_PTX2, 
            mean_ci_C9) %>% 
  dplyr::select(Species,Treatment,Toxin, 
                N, Mean, LCI, HCI, 
                Mean_prop, LCI_prop, HCI_prop) %>% 
  mutate(across(
    where(is.numeric), 
    round, 
    digits = 3 # round numeric variables to 3 digits
  )) %>% 
  data.frame()

desc_mean_ci_toxin_all$Toxin <- factor(desc_mean_ci_toxin_all$Toxin, 
                                       levels = c("C9", "PTX2", "OA"))


# Combine the descriptive statistics for OA, PTX2, and C9 toxins
desc_mean_ci_toxin_all <- bind_rows(
  mean_ci_OA, 
  mean_ci_PTX2, 
  mean_ci_C9
) %>%
  dplyr::select(
    Species, # select species
    Treatment, # select treatment
    Toxin, # select toxin type
    N, # sample size
    Mean, # mean toxin content
    LCI, # lower confidence interval
    HCI, # upper confidence interval
    Mean_prop, # mean proportion of toxin content
    LCI_prop, # lower confidence interval for proportion
    HCI_prop # upper confidence interval for proportion
  ) %>%
  mutate(across(
    where(is.numeric), 
    round, 
    digits = 3 # round numeric variables to 3 digits
  )) %>%
  data.frame()

# Set the order of levels in the Toxin factor
desc_mean_ci_toxin_all$Toxin <- factor(
  desc_mean_ci_toxin_all$Toxin, 
  levels = c("C9", "PTX2", "OA")
)
```

```
# Display the final rounded data frame in a styled table with scrolling
desc_mean_ci_toxin_all %>%
  kable() %>%
  kable_styling("striped") %>%
  scroll_box(
    width = "100%", # set width of the scroll box
    height = "450px" # set height of the scroll box
  )
```

Table C11.

| Species | Treatment | Toxin | N | Mean | LCI | HCI | Mean\_prop | LCI\_prop | HCI\_prop |
| --- | --- | --- | --- | --- | --- | --- | --- | --- | --- |
| D. sacculus | Control | OA | 3 | 7.209 | 3.563 | 10.855 | 0.049 | 0.044 | 0.054 |
| D. sacculus | Copepod | OA | 3 | 7.922 | 5.462 | 10.382 | 0.049 | 0.044 | 0.055 |
| D. sacculus | 1 nM | OA | 3 | 8.810 | -1.319 | 18.938 | 0.048 | 0.040 | 0.055 |
| D. sacculus | 5 nM | OA | 3 | 6.589 | 4.709 | 8.469 | 0.045 | 0.044 | 0.046 |
| D. sacculus | 10 nM | OA | 3 | 8.073 | 0.491 | 15.656 | 0.044 | 0.037 | 0.052 |
| D. acuminata | Control | OA | 3 | 122.088 | 89.733 | 154.444 | 1.000 | 1.000 | 1.000 |
| D. acuminata | Copepod | OA | 3 | 147.195 | 67.981 | 226.408 | 1.000 | 1.000 | 1.000 |
| D. acuminata | 1 nM | OA | 3 | 134.272 | 95.174 | 173.370 | 1.000 | 1.000 | 1.000 |
| D. acuminata | 5 nM | OA | 3 | 119.447 | 92.507 | 146.387 | 1.000 | 1.000 | 1.000 |
| D. acuminata | 10 nM | OA | 3 | 125.400 | 96.929 | 153.870 | 1.000 | 1.000 | 1.000 |
| D. sacculus | Control | PTX2 | 3 | 137.721 | 74.744 | 200.697 | 0.939 | 0.936 | 0.941 |
| D. sacculus | Copepod | PTX2 | 3 | 150.122 | 113.756 | 186.488 | 0.938 | 0.933 | 0.944 |
| D. sacculus | 1 nM | PTX2 | 3 | 171.216 | 3.732 | 338.699 | 0.940 | 0.939 | 0.942 |
| D. sacculus | 5 nM | PTX2 | 3 | 137.968 | 94.605 | 181.332 | 0.939 | 0.930 | 0.948 |
| D. sacculus | 10 nM | PTX2 | 3 | 175.067 | -20.076 | 370.209 | 0.943 | 0.927 | 0.958 |
| D. acuminata | Control | PTX2 | 3 | 0.000 | 0.000 | 0.000 | 0.000 | 0.000 | 0.000 |
| D. acuminata | Copepod | PTX2 | 3 | 0.000 | 0.000 | 0.000 | 0.000 | 0.000 | 0.000 |
| D. acuminata | 1 nM | PTX2 | 3 | 0.000 | 0.000 | 0.000 | 0.000 | 0.000 | 0.000 |
| D. acuminata | 5 nM | PTX2 | 3 | 0.000 | 0.000 | 0.000 | 0.000 | 0.000 | 0.000 |
| D. acuminata | 10 nM | PTX2 | 3 | 0.000 | 0.000 | 0.000 | 0.000 | 0.000 | 0.000 |
| D. sacculus | Control | C9 | 3 | 1.793 | 1.444 | 2.143 | 0.012 | 0.008 | 0.017 |
| D. sacculus | Copepod | C9 | 3 | 1.921 | 1.755 | 2.087 | 0.012 | 0.009 | 0.015 |
| D. sacculus | 1 nM | C9 | 3 | 2.061 | 1.238 | 2.885 | 0.012 | 0.006 | 0.018 |
| D. sacculus | 5 nM | C9 | 3 | 2.299 | 1.784 | 2.813 | 0.016 | 0.008 | 0.024 |
| D. sacculus | 10 nM | C9 | 3 | 2.231 | 1.553 | 2.910 | 0.013 | 0.003 | 0.024 |
| D. acuminata | Control | C9 | 3 | 0.000 | 0.000 | 0.000 | 0.000 | 0.000 | 0.000 |
| D. acuminata | Copepod | C9 | 3 | 0.000 | 0.000 | 0.000 | 0.000 | 0.000 | 0.000 |
| D. acuminata | 1 nM | C9 | 3 | 0.000 | 0.000 | 0.000 | 0.000 | 0.000 | 0.000 |
| D. acuminata | 5 nM | C9 | 3 | 0.000 | 0.000 | 0.000 | 0.000 | 0.000 | 0.000 |
| D. acuminata | 10 nM | C9 | 3 | 0.000 | 0.000 | 0.000 | 0.000 | 0.000 | 0.000 |

###### Visual data exploration

###### Okadaic acid

```
# Plot Okadaic acid (OA) content grouped by species and colored by treatment
data_both_species %>%
  ggplot(aes(
    x = Species, # x-axis: Species
    y = OA_total, # y-axis: Total OA content
    fill = Treatment # fill color: Treatment
  )) + 
  
  # Configure y-axis
  scale_y_continuous(
    n.breaks = 14 # number of breaks on y-axis
  ) +
  
  # Boxplot
  geom_boxplot(
    position = position_dodge(1) # position dodge for boxplots
  ) +
  
  # Points
  geom_point(
    position = position_dodge(1), # position dodge for points
    show.legend = FALSE # do not show legend for points
  ) +
  
  # Theme
  theme_classic() +
  theme(
    legend.position = "top" # position the legend at the top
  ) +
  
  # Fill color palette
  scale_fill_manual(
    values = custom_wes_palette # use custom "Moonrise3" palette for fill colors
  ) +
  
  # y-axis label
  ylab(
    bquote(Okadaic~acid~(ng~mL^-1)) # y-axis label with mathematical annotation
  )
```

Figure C27

Because of the large difference in absolute values between the
species, we place them on a log-scale to make visual interpretation
easier.

```
# Plot log-transformed Okadaic acid (OA) content grouped by species and colored by treatment
data_both_species %>%
  ggplot(aes(
    x = Species, # x-axis: Species
    y = log(OA_total), # y-axis: Log-transformed total OA content
    fill = Treatment # fill color: Treatment
  )) + 
  
  # Configure y-axis
  scale_y_continuous(
    n.breaks = 10 # number of breaks on y-axis
  ) +
  
  # Boxplot
  geom_boxplot(
    position = position_dodge(1) # position dodge for boxplots
  ) +
  
  # Points
  geom_point(
    position = position_dodge(1), # position dodge for points
    show.legend = FALSE # do not show legend for points
  ) +
  
  # Theme
  theme_classic() +
  theme(
    legend.position = "top" # position the legend at the top
  ) +
  
  # Fill color palette
  scale_fill_manual(
    values = custom_wes_palette # use custom "Moonrise3" palette for fill colors
  ) +
  
  # y-axis label
  ylab(
    bquote(Ln~Okadaic~acid~(ng~mL^-1)) # y-axis label with mathematical annotation
  )
```

Figure C28

Same approximate trend across treatments in both species, which
indicates that there is no interaction effect between the two factors.
Also, *D. acuminata* produces much more okadaic acid than *D.
sacculus*.

Grouped bar chart to visualise this with means and their 95%
confidence limits:

```
# Plot Okadaic acid (OA) content grouped by treatment and colored by species
data_both_species %>%
  ggplot(aes(
    x = Treatment, # x-axis: Treatment
    y = OA_total, # y-axis: Total OA content
    fill = Species # fill color: Species
  )) +
  
  # Bar plot with mean and confidence intervals
  stat_summary(
    fun.data = mean_ci, # function to compute mean and confidence interval
    geom = "bar", # geometric object: bar
    size = 4, # size of bars
    position = position_dodge(width = 0.9) # position dodge for bars
  ) +
  
  # Jittered points
  geom_jitter(
    size = 2, # size of points
    alpha = 0.6, # transparency of points
    show.legend = FALSE, # do not show legend for points
    position = position_jitterdodge(
      jitter.width = 0.5, # width of jitter
      dodge.width = 0.9 # width of dodge
    )
  ) +
  
  # Error bars for mean and confidence intervals
  stat_summary(
    fun.data = mean_ci, # function to compute mean and confidence interval
    geom = "errorbar", # geometric object: error bar
    width = 0.15, # width of error bars
    linewidth = 0.5, # line width of error bars
    position = position_dodge(width = 0.9) # position dodge for error bars
  ) +
  
  # Configure y-axis
  scale_y_continuous(
    n.breaks = 9, # number of breaks on y-axis
    expand = c(0, 0) # remove expansion at the axis limits
  ) +
  
  # Set the y-axis limits for the viewing window
  coord_cartesian(ylim = c(0, 240)) +
  
  # Fill color palette
  scale_fill_brewer(palette = "Dark2") +
  
  # Theme and axis labels
  theme_classic() +
  theme(
    legend.position = "top", # position the legend at the top
    legend.title = element_blank() # remove legend title
  ) +
  
  # y-axis label
  ylab(
    bquote(Okadaic~acid~(ng~mL^-1)) # y-axis label with mathematical annotation
  ) +
  
  # Additional theme modifications
  theme(
    axis.title.x = element_text(
      vjust = 0, 
      size = 15
    ), # vertical adjustment and font size for x-axis title
    
    axis.title.y = element_text(
      vjust = 1, 
      size = 15
    ), # vertical adjustment and font size for y-axis title
    
    axis.text = element_text(
      colour = "black", 
      size = 12
    ), # color and font size for axis text
    
    strip.background = element_rect(
      linetype = "blank"
    ), # remove background for strip
    
    strip.text = element_text(
      face = "bold.italic", 
      colour = "black", 
      size = 12
    ), # font style, color, and size for strip text
    
    legend.text = element_text(
      colour = "black", 
      size = 12, 
      face = "italic"
    ) # color, font size, and style for legend text
  )
```

Figure C29

No apparent difference in okadaic acid content between treatment
groups.

###### PTX2

```
# Plot PTX2 content for D. sacculus grouped by treatment
data_both_species %>%
  filter(Species == "D. sacculus") %>% # filter for D. sacculus
  ggplot(aes(
    x = Treatment, # x-axis: Treatment
    y = PTX2_total # y-axis: Total PTX2 content
  )) +
  
  # Bar plot with mean and confidence intervals
  stat_summary(
    fun.data = mean_ci, # function to compute mean and confidence interval
    geom = "bar", # geometric object: bar
    size = 4, # size of bars
    fill = c(custom_wes_palette), # custom fill colors
    show.legend = TRUE # show legend
  ) +
  
  # Jittered points
  geom_jitter(
    size = 2, # size of points
    alpha = 0.6, # transparency of points
    show.legend = FALSE, # do not show legend for points
    width = 0.3 # width of jitter
  ) +
  
  # Error bars for mean and confidence intervals
  stat_summary(
    fun.data = mean_ci, # function to compute mean and confidence interval
    geom = "errorbar", # geometric object: error bar
    width = 0.15, # width of error bars
    size = 0.2 # line width of error bars
  ) +
  
  # Configure y-axis
  scale_y_continuous(
    limits = c(-50, 450), # set y-axis limits
    n.breaks = 10, # number of breaks on y-axis
    expand = c(0, 0) # remove expansion at the axis limits
  ) +
  
  # Set the y-axis limits for the viewing window
  coord_cartesian(ylim = c(0, 400)) +
  
  # Title
  ggtitle("D. sacculus") +
  
  # Theme and axis labels
  theme_classic() +
  theme(
    legend.position = "top", # position the legend at the top
    legend.title = element_blank() # remove legend title
  ) +
  
  # y-axis label
  ylab(
    bquote(PTX2~(ng~mL^-1)) # y-axis label with mathematical annotation
  ) +
  
  # Horizontal line at y = 0
  geom_hline(
    yintercept = 0 # horizontal line at y = 0
  ) +
  
  # Additional theme modifications
  theme(
    axis.title.x = element_text(
      vjust = 0, 
      size = 15
    ), # vertical adjustment and font size for x-axis title
    
    axis.title.y = element_text(
      vjust = 1, 
      size = 15
    ), # vertical adjustment and font size for y-axis title
    
    axis.text = element_text(
      colour = "black", 
      size = 12
    ), # color and font size for axis text
    
    strip.background = element_rect(
      linetype = "blank"
    ), # remove background for strip
    
    strip.text = element_text(
      face = "bold.italic", 
      colour = "black", 
      size = 12
    ), # font style, color, and size for strip text
    
    legend.text = element_text(
      colour = "black", 
      size = 12, 
      face = "italic"
    ), # color, font size, and style for legend text
    
    plot.title = element_text(
      hjust = 0.5, 
      vjust = -2, 
      face = "italic"
    ) # horizontal and vertical adjustment, and font style for plot title
  )
```

Figure C30

No differences in PTX2 content \(mL^1\) for *D. sacculus*

##### C9

```
# Plot C9 content grouped by species and colored by treatment
data_both_species %>% 
  ggplot(aes(x=Species, 
             y= C9_total,
             fill=Treatment)) + 
  
  scale_y_continuous(n.breaks = 10)+

  geom_boxplot(position=position_dodge(1))+
  
  geom_point(position=position_dodge(1),
             show.legend = FALSE)+
  
  theme_classic()+
  
  theme(legend.position = "top")+
  
  scale_fill_manual(values = custom_wes_palette) +
  
  ylab(bquote(C9~(ng~mL^-1)))
```

Figure C31

*D. acuminata* does not produce any C9 either. C9 also seems
to increase slightly in *D. sacculus* when exposed to
grazing/chemical cues from copepods.

Bar chart to visualise this with means and 95% confidence
intervals.

```
# Plot C9 toxin content for D. sacculus grouped by treatment
data_both_species %>%
  filter(Species == "D. sacculus") %>% # filter for D. sacculus
  ggplot(aes(
    x = Treatment, # x-axis: Treatment
    y = C9_total # y-axis: Total C9 content
  )) +
  
  # Bar plot with mean and confidence intervals
  stat_summary(
    fun.data = mean_ci, # function to compute mean and confidence interval
    geom = "bar", # geometric object: bar
    size = 4, # size of bars
    fill = custom_wes_palette, # custom fill colors
    show.legend = TRUE # show legend
  ) +
  
  # Jittered points
  geom_jitter(
    size = 2, # size of points
    alpha = 0.6, # transparency of points
    show.legend = FALSE, # do not show legend for points
    width = 0.2 # width of jitter
  ) +
  
  # Error bars for mean and confidence intervals
  stat_summary(
    fun.data = mean_ci, # function to compute mean and confidence interval
    geom = "errorbar", # geometric object: error bar
    width = 0.15, # width of error bars
    size = 0.5 # line width of error bars
  ) +
  
  # Configure y-axis
  scale_y_continuous(
    limits = c(0, 3), # set y-axis limits
    n.breaks = 9, # number of breaks on y-axis
    expand = c(0, 0) # remove expansion at the axis limits
  ) +
  
  # Title
  ggtitle("D. sacculus") +
  
  # Theme and axis labels
  theme_classic() +
  theme(
    legend.position = "top", # position the legend at the top
    legend.title = element_blank() # remove legend title
  ) +
  
  # y-axis label
  ylab(
    bquote(C9~(ng~mL^-1)) # y-axis label with mathematical annotation
  ) +
  
  # Horizontal line at y = 0
  geom_hline(
    yintercept = 0 # horizontal line at y = 0
  ) +
  
  # Additional theme modifications
  theme(
    axis.title.x = element_text(
      vjust = 0, 
      size = 15
    ), # vertical adjustment and font size for x-axis title
    
    axis.title.y = element_text(
      vjust = 1, 
      size = 15
    ), # vertical adjustment and font size for y-axis title
    
    axis.text = element_text(
      colour = "black", 
      size = 12
    ), # color and font size for axis text
    
    strip.background = element_rect(
      linetype = "blank"
    ), # remove background for strip
    
    strip.text = element_text(
      face = "bold.italic", 
      colour = "black", 
      size = 12
    ), # font style, color, and size for strip text
    
    legend.text = element_text(
      colour = "black", 
      size = 12, 
      face = "italic"
    ), # color, font size, and style for legend text
    
    plot.title = element_text(
      hjust = 0.5, 
      vjust = -2, 
      face = "italic"
    ) # horizontal and vertical adjustment, and font style for plot title
  )
```

Figure C32

Let’s combine these into one plot.

```
# Filter and rename columns for the plot
x <-
desc_mean_ci_toxin_all %>%
  filter(Mean > 0) %>%
  dplyr::rename(
    Mean = Mean, # rename for clarity
    Lower = LCI, # rename lower confidence interval
    Upper = HCI  # rename upper confidence interval
  ) %>%
  mutate(
    Lower = Lower - Mean, # adjust lower confidence interval
    Upper = Upper - Mean  # adjust upper confidence interval
  ) %>%
  group_by(Species, Treatment) %>%
  arrange(
    rev(Toxin),  # arrange by toxin type
    by.group = TRUE # arrange within groups
  ) %>%
  mutate(
    mean2 = cumsum(Mean), # cumulative sum of means
    lower = Lower + mean2, # adjust lower confidence interval
    upper = Upper + mean2  # adjust upper confidence interval
  )
```

```
x %>% 
  # Create ggplot
  ggplot(aes(
    y = Mean, # y-axis: mean toxin content
    x = Treatment, # x-axis: treatment
    group = Toxin # group by toxin type
  )) +

  # Stacked bars
  geom_bar(
    aes(
      y = Mean, 
      fill = Toxin # fill color by toxin type
    ),
    stat = "identity", # use raw values
    colour = "black", # border color of bars
    size = 0.25, # line width of bar borders
    show.legend = TRUE # show legend
  ) +

  # Configure y-axis
  scale_y_continuous(
    n.breaks = 10, # number of breaks on y-axis
    minor_breaks = NULL, # remove minor breaks
    expand = c(0, 0) # remove expansion at the axis limits
  ) +

  # Set the y-axis limits for the viewing window
  coord_cartesian(ylim = c(0, 200)) +

  # Facet by species
  facet_wrap(~Species, nrow = 1) +

  # Error bars for D. sacculus
  geom_errorbar(
    data = filter(x, Species == "D. sacculus"), # filter data for D. sacculus
    aes(
      y = mean2, # adjusted mean for positioning error bars
      ymin = lower, # lower confidence interval
      ymax = mean2 # adjusted upper confidence interval
    ),
    position = position_dodge(width = 0.8), # position dodge for error bars
    width = 0.25, # width of error bars
    linewidth = 0.25 # line width of error bars
  ) +

  # Error bars for D. acuminata
  geom_errorbar(
    data = filter(x, Species == "D. acuminata"), # filter data for D. acuminata
    aes(
      y = mean2, # adjusted mean for positioning error bars
      ymin = lower, # lower confidence interval
      ymax = mean2 # adjusted upper confidence interval
    ),
    position = position_dodge(width = 0.8), # position dodge for error bars
    width = 0.1, # width of error bars
    linewidth = 0.25 # line width of error bars
  ) +

  # Fill color palette
  scale_fill_manual(
    values = c("grey70", "grey85", "grey100") # set fill colors
  ) +

  # Horizontal line at y = 0
  geom_hline(yintercept = 0) +

  # Axis labels
  xlab("Treatment") +
  ylab(
    bquote(Toxin~content~(ng~mL^-1)) # y-axis label with mathematical annotation
  ) +
  xlab("") +

  # Theme and axis labels
  theme_classic() +
  theme(
    legend.position = c(0.08, 0.9), # position the legend
    
    # legend settings
    legend.direction = "vertical",
    legend.background = element_rect( # transparent legend background
      fill = 'transparent', 
      colour = 'transparent'
    ), 
    legend.box.background = element_rect( # transparent legend panel background
      fill = 'transparent', 
      colour = 'transparent'
    ), 
    legend.title = element_blank(), # remove legend title
    
    legend.text = element_text( # legend text settings
      colour = "black",
      size = 8
    ),
    legend.box.margin = margin(
      c(0, -0.2, 0, -0.5), 
      unit = "cm"
    ),
    legend.key.size = unit(0.4, "cm"), # size of legend keys

    # y-axis title settings
    axis.title.y = element_text(
      vjust = 0, 
      size = 10
    ), 

    # axis text settings
    axis.text.y = element_text(
      colour = "black", 
      size = 8
    ),
    axis.text.x = element_text(
      colour = "black", 
      size = 7.5
    ),

    # strip settings
    strip.background = element_rect(
      linetype = "blank"
    ),
    strip.text = element_text(
      face = "italic", 
      colour = "black", 
      size = 10, 
      vjust = 0
    ),

    # plot margin settings
    plot.margin = margin(
      c(0, 0, 0, 0), 
      unit = "cm"
    )
  )
```

Figure C33

```
#ggsave("Raw_tox_comp_plot_Dino.tiff", width = 140, height = 70, units = "mm", dpi=700)
```

For *D. sacculus*, PTX2 is the main toxin that varies. For
*D. acuminata*, it is OA (which is the only toxin it also
produces).

##### Statistical analyses

We will statistically analyse whether the proportions of the three
toxins okadaic acid, PTX2 and C9 change due to treatment. Because *D.
acuminata* only produces okadaic acid, the analyses will only be
performed on *D. sacculus*. To analyse proportion/percentage
data, we first logit (\(ln(p/[1-p])\))
transform the data and then fit uni-factorial ANOVA models using the
transformed toxin proportion as response variable and treatment as
categorical predictor variable. #### OA

```
prop_OA_lm <- lm(logit(prop_OA)~ Treatment, 
                       data = filter(data_both_species,Species == "D. sacculus"))

# Fit a linear model for the logit-transformed proportion of OA toxin
prop_OA_lm <- lm(
  logit(prop_OA) ~ Treatment, 
  data = filter(data_both_species, Species == "D. sacculus") # filter data for D. sacculus
)
```

Assessing the assumptions of normality

```
# Plot a histogram of residuals
hist(
  prop_OA_lm$residuals, # residuals from the linear model
  xlab = "Residuals" # x-axis label
)
```

```
# QQ-plot for residuals
qqPlot(
  prop_OA_lm, # linear model
  grid = FALSE, # do not display grid
  ylab = "Studentised residuals" # y-axis label
)
```

```
## [1]  6 12
```

```
# Perform Shapiro-Wilks test for normality of residuals
shapiro.test(
  prop_OA_lm$residuals # residuals from the linear model
)
```

```
## 
##  Shapiro-Wilk normality test
## 
## data:  prop_OA_lm$residuals
## W = 0.97127, p-value = 0.8763
```

Assessing the assumptions of homoscedasticity

```
# Perform Levene's test for homogeneity of variances
levene_test(
  logit(prop_OA) ~ Treatment, 
  data = filter(data_both_species, Species == "D. sacculus") # filter data for D. sacculus
)
```

```
## # A tibble: 1 × 4
##     df1   df2 statistic     p
##   <int> <int>     <dbl> <dbl>
## 1     4    10     0.372 0.823
```

The transformed proportion data is approximately normal in
distribution and has equal variances, so we will move on to perform the
actual analysis on our fitted linear model.

```
# Perform ANOVA test on the linear model with detailed output and partial eta squared effect size
anova_test(
  prop_OA_lm, # linear model
  effect.size = "pes", # calculate partial eta squared effect size
  detailed = TRUE # provide detailed output
)
```

```
## ANOVA Table (type II tests)
## 
##      Effect   SSn   SSd DFn DFd     F     p p<.05   pes
## 1 Treatment 0.034 0.026   4  10 3.195 0.062       0.561
```

No effect of treatment on OA proportion.

###### PTX2

Linear model:

```
# Fit a linear model for the logit-transformed proportion of PTX2 toxin
prop_PTX2_lm <- lm(
  logit(prop_PTX2) ~ Treatment, 
  data = filter(data_both_species, Species == "D. sacculus") # filter data for D. sacculus
)
```

Assessing the assumptions of normality

```
# Plot a histogram of residuals
hist(
  prop_PTX2_lm$residuals, # residuals from the linear model
  xlab = "Residuals" # x-axis label
)
```

```
# QQ-plot for residuals
qqPlot(
  prop_PTX2_lm, # linear model
  grid = FALSE, # do not display grid
  ylab = "Studentised residuals" # y-axis label
)
```

```
## [1] 10 12
```

```
# Perform Shapiro-Wilks test for normality of residuals
shapiro.test(
  prop_PTX2_lm$residuals # residuals from the linear model
)
```

```
## 
##  Shapiro-Wilk normality test
## 
## data:  prop_PTX2_lm$residuals
## W = 0.94435, p-value = 0.4403
```

Assessing the assumptions of homoscedasticity

```
# Perform Levene's test for homogeneity of variances
levene_test(
  logit(prop_PTX2) ~ Treatment, 
  data = filter(data_both_species, Species == "D. sacculus") # filter data for D. sacculus
)
```

```
## # A tibble: 1 × 4
##     df1   df2 statistic     p
##   <int> <int>     <dbl> <dbl>
## 1     4    10     0.986 0.458
```

The transformed proportion data is approximately normal in
distribution and has equal variances, so we will move on to perform the
actual analysis on our fitted linear model.

```
anova_test(prop_PTX2_lm, effect.size = "pes", detailed = TRUE)
```

```
## ANOVA Table (type II tests)
## 
##      Effect   SSn   SSd DFn DFd     F     p p<.05   pes
## 1 Treatment 0.012 0.042   4  10 0.734 0.589       0.227
```

No effect of treatment on PTX2 proportion.

#### C9

Linear model:

```
# Fit a linear model for the logit-transformed proportion of C9 toxin
prop_C9_lm <- lm(
  logit(prop_C9) ~ Treatment, 
  data = filter(data_both_species, Species == "D. sacculus") # filter data for D. sacculus
)
```

Assessing the assumptions of normality

```
# Plot a histogram of residuals
hist(
  prop_C9_lm$residuals, # residuals from the linear model
  xlab = "Residuals" # x-axis label
)
```

```
# QQ-plot for residuals
qqPlot(
  prop_C9_lm, # linear model
  grid = FALSE, # do not display grid
  ylab = "Studentised residuals" # y-axis label
)
```

```
## [1] 10 12
```

```
# Perform Shapiro-Wilks test for normality of residuals
shapiro.test(
  prop_C9_lm$residuals # residuals from the linear model
)
```

```
## 
##  Shapiro-Wilk normality test
## 
## data:  prop_C9_lm$residuals
## W = 0.9808, p-value = 0.9747
```

Assessing the assumptions of homoscedasticity

```
# Perform Levene's test for homogeneity of variances
levene_test(
  data = filter(data_both_species, Species == "D. sacculus"),
  logit(prop_C9) ~ Treatment
)
```

```
## # A tibble: 1 × 4
##     df1   df2 statistic     p
##   <int> <int>     <dbl> <dbl>
## 1     4    10     0.472 0.756
```

The transformed proportion data is approximately normal in
distribution and has equal variances, so we will move on to perform the
actual analysis on our fitted linear model.

```
# Perform ANOVA test on the linear model with detailed output 
# and partial eta squared effect size

anova_test(
  prop_C9_lm, 
  effect.size = "pes", 
  detailed = TRUE
)
```

```
## ANOVA Table (type II tests)
## 
##      Effect   SSn   SSd DFn DFd     F     p p<.05   pes
## 1 Treatment 0.165 0.484   4  10 0.855 0.523       0.255
```

No effect of treatment on C9 proportion.

##### Final toxin composition plot

```
desc_mean_ci_toxin_all %>%
  dplyr::rename(
    Mean_prop = Mean_prop, # Rename mean proportion
    Lower_prop = LCI_prop, # Rename lower confidence interval
    Upper_prop = HCI_prop  # Rename upper confidence interval
  ) %>%
  mutate(
    Lower_prop = Lower_prop - Mean_prop, # Adjust lower confidence interval
    Upper_prop = Upper_prop - Mean_prop  # Adjust upper confidence interval
  ) %>%
  group_by(Species, Treatment) %>%
  arrange(
    rev(Toxin), # Arrange by toxin type
    by.group = TRUE # Arrange within groups
  ) %>%
  mutate(
    mean2 = cumsum(Mean_prop), # Cumulative sum of mean proportion
    lower = Lower_prop + mean2, # Adjust lower confidence interval
    upper = Upper_prop + mean2  # Adjust upper confidence interval
  ) %>%
  data.frame() %>%
  filter(Species == "D. sacculus") %>% # Filter for D. sacculus
  
  ### Create ggplot
  ggplot(aes(
    y = Mean_prop, # y-axis: mean proportion
    x = Treatment, # x-axis: treatment
    group = Toxin # Group by toxin type
  )) +
  
  ### Stacked bars
  geom_bar(aes(
    y = Mean_prop, 
    fill = Toxin # Fill color by toxin type
  ),
  stat = "identity", # Use raw values
  colour = "black", # Border color of bars
  size = 0.2, # Line width of bar borders
  show.legend = TRUE # Show legend
  ) +
  
  ### CIs
  geom_errorbar(aes(
    y = mean2, # Adjusted mean for positioning error bars
    ymin = lower, # Lower confidence interval
    ymax = upper # Upper confidence interval
  ),
  position = position_dodge(width = 0.35), # Position dodge for error bars
  width = 0.25, # Width of error bars
  size = 0.3 # Line width of error bars
  ) +
  
  # Configure y-axis
  scale_y_continuous(
    n.breaks = 10, # Number of breaks on y-axis
    labels = scales::percent, # Percent labels on y-axis
    minor_breaks = NULL, # Remove minor breaks
    expand = c(0, 0) # Remove expansion at the axis limits
  ) +
  
  # Set the y-axis limits for the viewing window
  coord_cartesian(ylim = c(0, 1)) +
  
  # Fill color palette
  scale_fill_manual(values = c("grey70", "grey90", "grey100")) +
  
  # Theme and axis labels
  theme_classic() +
  theme(
    # Legend settings
    legend.position = "right", # Position the legend
    legend.direction = "vertical", # Arrange legend items vertically
    legend.background = element_rect(
      fill = 'transparent', 
      colour = 'transparent'
    ), # Transparent legend background
    legend.box.background = element_rect(
      fill = 'transparent', 
      colour = 'transparent'
    ), # Transparent legend panel background
    legend.title = element_blank() # Remove legend title
  ) +
  
  # Axis labels
  xlab("") +
  ylab(
    bquote(Proportion~of~total~toxins~("%")) # y-axis label with mathematical annotation
  ) +
  
  # Horizontal line at y = 0
  geom_hline(yintercept = 0) +
  
  # Additional theme modifications
  theme(
    # y-axis title settings
    axis.title.y = element_text(
      vjust = -0.5, 
      size = 10
    ), 

    # Axis text settings
    axis.text.y = element_text(
      colour = "black", 
      size = 8
    ),
    axis.text.x = element_text(
      colour = "black", 
      size = 8
    ),

    # Strip settings
    strip.background = element_rect(
      linetype = "blank"
    ),
    strip.text = element_text(
      face = "italic", 
      colour = "black", 
      size = 10, 
      vjust = 0
    ),

    # Legend text settings
    legend.text = element_text(
      colour = "black", 
      size = 8
    ),
    legend.box.margin = margin(
      c(0, -0.2, 0, -0.7), 
      unit = "cm"
    ),
    legend.key.size = unit(0.4, "cm"), # Size of legend keys

    # Plot margin settings
    plot.margin = margin(
      c(0.15, 0.15, -0.2, 0), 
      unit = "cm"
    )
  )
```

Figure C34. Composition of toxin congeners (white = Okadaic acid, light
grey = PTX2 and dark grey = C9) for *Dinophysis sacculus* after
68.5 hours of exposure to three concentrations of chemical cues from
copepods (copepodamides), a positive control (one live *Acartia*
sp.) or a control treatment. Bars are the mean of three replicates (n =
3) and error bars show the 95% confidence intervals of the means. N.B
only the lower confidence limit is shown for C9 means to avoid a y-axis
> 100%.

```
#ggsave(path = "figures_manuscript", "Prop_tox_comp_plot_Dino.tiff", width = 85, height = 90, units = "mm", dpi=700)
```

### Additional plots

#### Metabolomics PCA plot

##### *D. sacculus*

###### Data

```
# Import data for PCA analysis
pca_sacculus <- read_delim(
  "pca_score_sacculus.csv", # file path
  delim = ";", # delimiter
  escape_double = FALSE, # do not escape double quotes
  col_types = cols(
    Treatment = col_factor(levels = c("Control", "Copepod", "1 nM", 
                                      "5 nM", "10 nM")), # factor column for Treatment
    Ion_mode = col_factor(levels = c("pos", "neg")), # factor column for Ion_mode
    PCA1 = col_number(), # numeric column for PCA1
    PCA2 = col_number()  # numeric column for PCA2
  ),
  trim_ws = TRUE # trim whitespace
)

# Calculate centroids for MS+ ion mode
centroids_sacculus_pos <- aggregate(
  cbind(PCA1, PCA2) ~ Treatment, # columns to aggregate and group by Treatment
  data = filter(pca_sacculus, Ion_mode == "pos"), # filter data for positive ion mode
  FUN = mean # calculate mean for each group
)

# Calculate centroids for MS- ion mode
centroids_sacculus_neg <- aggregate(
  cbind(PCA1, PCA2) ~ Treatment, # columns to aggregate and group by Treatment
  data = filter(pca_sacculus, Ion_mode == "neg"), # filter data for negative ion mode
  FUN = mean # calculate mean for each group
)
```

###### Plot for +MS

```
# Create the PCA plot for positive ion mode
pca_sacculus %>%
  filter(Ion_mode == "pos") %>% # filter for positive ion mode
  ggplot(aes(
    x = PCA1, # x-axis: PCA1
    y = PCA2  # y-axis: PCA2
  )) +
  
  # Points
  geom_point(aes(colour = Treatment), # color points by Treatment
             size = 3, # size of points
             shape = c(15, 15, 15, 16, 16, 16, 17, 17, 17, 17, 17, 17, 17, 17, 17), # shapes of points
             alpha = 0.4, # transparency of points
             show.legend = FALSE # do not show legend for points
  ) +
  
  # Centroid locations
  geom_point(aes(fill = Treatment), # fill color by Treatment
             data = centroids_sacculus_pos, # data for centroids
             size = 4, # size of centroids
             shape = c(22, 21, 24, 24, 24), # shapes of centroids
             colour = "black", # border color of centroids
             alpha = 0.9, # transparency of centroids
             show.legend = FALSE # do not show legend for centroids
  ) +

  # Set theme
  theme_classic() +
  
  # Configure y-axis
  scale_y_continuous(
    limits = c(-20, 20), # set y-axis limits
    expand = c(0, 0) # remove expansion at the axis limits
  ) +
  
  # Configure x-axis
  scale_x_continuous(
    limits = c(-12, 25), # set x-axis limits
    expand = c(0, 0) # remove expansion at the axis limits
  ) +
  
  # Color palette for points and centroids
  scale_color_manual(values = custom_wes_palette) +
  scale_fill_manual(values = custom_wes_palette) +
  
  # Adjust legend symbols
  guides(colour = guide_legend(
    override.aes = list(
      alpha = 0.9, # set transparency
      size = 3, # set size
      shape = c(15, 16, 17, 17, 17) # set shapes
    )
  )) +
  
  # Axis labels and title
  xlab("PCA1 (39.6%)") +
  ylab("PCA2 (21.9%)") +
  ggtitle("D. sacculus") +
  
  # Additional theme modifications
  theme(
    plot.title = element_text(
      face = "italic", # italicize title
      hjust = 0.5, # center align title
      size = 10 # font size of title
    ),
    axis.title.y = element_text(
      vjust = 0, # vertical adjustment for y-axis title
      size = 8 # font size of y-axis title
    ),
    axis.title.x = element_text(
      size = 8 # font size of x-axis title
    ),
    axis.text.y = element_text(
      colour = "black", # color of y-axis text
      size = 8 # font size of y-axis text
    ),
    axis.text.x = element_text(
      colour = "black", # color of x-axis text
      size = 8 # font size of x-axis text
    ),
    legend.text = element_text(
      colour = "black", # color of legend text
      size = 7, # font size of legend text
      margin = margin(r = -2, unit = "pt") # margin for legend text
    ),
    legend.title = element_blank(), # remove legend title
    legend.background = element_rect(
      fill = 'transparent', # transparent background
      colour = 'transparent' # transparent border
    ),
    legend.box.background = element_rect(
      fill = 'transparent', # transparent legend panel
      colour = 'transparent' # transparent border
    ),
    legend.position = c(0.14, 0.86), # position legend
    legend.direction = "vertical", # vertical legend
    legend.key.size = unit(10, 'pt') # size of legend keys
  )
```

Figure C35. PCA score plot of LC-HRMS derived metabolomic profiles in
+MS mode

```
#ggsave(path = "figures_manuscript","PCA_sacculus_pos.tiff", width = 70, height = 70, units = "mm", dpi=700)
```

###### Plot for -MS

```
# Create the nMDS plot for negative ion mode
pca_sacculus %>%
  filter(Ion_mode == "neg") %>% # filter for negative ion mode
  ggplot(aes(
    x = PCA1, # x-axis: PCA1
    y = PCA2  # y-axis: PCA2
  )) +
  
  # Points
  geom_point(aes(colour = Treatment), # color points by Treatment
             size = 3, # size of points
             shape = c(15, 15, 15, 16, 16, 16, 17, 17, 17, 17, 17, 17, 17, 17, 17), # shapes of points
             alpha = 0.4, # transparency of points
             show.legend = TRUE # show legend for points
  ) +
  
  # Centroid locations
  geom_point(aes(fill = Treatment), # fill color by Treatment
             data = centroids_sacculus_neg, # data for centroids
             size = 4, # size of centroids
             shape = c(22, 21, 24, 24, 24), # shapes of centroids
             colour = "black", # border color of centroids
             alpha = 0.9, # transparency of centroids
             show.legend = FALSE # do not show legend for centroids
  ) +

  # Set theme
  theme_classic() +
  
  # Configure y-axis
  scale_y_continuous(
    limits = c(-10, 15), # set y-axis limits
    expand = c(0, 0) # remove expansion at the axis limits
  ) +
  
  # Configure x-axis
  scale_x_continuous(
    limits = c(-20, 20), # set x-axis limits
    expand = c(0, 0) # remove expansion at the axis limits
  ) +
  
  # Color palette for points and centroids
  scale_color_manual(values = custom_wes_palette) +
  scale_fill_manual(values = custom_wes_palette) +
  
  # Adjust legend symbols
  guides(colour = guide_legend(
    override.aes = list(
      alpha = 0.9, # set transparency
      size = 3, # set size
      shape = c(15, 16, 17, 17, 17) # set shapes
    )
  )) +
  
  # Axis labels and title
  xlab("PCA1 (31.8%)") +
  ylab("PCA2 (24.5%)") +
  ggtitle("D. sacculus") +
  
  # Additional theme modifications
  theme(
    plot.title = element_text(
      face = "italic", # italicize title
      hjust = 0.5, # center align title
      size = 10 # font size of title
    ),
    axis.title.y = element_text(
      vjust = 0, # vertical adjustment for y-axis title
      size = 8 # font size of y-axis title
    ),
    axis.title.x = element_text(
      size = 8 # font size of x-axis title
    ),
    axis.text.y = element_text(
      colour = "black", # color of y-axis text
      size = 8 # font size of y-axis text
    ),
    axis.text.x = element_text(
      colour = "black", # color of x-axis text
      size = 8 # font size of x-axis text
    ),
    legend.text = element_text(
      colour = "black", # color of legend text
      size = 7, # font size of legend text
      margin = margin(r = -2, unit = "pt") # margin for legend text
    ),
    legend.title = element_blank(), # remove legend title
    legend.background = element_rect(
      fill = 'transparent', # transparent background
      colour = 'transparent' # transparent border
    ),
    legend.box.background = element_rect(
      fill = 'transparent', # transparent legend panel
      colour = 'transparent' # transparent border
    ),
    legend.position = c(0.15, 0.20), # position legend
    legend.direction = "vertical", # vertical legend
    legend.key.size = unit(10, 'pt') # size of legend keys
  )
```

Figure C36. PCA score plot of LC-HRMS derived metabolomic profiles in
-MS mode

```
#ggsave(path = "figures_manuscript","PCA_sacculus_neg.tiff", width = 70, height = 70, units = "mm", dpi=700)
```

##### *D. acuminata*

###### Data

```
#Import data
pca_acuminata <- 
  read_delim("pca_score_acuminata.csv",
             delim = ";", 
             escape_double = FALSE, 
             col_types = cols(Treatment = 
                                col_factor(
                                  levels = c("Control", 
                                             "Copepod", 
                                             "1 nM",
                                             "5 nM",
                                             "10 nM")),
                              Ion_mode =
                                col_factor(
                                  levels = c("pos", "neg")),
                                
                              PCA1 = col_number(), 
                              PCA2 = col_number()), 
             trim_ws = TRUE)


#Calculate centroids in MS-
centroids_acuminata_neg <- aggregate(cbind(PCA1, PCA2) ~ Treatment, 
                                   data = filter(pca_acuminata, Ion_mode == "neg"), 
                                FUN = mean)

#Calculate centroids in MS+
centroids_acuminata_pos <- aggregate(cbind(PCA1, PCA2) ~ Treatment, 
                                   data = filter(pca_acuminata, Ion_mode == "pos"), 
                                FUN = mean)
```

###### Plot for -MS

```
# Create the PCA plot for negative ion mode
pca_acuminata %>%
  filter(Ion_mode == "neg") %>% # filter for negative ion mode
  ggplot(aes(
    x = PCA1, # x-axis: PCA1
    y = PCA2  # y-axis: PCA2
  )) +
  
  # Points
  geom_point(aes(colour = Treatment), # color points by Treatment
             size = 3, # size of points
             shape = c(15, 15, 15, 16, 16, 16, 17, 17, 17, 17, 17, 17, 17, 17, 17), # shapes of points
             alpha = 0.4, # transparency of points
             show.legend = TRUE # show legend for points
  ) +
  
  # Centroid locations
  geom_point(aes(fill = Treatment), # fill color by Treatment
             data = centroids_acuminata_neg, # data for centroids
             size = 4, # size of centroids
             shape = c(22, 21, 24, 24, 24), # shapes of centroids
             colour = "black", # border color of centroids
             alpha = 0.9, # transparency of centroids
             show.legend = FALSE # do not show legend for centroids
  ) +

  # Set theme
  theme_classic() +
  
  # Configure y-axis
  scale_y_continuous(
    limits = c(-15, 15), # set y-axis limits
    expand = c(0, 0) # remove expansion at the axis limits
  ) +
  
  # Configure x-axis
  scale_x_continuous(
    limits = c(-8, 25), # set x-axis limits
    expand = c(0, 0) # remove expansion at the axis limits
  ) +
  
  # Color palette for points and centroids
  scale_color_manual(values = custom_wes_palette) +
  scale_fill_manual(values = custom_wes_palette) +
  
  # Adjust legend symbols
  guides(colour = guide_legend(
    override.aes = list(
      alpha = 0.9, # set transparency
      size = 3, # set size
      shape = c(15, 16, 17, 17, 17) # set shapes
    )
  )) +
  
  # Axis labels and title
  xlab("PCA1 (30.7%)") +
  ylab("PCA2 (17.8%)") +
  ggtitle("D. acuminata") +
  
  # Additional theme modifications
  theme(
    plot.title = element_text(
      face = "italic", # italicize title
      hjust = 0.5, # center align title
      size = 10 # font size of title
    ),
    axis.title.y = element_text(
      vjust = 0, # vertical adjustment for y-axis title
      size = 8 # font size of y-axis title
    ),
    axis.title.x = element_text(
      size = 8 # font size of x-axis title
    ),
    axis.text.y = element_text(
      colour = "black", # color of y-axis text
      size = 8 # font size of y-axis text
    ),
    axis.text.x = element_text(
      colour = "black", # color of x-axis text
      size = 8 # font size of x-axis text
    ),
    legend.text = element_text(
      colour = "black", # color of legend text
      size = 7, # font size of legend text
      margin = margin(r = -2, unit = "pt") # margin for legend text
    ),
    legend.title = element_blank(), # remove legend title
    legend.background = element_rect(
      fill = 'transparent', # transparent background
      colour = 'transparent' # transparent border
    ),
    legend.box.background = element_rect(
      fill = 'transparent', # transparent legend panel
      colour = 'transparent' # transparent border
    ),
    legend.position = c(0.85, 0.88), # position legend
    legend.direction = "vertical", # vertical legend
    legend.key.size = unit(10, 'pt') # size of legend keys
  )
```

Figure C37. PCA score plot of LC-HRMS derived metabolomic profiles in
-MS mode

```
#ggsave(path = "figures_manuscript", "PCA_acuminata_neg.tiff", width = 70, height = 70, units = "mm", dpi=700)
```

###### Plot for +MS

```
# Create the PCA plot for positive ion mode
pca_acuminata %>%
  filter(Ion_mode == "pos") %>% # filter for positive ion mode
  ggplot(aes(
    x = PCA1, # x-axis: PCA1
    y = PCA2  # y-axis: PCA2
  )) +
  
  # Points
  geom_point(aes(colour = Treatment), # color points by Treatment
             size = 3, # size of points
             shape = c(15, 15, 15, 16, 16, 16, 17, 17, 17, 17, 17, 17, 17, 17, 17), # shapes of points
             alpha = 0.4, # transparency of points
             show.legend = FALSE # do not show legend for points
  ) +
  
  # Centroid locations
  geom_point(aes(fill = Treatment), # fill color by Treatment
             data = centroids_acuminata_pos, # data for centroids
             size = 4, # size of centroids
             shape = c(22, 21, 24, 24, 24), # shapes of centroids
             colour = "black", # border color of centroids
             alpha = 0.9, # transparency of centroids
             show.legend = FALSE # do not show legend for centroids
  ) +

  # Set theme
  theme_classic() +
  
  # Configure y-axis
  scale_y_continuous(
    limits = c(-30, 20), # set y-axis limits
    expand = c(0, 0) # remove expansion at the axis limits
  ) +
  
  # Configure x-axis
  scale_x_continuous(
    limits = c(-40, 30), # set x-axis limits
    expand = c(0, 0) # remove expansion at the axis limits
  ) +
  
  # Color palette for points and centroids
  scale_color_manual(values = custom_wes_palette) +
  scale_fill_manual(values = custom_wes_palette) +
  
  # Adjust legend symbols
  guides(colour = guide_legend(
    override.aes = list(
      alpha = 0.9, # set transparency
      size = 3, # set size
      shape = c(15, 16, 17, 17, 17) # set shapes
    )
  )) +
  
  # Axis labels and title
  xlab("PCA1 (31.2%)") +
  ylab("PCA2 (17.0%)") +
  ggtitle("D. acuminata") +
  
  # Additional theme modifications
  theme(
    plot.title = element_text(
      face = "italic", # italicize title
      hjust = 0.5, # center align title
      size = 10 # font size of title
    ),
    axis.title.y = element_text(
      vjust = 0, # vertical adjustment for y-axis title
      size = 8 # font size of y-axis title
    ),
    axis.title.x = element_text(
      size = 8 # font size of x-axis title
    ),
    axis.text.y = element_text(
      colour = "black", # color of y-axis text
      size = 8 # font size of y-axis text
    ),
    axis.text.x = element_text(
      colour = "black", # color of x-axis text
      size = 8 # font size of x-axis text
    ),
    legend.text = element_text(
      colour = "black", # color of legend text
      size = 7, # font size of legend text
      margin = margin(r = -2, unit = "pt") # margin for legend text
    ),
    legend.title = element_blank(), # remove legend title
    legend.background = element_rect(
      fill = 'transparent', # transparent background
      colour = 'transparent' # transparent border
    ),
    legend.box.background = element_rect(
      fill = 'transparent', # transparent legend panel
      colour = 'transparent' # transparent border
    ),
    legend.position = c(0.15, 0.40), # position legend
    legend.direction = "vertical", # vertical legend
    legend.key.size = unit(10, 'pt') # size of legend keys
  )
```

Figure C38. PCA score plot of LC-HRMS derived metabolomic profiles in
+MS mode

```
#ggsave(path = "figures_manuscript", "PCA_acuminata_pos.tiff", width = 70, height = 70, units = "mm", dpi=700)
```

### R session info

```
devtools::session_info()
```

```
## ─ Session info ───────────────────────────────────────────────────────────────
##  setting  value
##  version  R version 4.4.1 (2024-06-14 ucrt)
##  os       Windows 11 x64 (build 22631)
##  system   x86_64, mingw32
##  ui       RTerm
##  language (EN)
##  collate  Swedish_Sweden.utf8
##  ctype    Swedish_Sweden.utf8
##  tz       Europe/Stockholm
##  date     2024-07-16
##  pandoc   3.1.11 @ C:/Program Files/RStudio/resources/app/bin/quarto/bin/tools/ (via rmarkdown)
## 
## ─ Packages ───────────────────────────────────────────────────────────────────
##  ! package      * version date (UTC) lib source
##    abind          1.4-5   2016-07-21 [1] CRAN (R 4.4.0)
##    AICcmodavg   * 2.3-3   2023-11-16 [1] CRAN (R 4.4.0)
##    backports      1.5.0   2024-05-23 [1] CRAN (R 4.4.0)
##    base64enc      0.1-3   2015-07-28 [1] CRAN (R 4.4.0)
##    bit            4.0.5   2022-11-15 [1] CRAN (R 4.4.0)
##    bit64          4.0.5   2020-08-30 [1] CRAN (R 4.4.0)
##    boot           1.3-30  2024-02-26 [1] CRAN (R 4.4.0)
##    broom        * 1.0.6   2024-05-17 [1] CRAN (R 4.4.1)
##    bslib          0.7.0   2024-03-29 [1] CRAN (R 4.4.0)
##    cachem         1.1.0   2024-05-16 [1] CRAN (R 4.4.0)
##    car          * 3.1-2   2023-03-30 [1] CRAN (R 4.4.0)
##    carData      * 3.0-5   2022-01-06 [1] CRAN (R 4.4.0)
##    cellranger     1.1.0   2016-07-27 [1] CRAN (R 4.4.0)
##    checkmate      2.3.1   2023-12-04 [1] CRAN (R 4.4.0)
##    chunkhooks   * 0.0.1   2020-08-05 [1] CRAN (R 4.4.0)
##    class          7.3-22  2023-05-03 [1] CRAN (R 4.4.0)
##    cli            3.6.1   2023-03-23 [1] CRAN (R 4.2.3)
##    cluster        2.1.6   2023-12-01 [2] CRAN (R 4.4.1)
##    codetools      0.2-20  2024-03-31 [1] CRAN (R 4.4.0)
##    coin           1.4-3   2023-09-27 [1] CRAN (R 4.4.0)
##  D colorspace     2.1-0   2023-01-23 [1] CRAN (R 4.3.2)
##    crayon         1.5.3   2024-06-20 [1] CRAN (R 4.4.1)
##    data.table     1.15.4  2024-03-30 [1] CRAN (R 4.4.0)
##    DescTools    * 0.99.54 2024-02-03 [1] CRAN (R 4.4.0)
##    devtools     * 2.4.5   2022-10-11 [1] CRAN (R 4.4.1)
##    digest         0.6.36  2024-06-23 [1] CRAN (R 4.4.1)
##    dplyr        * 1.1.4   2023-11-17 [1] CRAN (R 4.4.0)
##    e1071          1.7-14  2023-12-06 [1] CRAN (R 4.4.0)
##  D ellipsis       0.3.2   2021-04-29 [1] CRAN (R 4.4.1)
##    evaluate       0.24.0  2024-06-10 [1] CRAN (R 4.4.0)
##    Exact          3.2     2022-09-25 [1] CRAN (R 4.4.0)
##    expm           0.999-9 2024-01-11 [1] CRAN (R 4.4.0)
##    fansi          1.0.6   2023-12-08 [1] CRAN (R 4.4.0)
##    farver         2.1.2   2024-05-13 [1] CRAN (R 4.4.0)
##    fastmap        1.2.0   2024-05-15 [1] CRAN (R 4.4.0)
##    forcats      * 1.0.0   2023-01-29 [1] CRAN (R 4.4.0)
##    foreign        0.8-87  2024-06-26 [1] CRAN (R 4.4.1)
##    Formula        1.2-5   2023-02-24 [1] CRAN (R 4.4.0)
##    fs             1.6.4   2024-04-25 [1] CRAN (R 4.4.0)
##    generics       0.1.3   2022-07-05 [1] CRAN (R 4.4.0)
##    ggplot2      * 3.5.1   2024-04-23 [1] CRAN (R 4.4.0)
##    ggpmisc      * 0.6.0   2024-06-28 [1] CRAN (R 4.4.1)
##    ggpp         * 0.5.8-1 2024-07-01 [1] CRAN (R 4.4.1)
##    ggpubr       * 0.6.0   2023-02-10 [1] CRAN (R 4.4.1)
##    ggsignif       0.6.4   2022-10-13 [1] CRAN (R 4.4.0)
##    ggthemes     * 5.1.0   2024-02-10 [1] CRAN (R 4.4.1)
##    gld            2.6.6   2022-10-23 [1] CRAN (R 4.4.0)
##    glue           1.6.2   2022-02-24 [1] CRAN (R 4.1.3)
##    gridExtra      2.3     2017-09-09 [1] CRAN (R 4.4.0)
##    gtable         0.3.5   2024-04-22 [1] CRAN (R 4.4.0)
##    highr          0.11    2024-05-26 [1] CRAN (R 4.4.0)
##    Hmisc        * 5.1-3   2024-05-28 [1] CRAN (R 4.4.1)
##    hms            1.1.3   2023-03-21 [1] CRAN (R 4.4.1)
##    htmlTable      2.4.2   2023-10-29 [1] CRAN (R 4.4.1)
##    htmltools      0.5.8.1 2024-04-04 [1] CRAN (R 4.4.0)
##    htmlwidgets    1.6.4   2023-12-06 [1] CRAN (R 4.4.1)
##    httpuv         1.6.15  2024-03-26 [1] CRAN (R 4.4.0)
##    httr           1.4.7   2023-08-15 [1] CRAN (R 4.4.1)
##    jquerylib      0.1.4   2021-04-26 [1] CRAN (R 4.4.1)
##    jsonlite       1.8.8   2023-12-04 [1] CRAN (R 4.4.1)
##    kableExtra   * 1.4.0   2024-01-24 [1] CRAN (R 4.4.1)
##    knitr        * 1.48    2024-07-07 [1] CRAN (R 4.4.1)
##    labeling       0.4.3   2023-08-29 [1] CRAN (R 4.4.0)
##    later          1.3.2   2023-12-06 [1] CRAN (R 4.3.2)
##    lattice      * 0.22-6  2024-03-20 [2] CRAN (R 4.4.1)
##  D libcoin        1.0-10  2023-09-27 [1] CRAN (R 4.4.1)
##    lifecycle      1.0.4   2023-11-07 [1] CRAN (R 4.4.1)
##    lmom           3.0     2023-08-29 [1] CRAN (R 4.4.0)
##    lmtest         0.9-40  2022-03-21 [1] CRAN (R 4.4.1)
##    lubridate    * 1.9.3   2023-09-27 [1] CRAN (R 4.4.1)
##    magrittr       2.0.3   2022-03-30 [1] CRAN (R 4.1.3)
##    MASS         * 7.3-60  2023-05-04 [1] CRAN (R 4.3.2)
##    Matrix         1.7-0   2024-03-22 [1] CRAN (R 4.4.1)
##    MatrixModels   0.5-3   2023-11-06 [1] CRAN (R 4.4.1)
##    matrixStats    1.3.0   2024-04-11 [1] CRAN (R 4.4.0)
##    memoise        2.0.1   2021-11-26 [1] CRAN (R 4.4.1)
##  D mime           0.12    2021-09-28 [1] CRAN (R 4.3.1)
##    miniUI         0.1.1.1 2018-05-18 [1] CRAN (R 4.4.1)
##    modeltools     0.2-23  2020-03-05 [1] CRAN (R 4.4.0)
##    multcomp     * 1.4-25  2023-06-20 [1] CRAN (R 4.4.1)
##    multcompView   0.1-10  2024-03-08 [1] CRAN (R 4.4.0)
##    munsell        0.5.1   2024-04-01 [1] CRAN (R 4.4.0)
##    mvtnorm      * 1.2-5   2024-05-21 [1] CRAN (R 4.4.1)
##    nlme           3.1-165 2024-06-06 [1] CRAN (R 4.4.1)
##    nnet           7.3-19  2023-05-03 [2] CRAN (R 4.4.1)
##    nortest        1.0-4   2015-07-30 [1] CRAN (R 4.4.0)
##    pacman       * 0.5.1   2019-03-11 [1] CRAN (R 4.4.1)
##    pillar         1.9.0   2023-03-22 [1] CRAN (R 4.3.2)
##    pkgbuild       1.4.4   2024-03-17 [1] CRAN (R 4.4.0)
##    pkgconfig      2.0.3   2019-09-22 [1] CRAN (R 4.3.2)
##    pkgload        1.4.0   2024-06-28 [1] CRAN (R 4.4.1)
##  D plyr         * 1.8.9   2023-10-02 [1] CRAN (R 4.3.2)
##    polynom        1.4-1   2022-04-11 [1] CRAN (R 4.3.2)
##    profvis        0.3.8   2023-05-02 [1] CRAN (R 4.3.2)
##    promises       1.3.0   2024-04-05 [1] CRAN (R 4.4.0)
##  D proxy          0.4-27  2022-06-09 [1] CRAN (R 4.3.2)
##    purrr        * 1.0.2   2023-08-10 [1] CRAN (R 4.4.0)
##    quantreg       5.98    2024-05-26 [1] CRAN (R 4.4.0)
##    R6             2.5.1   2021-08-19 [1] CRAN (R 4.3.2)
##    rcompanion   * 2.4.36  2024-05-27 [1] CRAN (R 4.4.0)
##    Rcpp           1.0.12  2024-01-09 [1] CRAN (R 4.4.0)
##    readr        * 2.1.5   2024-01-10 [1] CRAN (R 4.4.1)
##    readxl         1.4.3   2023-07-06 [1] CRAN (R 4.3.2)
##    remotes        2.5.0   2024-03-17 [1] CRAN (R 4.4.0)
##    rlang          1.1.4   2024-06-04 [1] CRAN (R 4.4.1)
##    rmarkdown      2.27    2024-05-17 [1] CRAN (R 4.4.0)
##    Rmisc        * 1.5.1   2022-05-02 [1] CRAN (R 4.3.2)
##    rootSolve      1.8.2.4 2023-09-21 [1] CRAN (R 4.3.1)
##    rpart          4.1.23  2023-12-05 [2] CRAN (R 4.4.1)
##    rstatix      * 0.7.2   2023-02-01 [1] CRAN (R 4.4.1)
##    rstudioapi     0.16.0  2024-03-24 [1] CRAN (R 4.4.0)
##    sandwich       3.1-0   2023-12-11 [1] CRAN (R 4.3.2)
##    sass           0.4.9   2024-03-15 [1] CRAN (R 4.4.0)
##    scales         1.3.0   2023-11-28 [1] CRAN (R 4.3.2)
##    sessioninfo    1.2.2   2021-12-06 [1] CRAN (R 4.3.2)
##    shiny          1.8.1.1 2024-04-02 [1] CRAN (R 4.4.0)
##    SparseM        1.84    2024-06-25 [1] CRAN (R 4.4.1)
##    stringi        1.7.12  2023-01-11 [1] CRAN (R 4.2.2)
##    stringr      * 1.5.1   2023-11-14 [1] CRAN (R 4.3.2)
##    survival     * 3.7-0   2024-06-05 [1] CRAN (R 4.4.1)
##    svglite        2.1.3   2023-12-08 [1] CRAN (R 4.3.2)
##    systemfonts    1.1.0   2024-05-15 [1] CRAN (R 4.4.0)
##    TH.data      * 1.1-2   2023-04-17 [1] CRAN (R 4.3.2)
##    tibble       * 3.2.1   2023-03-20 [1] CRAN (R 4.2.3)
##    tidyr        * 1.3.1   2024-01-24 [1] CRAN (R 4.4.0)
##    tidyselect     1.2.1   2024-03-11 [1] CRAN (R 4.4.0)
##    tidyverse    * 2.0.0   2023-02-22 [1] CRAN (R 4.4.1)
##    timechange     0.3.0   2024-01-18 [1] CRAN (R 4.4.0)
##  D tzdb           0.4.0   2023-05-12 [1] CRAN (R 4.3.2)
##    unmarked       1.4.1   2024-01-09 [1] CRAN (R 4.4.0)
##    urlchecker     1.0.1   2021-11-30 [1] CRAN (R 4.3.2)
##    usethis      * 2.2.3   2024-02-19 [1] CRAN (R 4.4.0)
##    utf8           1.2.4   2023-10-22 [1] CRAN (R 4.4.0)
##    vctrs          0.6.4   2023-10-12 [1] CRAN (R 4.2.3)
##    VGAM           1.1-11  2024-05-15 [1] CRAN (R 4.4.0)
##    viridis      * 0.6.5   2024-01-29 [1] CRAN (R 4.4.1)
##    viridisLite  * 0.4.2   2023-05-02 [1] CRAN (R 4.3.2)
##    vroom          1.6.5   2023-12-05 [1] CRAN (R 4.3.2)
##    wesanderson  * 0.3.7   2023-10-31 [1] CRAN (R 4.3.3)
##    withr          3.0.0   2024-01-16 [1] CRAN (R 4.4.0)
##    xfun           0.45    2024-06-16 [1] CRAN (R 4.4.1)
##    xml2           1.3.6   2023-12-04 [1] CRAN (R 4.4.0)
##    xtable         1.8-4   2019-04-21 [1] CRAN (R 4.3.2)
##    yaml           2.3.9   2024-07-05 [1] CRAN (R 4.4.1)
##    zoo            1.8-12  2023-04-13 [1] CRAN (R 4.3.2)
## 
##  [1] C:/Users/xpoumi/OneDrive/Dokument/R/win-library/4.0
##  [2] C:/Program Files/R/R-4.4.1/library
## 
##  D ── DLL MD5 mismatch, broken installation.
## 
## ──────────────────────────────────────────────────────────────────────────────
```

### Package references

```
## To cite package 'devtools' in publications use:
## 
##   Wickham H, Hester J, Chang W, Bryan J (2022). _devtools: Tools to
##   Make Developing R Packages Easier_. R package version 2.4.5,
##   <https://CRAN.R-project.org/package=devtools>.
## 
## A BibTeX entry for LaTeX users is
## 
##   @Manual{,
##     title = {devtools: Tools to Make Developing R Packages Easier},
##     author = {Hadley Wickham and Jim Hester and Winston Chang and Jennifer Bryan},
##     year = {2022},
##     note = {R package version 2.4.5},
##     url = {https://CRAN.R-project.org/package=devtools},
##   }
```

```
## To cite package 'readr' in publications use:
## 
##   Wickham H, Hester J, Bryan J (2024). _readr: Read Rectangular Text
##   Data_. R package version 2.1.5,
##   <https://CRAN.R-project.org/package=readr>.
## 
## A BibTeX entry for LaTeX users is
## 
##   @Manual{,
##     title = {readr: Read Rectangular Text Data},
##     author = {Hadley Wickham and Jim Hester and Jennifer Bryan},
##     year = {2024},
##     note = {R package version 2.1.5},
##     url = {https://CRAN.R-project.org/package=readr},
##   }
```

```
## To cite package 'ggpubr' in publications use:
## 
##   Kassambara A (2023). _ggpubr: 'ggplot2' Based Publication Ready
##   Plots_. R package version 0.6.0,
##   <https://CRAN.R-project.org/package=ggpubr>.
## 
## A BibTeX entry for LaTeX users is
## 
##   @Manual{,
##     title = {ggpubr: 'ggplot2' Based Publication Ready Plots},
##     author = {Alboukadel Kassambara},
##     year = {2023},
##     note = {R package version 0.6.0},
##     url = {https://CRAN.R-project.org/package=ggpubr},
##   }
```

```
## Please cite the multcomp package by the following reference:
## 
##   Hothorn T, Bretz F, Westfall P (2008). "Simultaneous Inference in
##   General Parametric Models." _Biometrical Journal_, *50*(3), 346-363.
## 
## A BibTeX entry for LaTeX users is
## 
##   @Article{,
##     title = {Simultaneous Inference in General Parametric Models},
##     author = {Torsten Hothorn and Frank Bretz and Peter Westfall},
##     journal = {Biometrical Journal},
##     year = {2008},
##     volume = {50},
##     number = {3},
##     pages = {346--363},
##   }
```

```
## To cite package 'rstatix' in publications use:
## 
##   Kassambara A (2023). _rstatix: Pipe-Friendly Framework for Basic
##   Statistical Tests_. R package version 0.7.2,
##   <https://CRAN.R-project.org/package=rstatix>.
## 
## A BibTeX entry for LaTeX users is
## 
##   @Manual{,
##     title = {rstatix: Pipe-Friendly Framework for Basic Statistical Tests},
##     author = {Alboukadel Kassambara},
##     year = {2023},
##     note = {R package version 0.7.2},
##     url = {https://CRAN.R-project.org/package=rstatix},
##   }
```

```
## To cite package 'broom' in publications use:
## 
##   Robinson D, Hayes A, Couch S (2024). _broom: Convert Statistical
##   Objects into Tidy Tibbles_. R package version 1.0.6,
##   <https://CRAN.R-project.org/package=broom>.
## 
## A BibTeX entry for LaTeX users is
## 
##   @Manual{,
##     title = {broom: Convert Statistical Objects into Tidy Tibbles},
##     author = {David Robinson and Alex Hayes and Simon Couch},
##     year = {2024},
##     note = {R package version 1.0.6},
##     url = {https://CRAN.R-project.org/package=broom},
##   }
```

```
## To cite package 'rcompanion' in publications use:
## 
##   Mangiafico, Salvatore S. (2024). rcompanion: Functions to Support
##   Extension Education Program Evaluation. version 2.4.36. Rutgers
##   Cooperative Extension. New Brunswick, New Jersey.
##   https://CRAN.R-project.org/package=rcompanion
## 
## A BibTeX entry for LaTeX users is
## 
##   @Manual{,
##     title = {{rcompanion}: Functions to Support Extension Education Program Evaluation},
##     author = {Salvatore S. Mangiafico},
##     organization = {Rutgers Cooperative Extension},
##     address = {New Brunswick, New Jersey},
##     note = {version 2.4.36},
##     year = {2024},
##     url = {https://CRAN.R-project.org/package=rcompanion/},
##   }
```

```
## To cite the car package in publications use:
## 
##   Fox J, Weisberg S (2019). _An R Companion to Applied Regression_,
##   Third edition. Sage, Thousand Oaks CA.
##   <https://socialsciences.mcmaster.ca/jfox/Books/Companion/>.
## 
## A BibTeX entry for LaTeX users is
## 
##   @Book{,
##     title = {An {R} Companion to Applied Regression},
##     edition = {Third},
##     author = {John Fox and Sanford Weisberg},
##     year = {2019},
##     publisher = {Sage},
##     address = {Thousand Oaks {CA}},
##     url = {https://socialsciences.mcmaster.ca/jfox/Books/Companion/},
##   }
```

```
## To cite package 'DescTools' in publications use:
## 
##   Signorell A (2024). _DescTools: Tools for Descriptive Statistics_. R
##   package version 0.99.54,
##   <https://CRAN.R-project.org/package=DescTools>.
## 
## A BibTeX entry for LaTeX users is
## 
##   @Manual{,
##     title = {DescTools: Tools for Descriptive Statistics},
##     author = {Andri Signorell},
##     year = {2024},
##     note = {R package version 0.99.54},
##     url = {https://CRAN.R-project.org/package=DescTools},
##   }
```

```
## To cite package 'Rmisc' in publications use:
## 
##   Hope RM (2022). _Rmisc: Ryan Miscellaneous_. R package version 1.5.1,
##   <https://CRAN.R-project.org/package=Rmisc>.
## 
## A BibTeX entry for LaTeX users is
## 
##   @Manual{,
##     title = {Rmisc: Ryan Miscellaneous},
##     author = {Ryan M. Hope},
##     year = {2022},
##     note = {R package version 1.5.1},
##     url = {https://CRAN.R-project.org/package=Rmisc},
##   }
## 
## ATTENTION: This citation information has been auto-generated from the
## package DESCRIPTION file and may need manual editing, see
## 'help("citation")'.
```

```
## To cite viridis/viridisLite in publications use:
## 
##   Simon Garnier, Noam Ross, Robert Rudis, Antônio P. Camargo, Marco
##   Sciaini, and Cédric Scherer (2024). viridis(Lite) -
##   Colorblind-Friendly Color Maps for R. viridis package version 0.6.5.
## 
## A BibTeX entry for LaTeX users is
## 
##   @Manual{,
##     title = {{viridis(Lite)} - Colorblind-Friendly Color Maps for R},
##     author = {{Garnier} and {Simon} and {Ross} and {Noam} and {Rudis} and {Robert} and {Camargo} and Antônio Pedro and {Sciaini} and {Marco} and {Scherer} and {Cédric}},
##     year = {2024},
##     note = {viridis package version 0.6.5},
##     url = {https://sjmgarnier.github.io/viridis/},
##     doi = {10.5281/zenodo.4679423},
##   }
```

```
## To cite package 'Hmisc' in publications use:
## 
##   Harrell Jr F (2024). _Hmisc: Harrell Miscellaneous_. R package
##   version 5.1-3, <https://CRAN.R-project.org/package=Hmisc>.
## 
## A BibTeX entry for LaTeX users is
## 
##   @Manual{,
##     title = {Hmisc: Harrell Miscellaneous},
##     author = {Frank E {Harrell Jr}},
##     year = {2024},
##     note = {R package version 5.1-3},
##     url = {https://CRAN.R-project.org/package=Hmisc},
##   }
```

```
## To cite package 'ggthemes' in publications use:
## 
##   Arnold J (2024). _ggthemes: Extra Themes, Scales and Geoms for
##   'ggplot2'_. R package version 5.1.0,
##   <https://CRAN.R-project.org/package=ggthemes>.
## 
## A BibTeX entry for LaTeX users is
## 
##   @Manual{,
##     title = {ggthemes: Extra Themes, Scales and Geoms for 'ggplot2'},
##     author = {Jeffrey B. Arnold},
##     year = {2024},
##     note = {R package version 5.1.0},
##     url = {https://CRAN.R-project.org/package=ggthemes},
##   }
```

```
## To cite package 'AICcmodavg' in publications use:
## 
##   Mazerolle MJ (2023). _AICcmodavg: Model selection and multimodel
##   inference based on (Q)AIC(c)_. R package version 2.3.3,
##   <https://cran.r-project.org/package=AICcmodavg>.
## 
## A BibTeX entry for LaTeX users is
## 
##   @Manual{AICcmodavg-package,
##     title = {AICcmodavg: Model selection and multimodel inference based on (Q)AIC(c)},
##     author = {Marc J. Mazerolle},
##     year = {2023},
##     note = {R package version 2.3.3},
##     url = {https://cran.r-project.org/package=AICcmodavg},
##     bibtex = {TRUE},
##   }
```

```
## To cite package 'ggpmisc' in publications use:
## 
##   Aphalo P (2024). _ggpmisc: Miscellaneous Extensions to 'ggplot2'_. R
##   package version 0.6.0, <https://CRAN.R-project.org/package=ggpmisc>.
## 
## A BibTeX entry for LaTeX users is
## 
##   @Manual{,
##     title = {ggpmisc: Miscellaneous Extensions to 'ggplot2'},
##     author = {Pedro J. Aphalo},
##     year = {2024},
##     note = {R package version 0.6.0},
##     url = {https://CRAN.R-project.org/package=ggpmisc},
##   }
```

```
## To cite package 'knitr' in publications use:
## 
##   Xie Y (2024). _knitr: A General-Purpose Package for Dynamic Report
##   Generation in R_. R package version 1.48, <https://yihui.org/knitr/>.
## 
##   Yihui Xie (2015) Dynamic Documents with R and knitr. 2nd edition.
##   Chapman and Hall/CRC. ISBN 978-1498716963
## 
##   Yihui Xie (2014) knitr: A Comprehensive Tool for Reproducible
##   Research in R. In Victoria Stodden, Friedrich Leisch and Roger D.
##   Peng, editors, Implementing Reproducible Computational Research.
##   Chapman and Hall/CRC. ISBN 978-1466561595
## 
## To see these entries in BibTeX format, use 'print(<citation>,
## bibtex=TRUE)', 'toBibtex(.)', or set
## 'options(citation.bibtex.max=999)'.
```

```
## To cite package 'chunkhooks' in publications use:
## 
##   Yasumoto A (2020). _chunkhooks: Chunk Hooks for 'R Markdown'_. R
##   package version 0.0.1,
##   <https://CRAN.R-project.org/package=chunkhooks>.
## 
## A BibTeX entry for LaTeX users is
## 
##   @Manual{,
##     title = {chunkhooks: Chunk Hooks for 'R Markdown'},
##     author = {Atsushi Yasumoto},
##     year = {2020},
##     note = {R package version 0.0.1},
##     url = {https://CRAN.R-project.org/package=chunkhooks},
##   }
```

```
## To cite package 'kableExtra' in publications use:
## 
##   Zhu H (2024). _kableExtra: Construct Complex Table with 'kable' and
##   Pipe Syntax_. R package version 1.4.0,
##   <https://CRAN.R-project.org/package=kableExtra>.
## 
## A BibTeX entry for LaTeX users is
## 
##   @Manual{,
##     title = {kableExtra: Construct Complex Table with 'kable' and Pipe Syntax},
##     author = {Hao Zhu},
##     year = {2024},
##     note = {R package version 1.4.0},
##     url = {https://CRAN.R-project.org/package=kableExtra},
##   }
```

```
## To cite package 'wesanderson' in publications use:
## 
##   Ram K, Wickham H (2023). _wesanderson: A Wes Anderson Palette
##   Generator_. R package version 0.3.7,
##   <https://CRAN.R-project.org/package=wesanderson>.
## 
## A BibTeX entry for LaTeX users is
## 
##   @Manual{,
##     title = {wesanderson: A Wes Anderson Palette Generator},
##     author = {Karthik Ram and Hadley Wickham},
##     year = {2023},
##     note = {R package version 0.3.7},
##     url = {https://CRAN.R-project.org/package=wesanderson},
##   }
```

```
## To cite package 'tidyverse' in publications use:
## 
##   Wickham H, Averick M, Bryan J, Chang W, McGowan LD, François R,
##   Grolemund G, Hayes A, Henry L, Hester J, Kuhn M, Pedersen TL, Miller
##   E, Bache SM, Müller K, Ooms J, Robinson D, Seidel DP, Spinu V,
##   Takahashi K, Vaughan D, Wilke C, Woo K, Yutani H (2019). "Welcome to
##   the tidyverse." _Journal of Open Source Software_, *4*(43), 1686.
##   doi:10.21105/joss.01686 <https://doi.org/10.21105/joss.01686>.
## 
## A BibTeX entry for LaTeX users is
## 
##   @Article{,
##     title = {Welcome to the {tidyverse}},
##     author = {Hadley Wickham and Mara Averick and Jennifer Bryan and Winston Chang and Lucy D'Agostino McGowan and Romain François and Garrett Grolemund and Alex Hayes and Lionel Henry and Jim Hester and Max Kuhn and Thomas Lin Pedersen and Evan Miller and Stephan Milton Bache and Kirill Müller and Jeroen Ooms and David Robinson and Dana Paige Seidel and Vitalie Spinu and Kohske Takahashi and Davis Vaughan and Claus Wilke and Kara Woo and Hiroaki Yutani},
##     year = {2019},
##     journal = {Journal of Open Source Software},
##     volume = {4},
##     number = {43},
##     pages = {1686},
##     doi = {10.21105/joss.01686},
##   }
```
