## Supplementary information for "Effects of copepod chemical cues on intra- and extracellular toxins in two species of *Dinophysis*"

b: IFREMER, PHYTOX, PHYSALG, F-44000 Nantes, France

c: IFREMER, PHYTOX, METALG, F-44000 Nantes, France

d: Functional ecology section - Department of Biology, Lund University, Lund, Sweden

Table S1: Composition of individual copepodamide congeners in the purified extract used for the copepodamide treatments. Congeners lacking a fatty acid group (-) are deacylated scaffolds.

| Scaffold | Fatty acid | *m/z* precursor ion | | Concentration (µM) |
| --- | --- | --- | --- | --- |
| Copepodamides (*m/z* product ion: 430) | - | 448.3 | 0.030 | |
|  | 14:0 (Myritic) | 658.5 | 0.522 | |
|  | 16:0 (Palmitic) | 686.5 | 1.561 | |
|  | 18:4 (Stearidonic) | 706.5 | 2.832 | |
|  | 20:5 (Eicosapentaenoic) | 732.5 | 5.007 | |
|  | 22:6 (Cervonic) | 758.6 | 18.41 | |
| Dihydro-copepodamides (*m/z* product ion: 432) | - | 450.3 | 0.002 | |
|  | 18:4 (Stearidonic) | 708.5 | 0.236 | |
|  | 20:5 (Eicosapentaenoic) | 734.5 | 0.330 | |
|  | 22:6 (Cervonic) | 760.5 | 0.961 | |
|  |  |  | **Total: 29.9 µM** | |

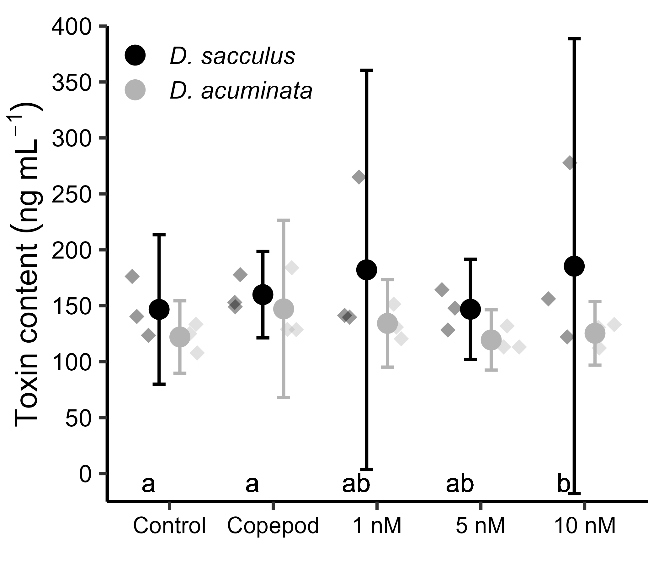

Fig. S1: Total diarrhetic toxin content normalised by sample volume, but with means and 95% confidence intervals **not** adjusted for growth rate, for *Dinophysis sacculus* (black) and *Dinophysis acuminata* (grey) after 68.5 hours of exposure to 1-10 nM concentrations of copepodamides, a living *Acartia* sp. copepod, or a solvent only control. Transparent diamonds show individual toxin content in each replicate, opaque circles are average toxin content of n = 3 replicates and error bars denote their 95% confidence intervals. Letters denote statistically homogeneous subgroups for *D sacculus.* There were no significantly different subgroups in *D acuminata*. Figure corresponds to Fig. 3 in the paper.

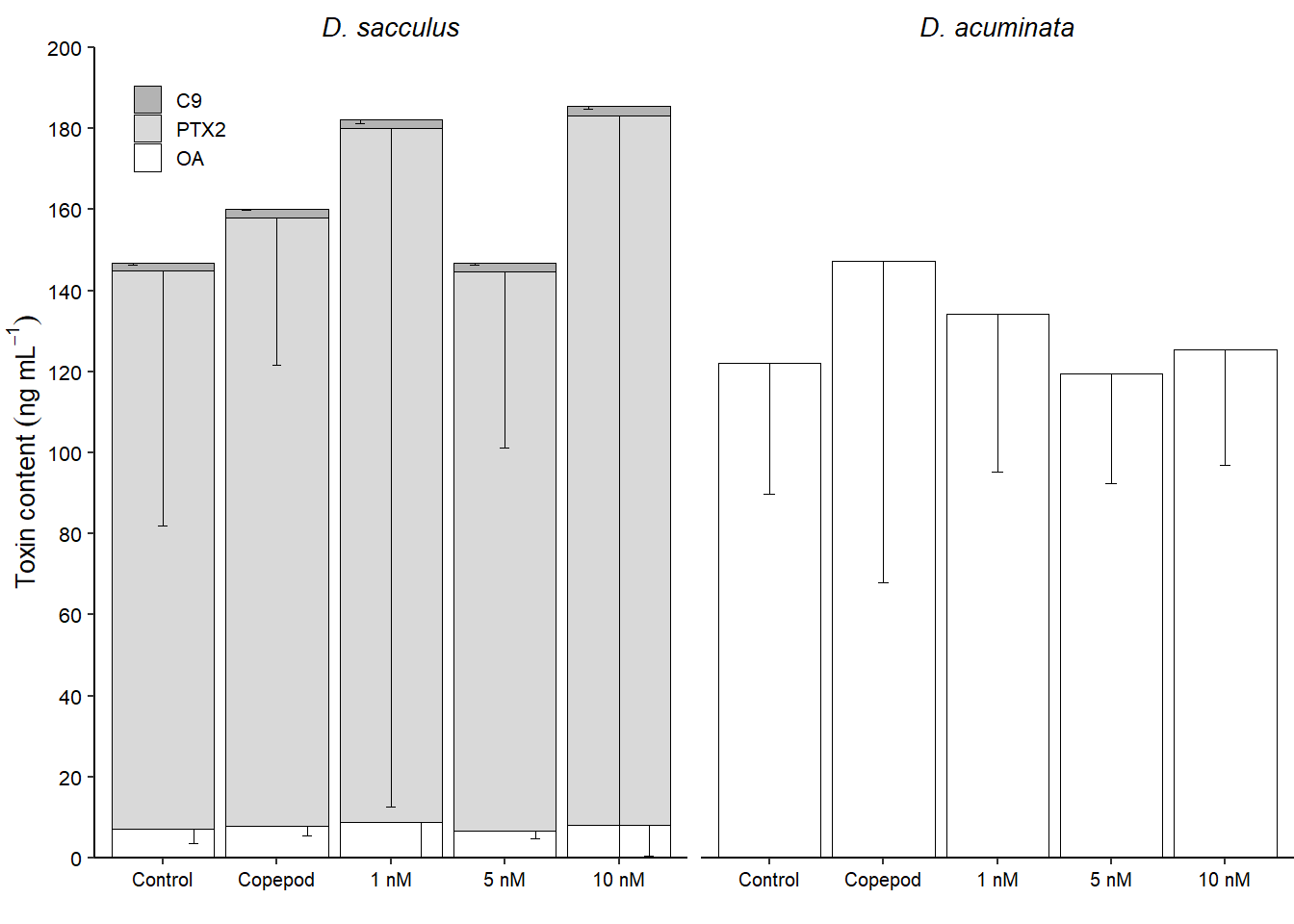

Fig. S2: Composition of *Dinophysis*-produced shellfish toxins (white = OA (Okadaic acid), light grey = PTX-2 (Pectenotoxin-2) and dark grey = C9 (OA-C9-diol ester) for *Dinophysis sacculus* (Left) and *D. acuminata* (Right) after 68.5 hours of exposure to 1-10 nM concentrations of copepodamides, a living *Acartia* sp. copepod or control conditions without copepods or copepod cues. Bars are the mean values of three replicates (n = 3) and error bars denote the 95% confidence intervals of the means. Note that only the lower confidence intervals are shown.

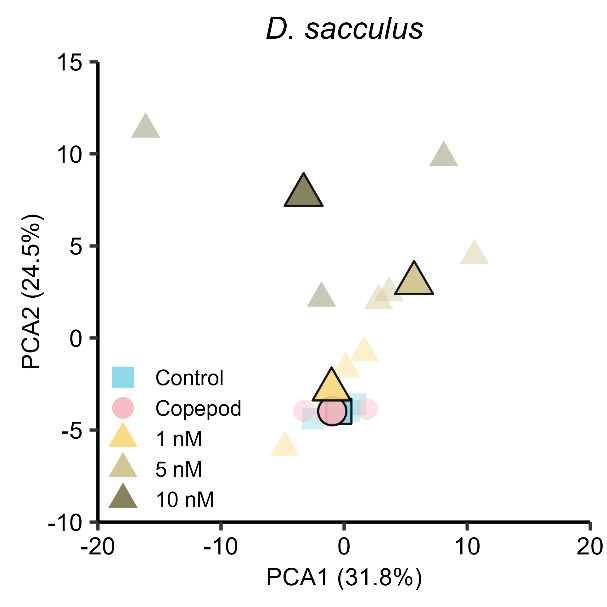

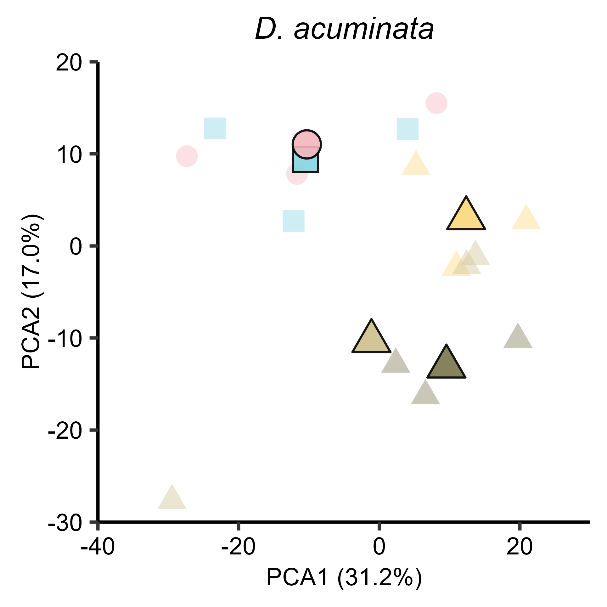

Fig. S3: PCA score plot from LC-HRMS derived metabolomic profiles for *D. sacculus* (left, -MS data) and *D. acuminata* (right, +MS data) after 68.5 hours of exposure to three concentrations of chemical cues purified from copepods (copepodamides, triangles), a living *Acartia* sp. copepod (circles), or control conditions (squares) without copepods or copepodamides. Transparent symbols without borders denote individual replicates and opaque symbols with black border denote the centroids (averages) for each group.
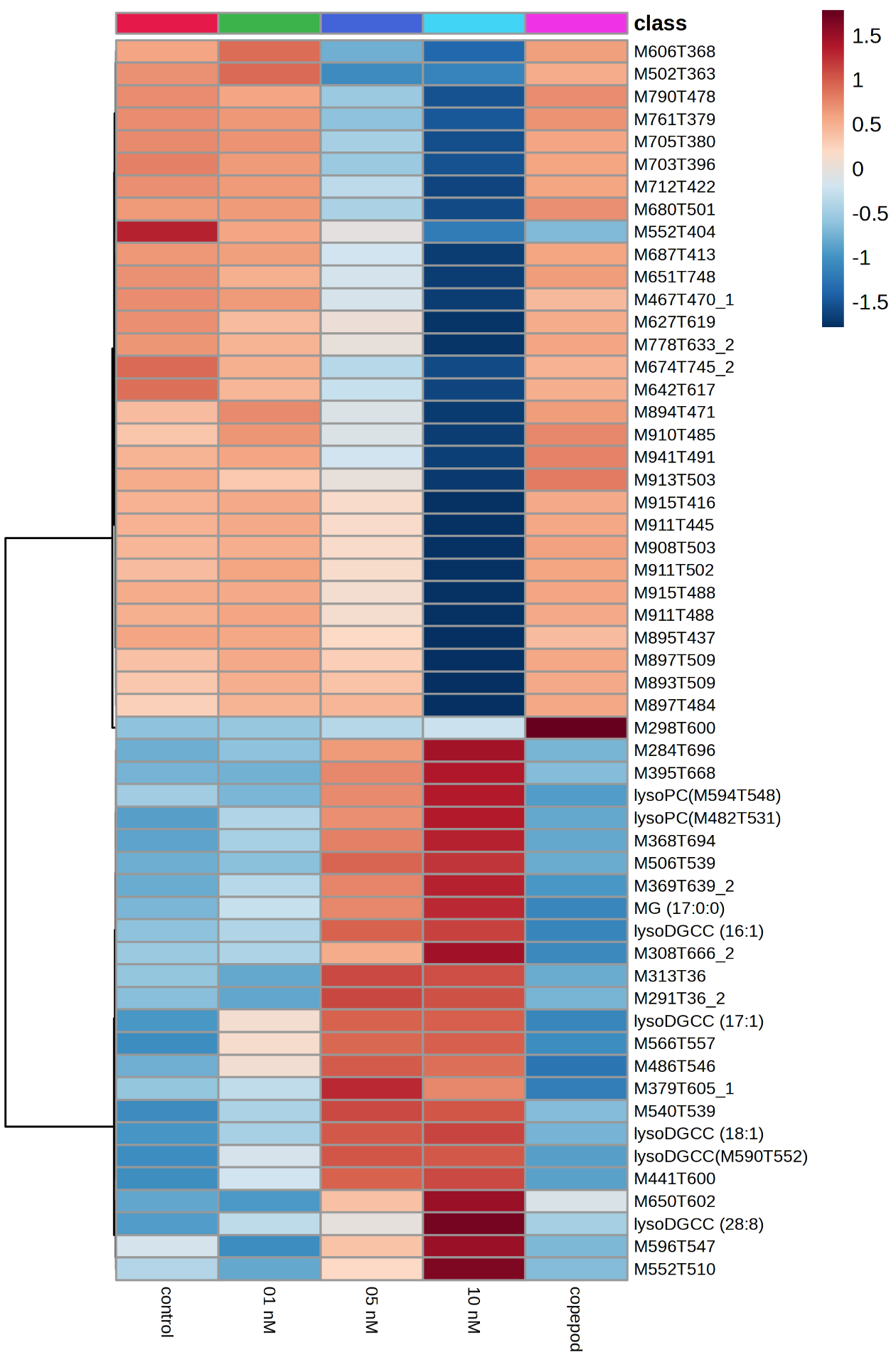

Fig. S4: Heatmap of significantly affected features of *D. sacculus* (+MS data) after 68.5 hours of exposure to three concentrations of copepodamides, a living *Acartia* sp. copepod, or control conditions. Data represented are mean normalized areas of triplicates. Putatively annotated features are specified (e.g., lysoDGCC (16:1)), otherwise features are defined as M:*m/z* of the parent ion and T:retention time in seconds. The color coding corresponded to features with a relatively lower (blue) and higher (red) level of expression for a given feature (e.g. the feature M298T600 was over-expressed only in the copepod treatment). DGCC: diacylglycerylcarboxyhydroxymethylcholine, MG: monoacylglycerol; PC: phosphatidylcholine.

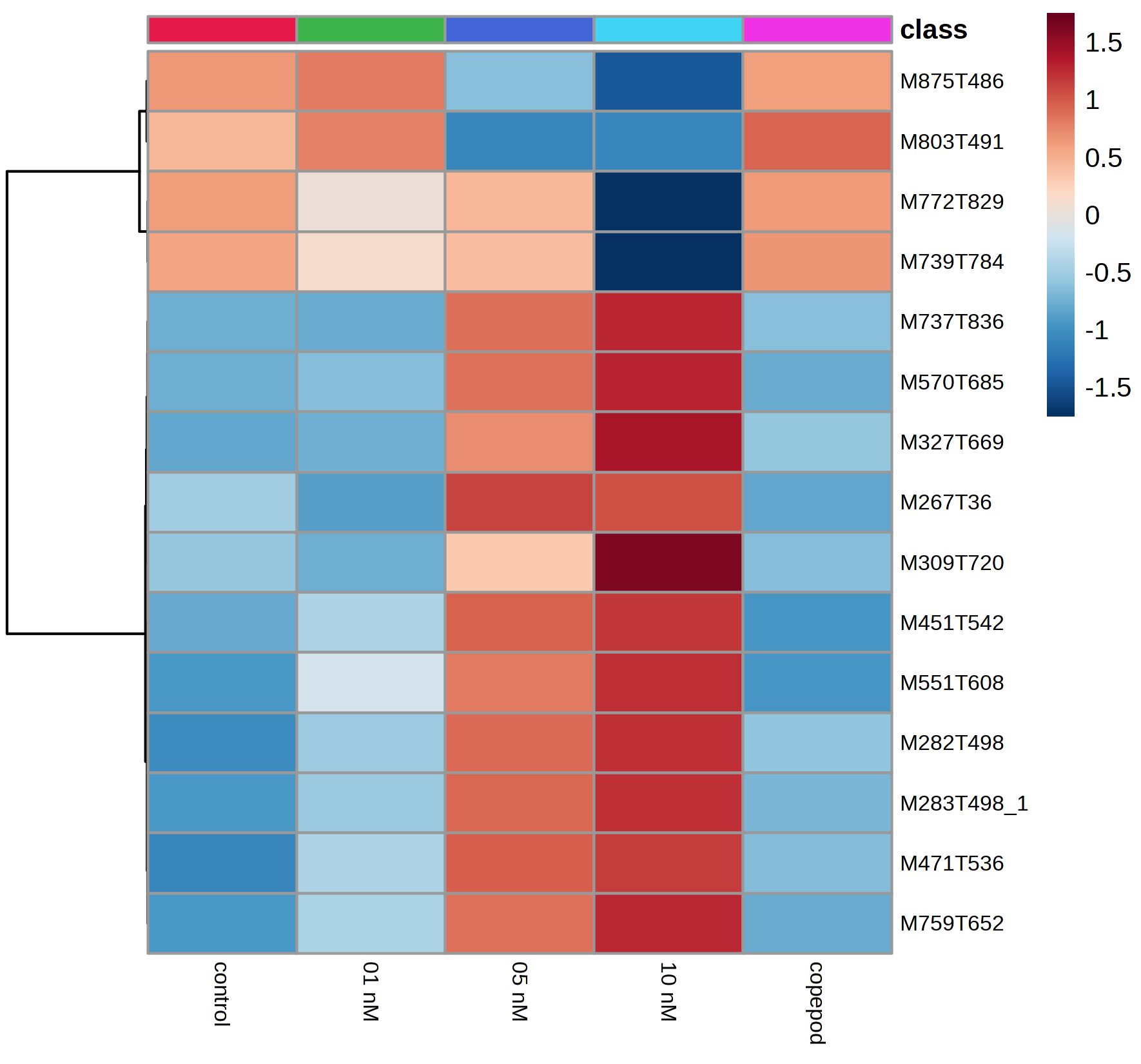

Fig. S5: Heatmap of significantly affected features of *D. sacculus* (-MS data) after 68.5 hours of exposure to three concentrations of copepodamides, a living *Acartia* sp. copepod, or control conditions. Data represented are mean normalized areas of triplicates. Features were defined as M:*m/z* of the parent ion and T:retention time in seconds. The color coding corresponded to features with a relatively lower (blue) and higher (red) level of expression for a given feature. See Fig. S3 for further details.

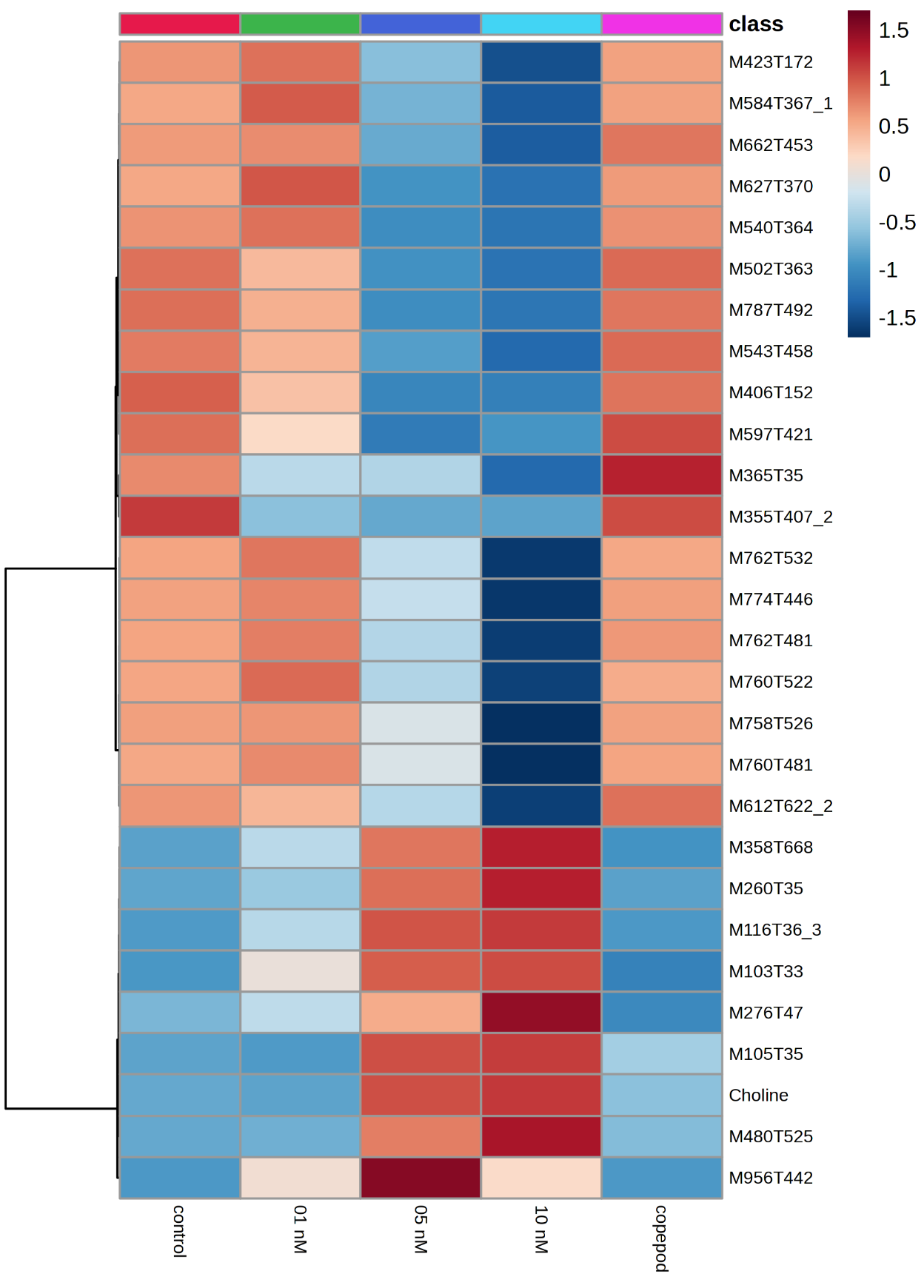

Fig. S6: Heatmap of significantly affected features of *D. acuminata* (+MS data) after 68.5 hours of exposure to three concentrations of copepodamides, a living *Acartia* sp. copepod, or control conditions. Data represented are mean normalized areas of triplicates. Putatively annotated features are specified, otherwise features are defined as M:*m/z* of the parent ion and T:retention time in seconds. The color coding corresponded to features with a relatively lower (blue) and higher (red) level of expression for a given feature. See Fig. S3 for further details.

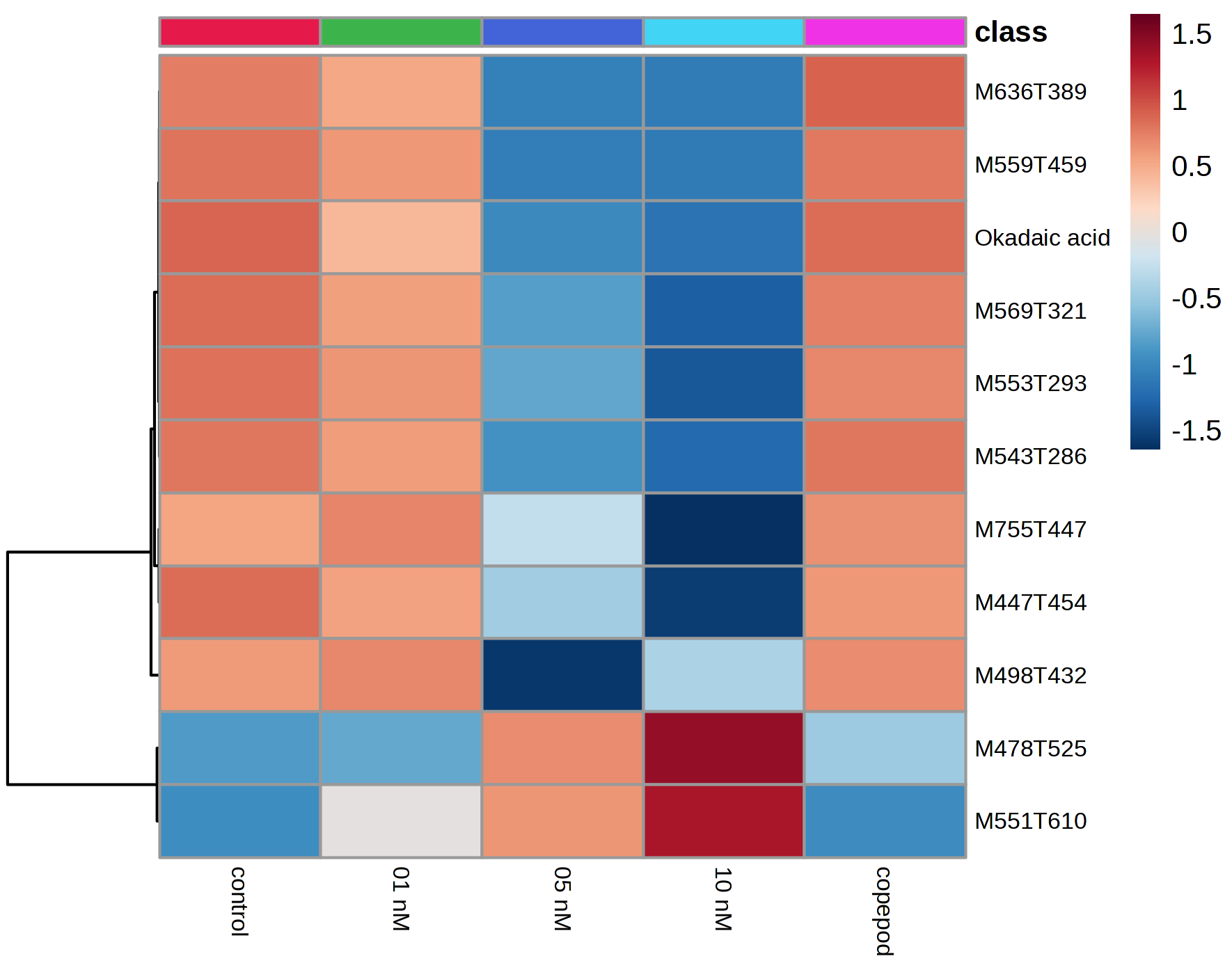
 Fig. S7: Heatmap of significantly affected features of *D. acuminata* (-MS data) after 68.5 hours of exposure to three concentrations of copepodamides, a living *Acartia* sp. copepod, or control conditions. Data represented are mean normalized areas of triplicates. Putatively annotated features are specified, otherwise features are defined as M:*m/z* of the parent ion and T:retention time in seconds. The color coding corresponded to features with a relatively lower (blue) and higher (red) level of expression for a given feature. See Fig. S3 for further details.
